# Microfluidics-enabled fluorescence-activated cell sorting of single pathogen-specific antibody secreting cells for the rapid discovery of monoclonal antibodies

**DOI:** 10.1101/2023.01.10.523494

**Authors:** Katrin Fischer, Aleksei Lulla, Tsz Y So, Pehuén Pereyra-Gerber, Matthew I. J. Raybould, Timo N. Kohler, Tomasz S. Kaminski, Juan Carlos Yam-Puc, Robert Hughes, Florian Leiß-Maier, Paul Brear, Nicholas J. Matheson, Charlotte M. Deane, Marko Hyvönen, James E. D. Thaventhiran, Florian Hollfelder

## Abstract

Monoclonal antibodies are increasingly used to prevent and treat viral infections, playing a pivotal role in pandemic response efforts. Antibody secreting cells (ASCs, plasma cells and plasmablasts) are an excellent source of high-affinity antibodies with therapeutic potential. Current methodologies to study antigen-specific ASCs either have low throughput, require expensive and labour-intensive screening or are technically demanding and therefore not accessible to the wider research community. Here, we present a straightforward technology for the rapid discovery of monoclonal antibodies from ASCs: we combine microfluidic encapsulation of single cells into an antibody capture hydrogel with antigen bait sorting by conventional flow cytometry. With our technology, we screened millions of mouse and human ASCs and obtained anti-SARS-CoV-2 monoclonal antibodies with high affinity (pM) and neutralising capacity (<100 ng/mL) in two weeks with a high hit rate (>85%). By facilitating access into the underexplored ASC compartment, we enable fast and efficient antibody discovery as well as immunological studies into the generation of protective antibodies.

## Introduction

Antibodies produced by our immune system in response to infection or vaccination are invaluable tools in therapy, diagnostics, and research. Human immune repertoires are an excellent source for the generation of potent antibody therapeutics and have been successfully used in the fight against viral infections such as COVID-19^1,2^.

Human-derived antibodies have advantages over animal-derived or *in vitro* selected binders: they are human-compatible without optimisation and are thought to have less off-target binding, a more favourable safety profile, and better overall developability^3^. Until now, human-derived therapeutic monoclonal antibodies against various infectious diseases have been primarily sourced from memory B cells^1,4–6^. These cells can be readily selected in a high-throughput manner with fluorescently labelled antigen baits by fluorescence-activated cell sorting (FACS), because they retain the membrane-bound version of their immunoglobulins on the cell surface^6^. Single sorted cells are then sequenced to obtain the antibody genes for recombinant expression.

In contrast, antibody secreting cells (ASCs, plasma cells and plasmablasts) have been undervalued as a rich source of highly specific antibodies because of technological limitations and their limited accessibility in peripheral blood^3^. Antibodies isolated from ASCs are thought to have on average higher affinity than those derived from memory B cells^7–9^. Additionally, ASCs are responsible for the active humoral immune response, i.e. they secrete the circulating antibodies that confer protection against invading pathogens^10^. Interrogating this compartment therefore brings us closer to understanding the secreted antibody repertoire and can help bridge the gap between proteomic profiling of plasma antibodies and bulk sequencing of the B cell repertoire^11^.

However, studying the specificity of single ASCs is difficult because they secrete their antibodies and express few or no immunoglobulins on the surface^12^. For this reason, ASCs cannot be interrogated by conventional FACS in an antigen-specific manner with high throughput. Traditionally, the ASC compartment has therefore been interrogated by either unbiased plasmablast sorting^13,14^, which does not give information about the antigen specificity of single cells or enzyme-linked immunospot (ELISpot) assays, which enumerate the frequency of antigen-specific cells, but are not able to recover their genotype^15^.

Since the percentage of suitable antigen binders within the plasmablast population can be low, unbiased sorting can require significant time and resource investment to screen antibodies from all cells for validation of binding. Recently, approaches based on compartmentalisation of single cells in water-in-oil emulsion droplets have been applied in antibody discovery^16–20^ and sequencing^21–23^. Alternatively, antibody discovery from ASCs can be performed with commercial optofluidic systems^24^. However, functional assays in droplets are technically demanding and standalone machines are prohibitively expensive and both approaches are therefore not accessible to the wider research community.

We address the limitations of current ASC screening methods by enabling the high-throughput interrogation of antigen-specific ASCs by conventional FACS. In our simple and robust workflow, we first use droplet microfluidics to encapsulate single cells into an antibody capture hydrogel, creating a stable capture matrix around the cell that enables the concentration of secreted antibodies and simple addition and removal of detection reagents. We then use the multiplexed detection and high throughput sorting capabilities of FACS to isolate antigen-specific ASCs for single cell sequencing and recombinant antibody expression.

We demonstrate the utility of this approach by screening millions of primary immune cells to isolate monoclonal antibodies against SARS-CoV-2 from mouse and human ASCs. Tapping into these repertoires allowed us to harvest a diverse pool of sequences of secreted antibodies. Of a representative subset of human antibodies, 95% bound the respective antigen, many with sub-nanomolar affinities and high neutralising capacities (<100 ng/mL), highlighting the benefits of interrogating the locus of the active humoral response. Importantly, our low-cost technology can generate pathogen-specific antibodies within two weeks, both democratising and fast-tracking the development of antibody drug candidates.

## Results

### Development of a single-cell antibody capture system and screening workflow

Identification of single antigen-specific ASCs presents a considerable challenge: target cells secreting antibodies with desirable properties have to be selected from a vast pool of cells producing non-specific binders whilst maintaining the link between the phenotype of the secreted antibody and the cell that encodes its sequence (genotype).

We have addressed this challenge by compartmentalising single ASCs into an antibody capture hydrogel by automated droplet microfluidics (at a rate of up to 10^7^ cells/hour), followed by the selection of the secreted antibody specificity with fluorescently labelled antigens by FACS at a similar speed (**Fig. 1a**). This combination of microfluidics and FACS enables the high throughput that is crucial for the success of antibody discovery campaigns.

**Fig. 1.**
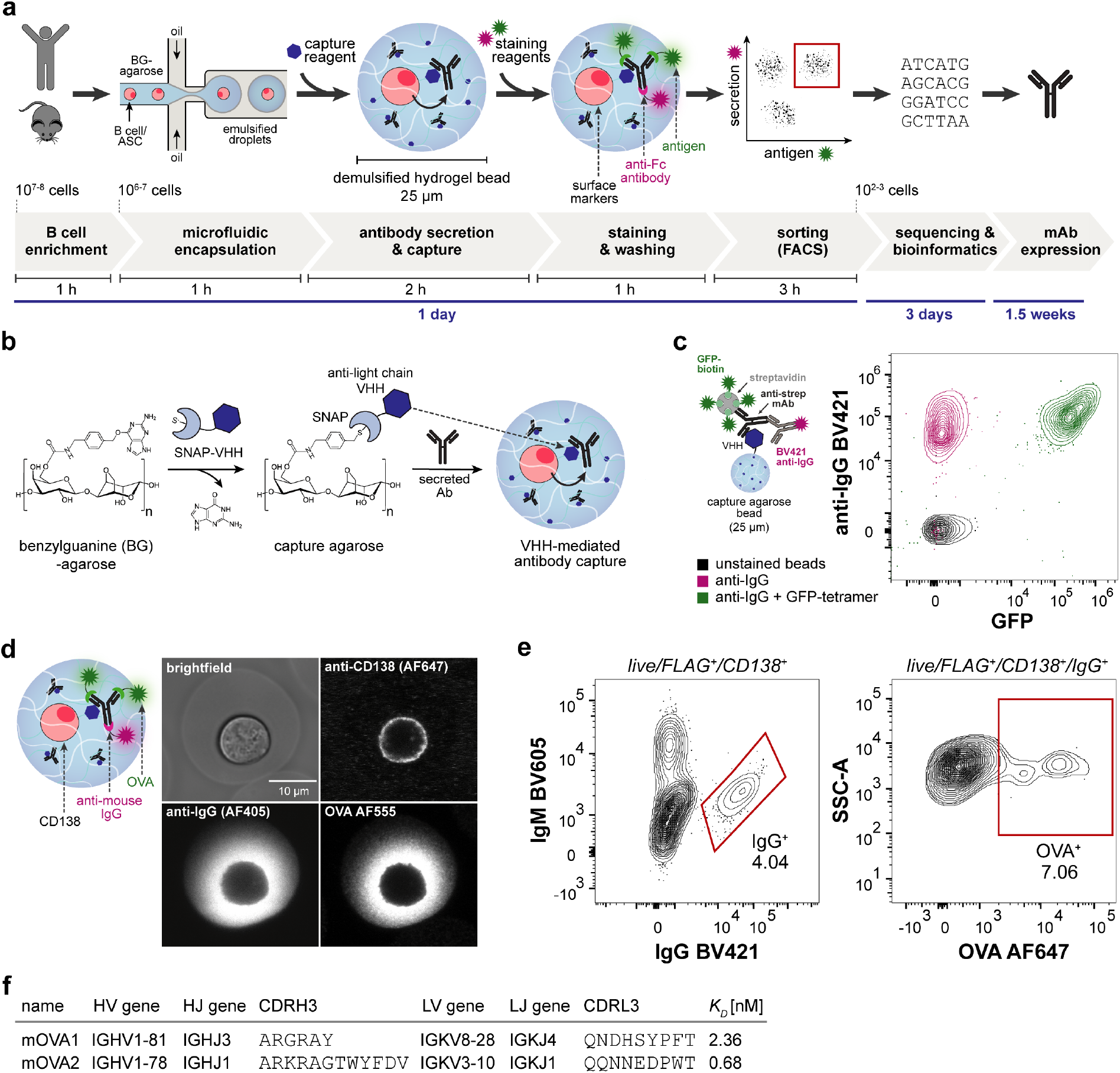
Workflow for high-throughput functional analysis of antibodies secreted by single cells. (a) Overview of the workflow: B cells (or enriched ASCs) are isolated from mouse (bone marrow or spleen) or human peripheral blood mononuclear cells (PBMCs). Cells are mixed with liquid benzylguanine (BG)-agarose at 37 °C and encapsulated into picolitre water-in-oil emulsion droplets using a flow-focusing junction. Droplets are collected on ice for agarose gelation and are then demulsified, creating stable hydrogel beads around each cell. The BG-agarose is converted into an antibody capture matrix by addition of recombinant capture reagents that are fused to the SNAP-tag, an enzyme that reacts with BG-moieties. During incubation, antibodies secreted by a single cell are captured in the hydrogel surrounding the cell. Cells that have secreted antigen-specific antibodies are identified with fluorescently labelled detection reagents (antigen, secondary antibodies, antibodies against cell surface markers), sorted using flow cytometry, and sequenced. The workflow was validated by obtaining antibody sequences within four days; recombinant antibodies for testing can be generated in two weeks. (b) Agarose-based antibody capture matrix. Agarose is chemically modified to contain benzylguanine (BG) moieties which react covalently with the SNAP-tag. Single-domain antibodies (VHHs) against the constant region of antibody light chains are expressed as SNAP-tag fusions and immobilised in the BG-agarose hydrogel, creating a capture matrix for secreted antibodies. (c) Antibody capture by BG-agarose hydrogel beads functionalised with VHH-SNAP. Antibody capture (mouse anti-streptavidin) and antigen binding (streptavidin-GFP) were analysed by flow cytometry. The plot shows at least 220 events per condition at 5% contour level. (d) Antibody secretion by single OVA-specific mouse bone marrow plasma cells. Confocal microscopy image shows a single cell encapsulated into VHH-functionalised BG-agarose stained with fluorescently labelled OVA (AF555), anti-CD138 antibodies (AF647) and anti-mouse IgG antibodies (AF405). (e) Sorting of OVA-specific mouse bone marrow plasma cells. Hydrogel beads containing plasma cells that secreted OVA-specific IgG were sorted by FACS (gated as live/FLAG^+^/CD138^+^/IgM^-^/IgG^+^). The plots show 10,168 (CD138^+^) and 411 (IgG^+^) events at 2% contour level. (f) Characteristics of mouse anti-OVA antibodies: variable domain genes (V and J), third complementarity-determining region amino acid sequences (CDR3s) and equilibrium dissociation constants (K_D_).

Specifically, B cells are mixed with liquid benzylguanine (BG)-agarose (at 37°C) and encapsulated into monodisperse water-in-oil emulsion droplets of 25 µm diameter at kHz rates. The droplets are collected on ice (to solidify the agarose) and demulsified, creating a stable agarose microcompartment around each cell. BG-modification creates covalent attachment sites for the SNAP-tag, a 20 kDa engineered human O6-alkylguanine-DNA alkyltransferase which reacts specifically, quickly, and irreversibly with different BG-containing substrates^25^. We synthesised BG-agarose by chemical modification of commercial low melting point agarose in a simple, two-step procedure (**Extended Data Fig. 1a**). To create an antibody capture matrix, we functionalised the BG-sites with recombinant SNAP-tag fusion proteins of single domain antibodies (VHHs) that bind to the constant region of antibody light chains (**Fig. 1b**)^26,27^. Through this covalent functionalisation with antibody capture reagents, antibodies secreted by encapsulated cells are immobilised around the cell, physically linking the secreted antibody and the cell of origin. With two different VHHs, recognising either κ or λ light chains, any secreted antibody from any ASC can be captured efficiently. Importantly, this capture approach is highly modular, as any protein fused to the SNAP-tag can be immobilised in the agarose matrix. In the hydrogel, antibodies and cells are accessible for staining with fluorescently labelled detection reagents and antigens due to the large pore size of the agarose (>100 nm^28,29^) while unbound reagents can be removed by washing. Additionally, the small size (25 µm diameter) and monodispersity of the hydrogel beads enables sorting by conventional FACS devices. Hydrogel beads can therefore essentially be stained, washed, and sorted like single cells but provide additional information about the secreted antibody (secreted amount, isotype, antigen specificity). Lastly, single sorted cells in hydrogel beads are sequenced to enable recombinant expression and characterisation of antigen-specific antibodies. Importantly, FACS allows the detection of several phenotypic parameters or different antigens in one experiment and the correlation of the sequence of single ASCs with phenotypic antibody information through index sorting.

We estimated the capture capacity of the BG-agarose by reacting BG-agarose beads (25 µm diameter, 1.5% w/v agarose) with different quantities of GFP-SNAP, followed by the analysis of the GFP-fluorescence of the beads by flow cytometry (**Extended Data Fig. 1b**). We observed saturation of the GFP signal at over 10^9^ immobilised GFP molecules per bead. In the literature, average secretion rates of single ASCs range from 10^3^ – 10^5^ antibodies per second^30,31^, indicating that a completely functionalised capture matrix would not become saturated even after several hours of antibody secretion. Notably, in our system, the number of antibody capture sites is a function of bead size and amount of VHH-SNAP added and is therefore well-defined and controllable. In contrast to capture on the cell surface^32,33^, antibody capture capacity does not depend on cell-intrinsic properties such as cell-surface molecules and should therefore be relatively uniform across the ASC population.

Next, we showed that light-chain mediated capture enables interrogation of both antigen binding and detection of immobilised antibodies by flow cytometry (**Fig. 1c**). We captured a commercial anti-streptavidin IgG antibody in VHH-functionalised BG-agarose beads and stained the beads with both streptavidin-biotin-GFP complexes and anti-IgG antibodies. Analysis by flow cytometry confirmed that captured antibodies can simultaneously bind to the functionalised agarose, the antigen, and the detection antibodies.

### Capture of antibodies secreted by single murine bone marrow plasma cells

With a capture system for secreted proteins at hand, we established an antibody discovery workflow with primary immune cells. We immunized mice with hen egg ovalbumin (OVA) to obtain OVA-specific B cells and isolated bone marrow plasma cells (CD138^+^) by positive magnetic enrichment (**Extended Data Fig. 2a**). Enriched plasma cells were then encapsulated into BG-agarose functionalised with anti-mouse κ VHH-SNAP.

To visualize secretion and capture of OVA-specific antibodies, we stained cells compartmentalised in hydrogels with fluorescently labelled antibodies against the plasma cell marker CD138, secondary antibodies against IgG (to visualise secreted antibodies), and fluorescently labelled OVA. By confocal fluorescence microscopy, we observed co-localisation of IgG and OVA-signal in the hydrogel around the cell, indicating secretion of OVA-specific IgG (**Fig. 1d**). To obtain OVA-specific antibody sequences for recombinant expression, we encapsulated bone marrow-derived plasma cells for sorting by FACS. After incubation, hydrogel beads were stained with fluorescently labelled monomeric OVA, antibodies against B cell and ASC surface markers (B220, CD138), an anti-FLAG antibody for labelling of VHH-functionalised hydrogels, as well as anti-IgG and IgM secondary antibodies. We hypothesized that a substantial IgG signal is only generated in the presence of VHH because mouse bone marrow plasma cells do not express surface IgG^34^. Additionally, in the absence of VHH, secreted antibodies are removed during washing steps due to the low background binding and large pore size of the agarose. We detected a distinct IgG^+^ population only in samples to which VHH-SNAP was added, confirming that the antibody signal originated from secreted antibodies rather than surface Ig (**Extended Data Fig. 2b**). Plasma cells secreting OVA-specific antibodies (**Fig. 1e**, representative gating strategy in **Extended Data Fig. 2c**) were sorted into single wells of 96 well plates and antibody variable genes were obtained by RT-PCR followed by Sanger sequencing. We recombinantly expressed two antibodies and found that they both bound OVA with high affinity using biolayer interferometry (K_D_s 2.36 nM and 0.68 nM, respectively) (**Fig. 1f, Extended Data Fig. 2d**), suggesting that high affinity binders can be identified by our workflow. In addition to bone marrow-derived plasma cells, we also analysed OVA-specific spleen-derived ASCs with the workflow (**Extended Data Fig. 3**). With this setup, we can therefore screen millions of cells in a matter of hours and generate high affinity monoclonal antibodies from primary mouse immune cells.

### Direct discovery of mouse anti-SARS-CoV-2 RBD antibodies

We then assessed our ability to isolate high-affinity antibodies against a relevant pathogen antigen. We immunised mice with the receptor binding domain (RBD) of the wildtype (WT) SARS-CoV-2 spike protein, enriched plasma cells from bone marrow, and used them in workflow (**Fig. 2a**). Plasma cells secreting RBD-specific IgG antibodies were sorted with fluorescently labelled RBD-streptavidin tetramers into single wells of 96 well plates for Sanger sequencing (**Fig. 2b**, gating strategy in **Extended Data Fig. 4b**). Secreted antibodies could be identified by comparing the IgG signal of samples with and without VHH addition (**Extended Data Fig. 4a**).

**Fig. 2.**
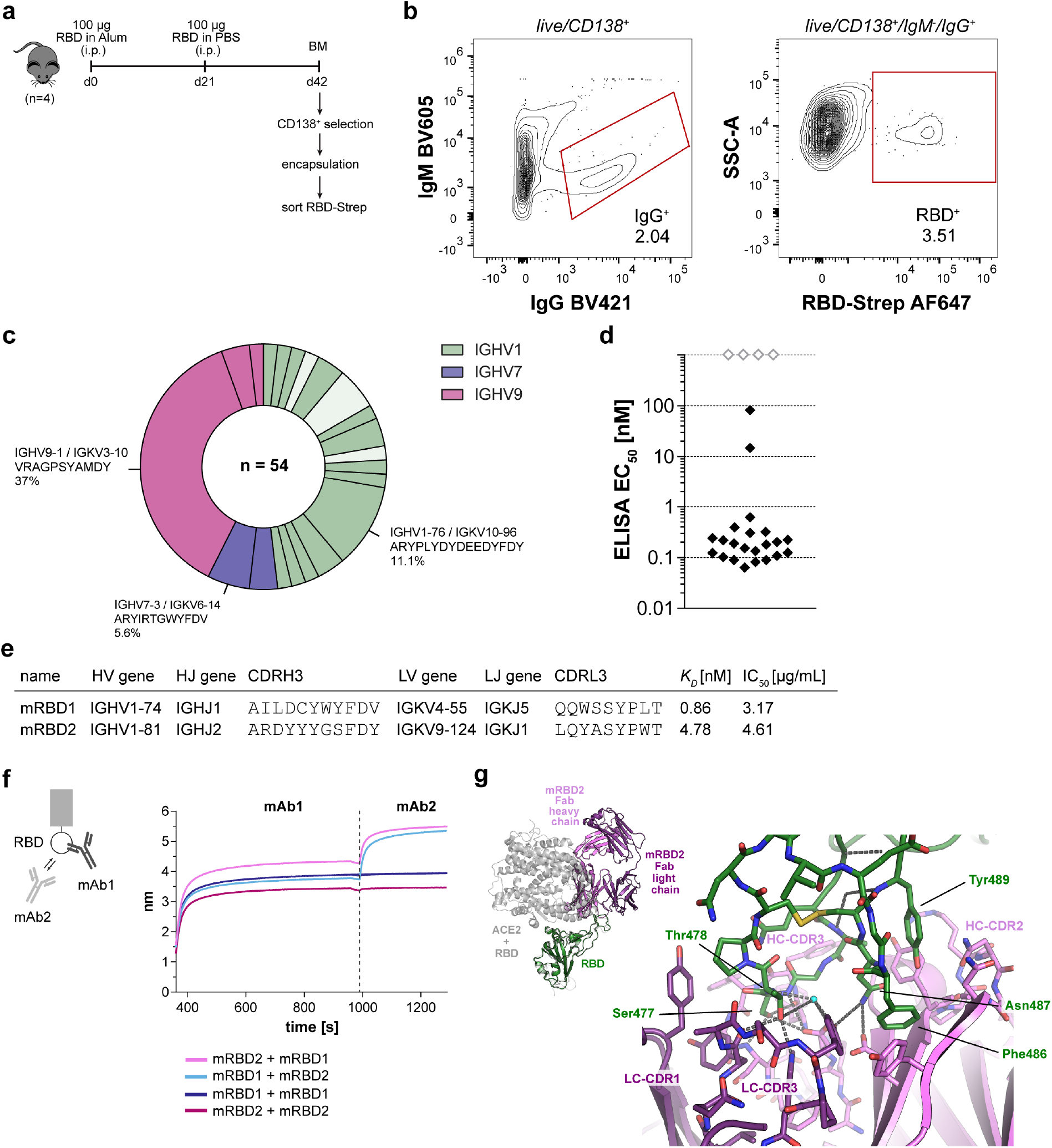
Generation of mouse anti-SARS-CoV-2 RBD antibodies. (a) Mouse immunisation and analysis scheme. Bone marrow plasma cells (CD138^+^) were magnetically enriched and then used in workflow. RBD-specific plasma cells were sorted with fluorescently labelled RBD-streptavidin tetramers. (b) Sorting of RBD-specific mouse plasma cells. Cells were gated as live/CD138^+^ and IgG-secreting RBD-specific plasma cells were sorted by FACS. The plots show 79,629 (CD138^+^) and 1,623 (IgG^+^) events at 2% contour level. (c) Overview of antibody sequences of sorted plasma cells. In total, 54 paired heavy and light chain sequences were obtained. The pie chart shows the 21 observed heavy variable (HV) and light variable (LV) gene combinations (HV-LV). HV-LV pairings are coloured by HV gene, combinations that were characterised are shown in darker shades while combinations that were not expressed are shown in a lighter shade. The three most expanded expressed HV-LV combinations are highlighted with their CDRH3 amino acid sequence and frequency. (d) Summary of anti-RBD ELISA. The plot shows the median effective concentrations (EC_50_) with an antibody concentration range of 0.0002 – 400 nM. Antibodies that did not bind RBD at 400 nM are shown at an arbitrary EC_50_ of 1,000 nM (grey diamonds). (e) Characteristics of mouse anti-SARS-CoV-2 RBD antibodies with neutralising capacity: variable domain sequences (V and J genes), third complementarity-determining region amino acid sequences (CDR3), equilibrium dissociation constants (K_D_) and half-maximal inhibitory concentrations (IC_50_) against wildtype SARS-CoV-2. (f) In-tandem epitope binning experiment with mRBD1 and mRBD2. (g) Crystal structure of mRBD2 with SARS-CoV-2 RBD. On the top left the RBD (green) in complex with mRBD2 Fab fragment (purple and pink for light and heavy chains) is superimposed with RBD complexed with ACE2 (gray, PDB:6m0j) showing how the Fab fragment overlaps significantly with ACE2. The main figure shows details of the RBD loop (green carbon atoms) binding to the CDRs of the mRBD2 Fab fragment.

We found 54 antibody sequences, belonging to 21 different heavy/light V gene pairs (**Fig. 2c**) and representing 20 distinct VH clonotypes (see **Methods**). Within this diversity, there were five highly expanded clones (≥ 3 members), including an IGHV9-1 clone (20 members, 37% of all, representative CDRH3 VRAGPSYAMDY) and an IGHV1-76 clone (7 members, 13% of all, representative CDRH3 ARYPLYDYDEEDYFDY). This observation of exceptional clonality amongst an antigen-specific plasma cell-derived population of antibodies is consistent with a recent mass-spectrometry-based proteomic study of resting and post-infection human IgG antibody repertoires^11^, and highlights that our method can offer insight into the most dominant antibodies in an organism’s active antigen response. Clonal comparisons to the Coronavirus Antibody Database (CoV-AbDab)^35^ revealed that two of our antibodies belonged to the same VH clonotype as antibodies from a separate survey^36^ into the murine lymph-node derived plasmablast response to SARS-CoV-2 spike (mRBD1 clustered with 2C02 and 2D01, mRBD10 clustered with 1A10), suggesting that immunodominance can be detected amongst ASC-derived antibodies. Compared to the surface immunoglobulin-based sorting of plasma cell precursors^36^, our study adds insight into the secreted antibody response by fully differentiated plasma cells.

To confirm SARS-CoV-2 RBD binding and perform further functional profiling, we recombinantly expressed 26 representative antibodies as full-length mouse IgG1 (**Extended Data Table 1**, full length sequences in **Supplementary Table 4**). In an indirect ELISA against RBD, we found that 22 antibodies (84.6%) bound RBD at the tested concentrations (up to 400 nM antibody) (**Fig. 2d, Extended Data Fig. 4c**), including the representatives of all five of the most expanded clones. To determine whether the RBD-specific binders were also able to neutralise SARS-CoV-2, we used luminescent reporter cells to perform live virus neutralisation assays^37–39^. Despite the high apparent affinity of our plasma cell clones, only two antibodies (mRBD1 and mRBD2 (belonging to the fifth most expanded clone)), were able to neutralise WT SARS-CoV-2 with IC_50_s of 3.17 µg/mL and 4.61 µg/mL, respectively (**Fig. 2e** and **Extended Data Fig. 4d**), highlighting that selection of an antibody in an immune response does not guarantee neutralising capacity. To further characterise mRBD1 and mRBD2, we determined their binding affinities to RBD (**Extended Data Fig. 4e**) and performed in-tandem epitope binning assays by biolayer interferometry (**Fig. 2f**). The binning assay indicated that the two antibodies do not share the same binding site on RBD as an increase in BLI signal was observed when antibodies were added sequentially to immobilised RBD.

To provide further insights into the binding mechanism of murine ASC-derived RBD-binding antibodies^36,40^ we obtained a co-crystal structure of the Fab fragment of mRBD2 with RBD. The structure shows that mRBD2 binds to a long disulphide-bonded loop (residues 474-489) on the RBD that is part of the ACE-2 binding interface (**Fig. 2g**). In the antibody, the binding site is formed of CDR2 and 3 from both light and heavy chains. Phe487 and Tyr489 of the RBD interact with CDR2 of the heavy chain and Asn487 is hydrogen bonded to both heavy and light chains. Ser477 hydrogen bonds to heavy chain CDR3 while Thr478 hydrogens bonds to light chain CDR3. The conformation of the RBD loop itself is almost identical to that seen in ACE2:RBD complex, while moving through a hinge-like motion by a few ångstrom relative to rest of the RBD (**Supplementary Figure 2**). Comparison of mRBD2 with the ACE-2 blocking murine plasmablast-derived antibody 2B04^40^ showed that their binding sites on RBD partially overlap, hinting at similar neutralisation mechanisms.

### Direct discovery of anti-SARS-CoV-2 RBD antibodies from humans

To demonstrate that our technology has utility in pandemic response efforts, we adapted our protocol to isolate antibodies directly from humans. Monoclonal antibodies derived from humans after infection or vaccination have been shown be efficacious and well-tolerated in the human body, representing a fast track to therapeutic development^3^. Furthermore, studying antigen-specific plasmablasts gives unique insights into the post-infection and vaccine responses of different individuals^41,42^.

To adjust the workflow for the capture of human antibodies, we exploited the modular setup of the capture system and exchanged the anti-mouse VHH with anti-human κ and λ VHHs^27^. Firstly, we activated normal peripheral blood mononuclear cells (PBMCs) *in vitro* to stimulate antibody secretion^43^ and were able to detect antibody secretion and capture from single cells (**Extended Data Fig. 5a** and **b**). The compatibility of the workflow with *ex vivo* activated cells highlights the versatility of the technology and expands the scope of the method to the analysis of activated memory B cells.

Next, we identified SARS-CoV-2 RBD-specific antibodies directly from human antibody secreting plasmablasts. We isolated PBMCs 7 – 9 days after the second dose of the mRNA vaccine BNT162b2 and enriched for B cells by negative selection (**Fig. 3a**). After encapsulation into BG-agarose and functionalisation with anti-human κ and λ VHH-SNAP, we observed a distinct IgG-secreting, RBD-positive ASC population (**Fig. 3b, Extended Data Fig. 6a** and **b** for gating and IgG secretion control). We sorted IgG-secreting RBD-positive cells from fresh or cryopreserved samples into single wells of 96 well plates for sequencing.

**Fig. 3.**
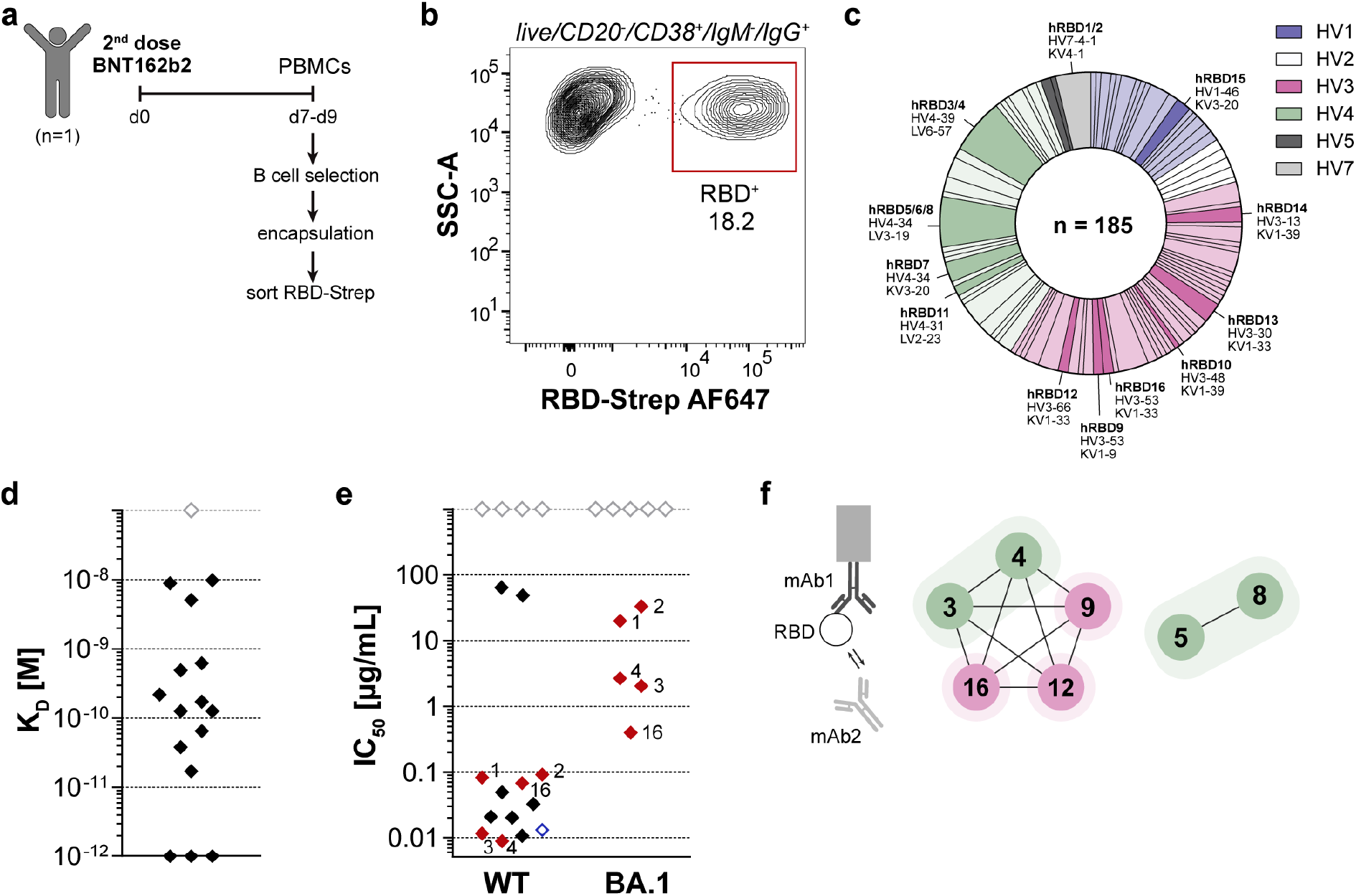
Generation of human anti-SARS-CoV-2 RBD antibodies. (a) Human anti-RBD antibody discovery. B cells were negatively selected from fresh or frozen PBMCs 7-9 days post the second BNT162b2 vaccine dose and then used in the workflow. RBD-specific ASCs were sorted with fluorescently labelled RBD-streptavidin tetramers. (b) Sorting of RBD-specific human ASCs. Representative layout for sorting of IgG^+^/RBD^+^ hydrogel beads obtained from encapsulation of fresh B cells on d9 post vaccination. Hydrogel beads were gated as live/CD20^-^/CD38^+^/IgM^-^/IgG^+^. The plot shows 1,484 events at 2% contour level. (c) Overview of sequences of human ASCs sorted with RBD. In total, 185 paired heavy and light chain sequences were obtained. The pie chart shows the observed HV-LV combination coloured by HV gene. HV-LV combinations that were expressed recombinantly are highlighted with their clone names and are shown in a darker shade. (d) Affinities (K_D_) of human anti-RBD antibodies. One antibody (grey diamond) did not show binding to RBD at the tested concentrations. (e) Neutralisation of wildtype (WT) and Omicron BA.1 SARS-CoV-2 by human anti-RBD antibodies. IC_50_, half-maximal inhibitory concentration calculated from experiments performed in duplicate. Antibodies with no quantifiable neutralising capacity at the highest concentration tested (100 μg/mL) are shown at an arbitrary IC_50_ of 1,000 μg/mL (grey diamonds). The ten antibodies with lowest IC_50_s against WT SARS-CoV-2 were also tested against Omicron BA.1 SARS-CoV-2. Antibodies neutralising both variants are highlighted in red. For comparison, neutralisation of WT SARS-CoV-2 by the REGEN-COV (Ronapreve) monoclonal antibody cocktail (casirivimab and imdevimab) was tested in the same assay (blue diamond). (f) Sandwich epitope binning experiment with pairs of neutralising human anti-RBD antibodies. In the network plot, the nodes show the antibody clones, the connections indicate pairwise blocking and the shaded areas indicate whether the antibodies belong to the same clonotype. Colours indicate the VH gene as in (c).

From just one individual, we obtained 185 plasmablast sequences, belonging to a broad spread of heavy/light V gene pairs (**Fig. 3c**). The 185 antibodies spanned 156 different VH sequences and 111 different VH clonotypes (see **Methods**). As in the mouse study, we observed a population of extremely expanded VH clonotypes. Fifteen clonotypes had an occupancy of ≥ 3 members, including an IGHV4-34 clone (10 members, 5.4% of total, representative CDRH3 ARAHLIGDCGGGSCYSGPDPSNWFDP), an IGHV4-39 clone (8 members, 4.3% of total, representative CDRH3 ARRRAGSYFKDLFDY) and an IGHV7-4-1 clone (8 members, 4.3% of total, representative CDRH3 ARVGRAAIAALDDAFDI).

Clonal clustering against the thousands of human SARS-CoV-2 response antibodies in CoV-AbDab showed that 17 (9%) of our plasmablast sequences belonged to the same VH clonotype as an antibody already characterised to be a SARS-CoV-2 RBD binder^35^. Several of these belonged to immunodominant VH clonotypes based on their identification across numerous independent studies on alternative B cell compartments from SARS-CoV-2 convalescent patients (such as IGHV3-53 with representative CDRH3 ARDLGEAGGMDV: 4 sequences in our study, and six sequences from six other studies as of February 2022^44–49^. This clustering demonstrates the presence of RBD-reactive bulk B cell and memory B cell clones amongst a post-vaccination ASC population, confirming their differentiation/expansion as part of the active immune response against SARS-CoV-2.

Additionally, we performed Fv region structural clustering of our sequences alongside CoV-AbDab using the SPACE protocol^50^, which clusters anti-coronavirus antibodies by predicted structure and can group them into sets that bind the same antigenic site. This enables an orthogonal consideration of immunodominant epitopes since antibodies that have the same topology but belong to a different VH clonotype can recognise the antigen with the same binding mode. SPACE yielded an additional 13 shared clusters to CoV-AbDab, highlighting cases where the same epitope was likely to be targeted even though the sequences did not belong to the same clonotype.

We then expressed 16 antibodies that either belonged to the most prevalent clonotypes not yet present in the CoV-AbDab (hRBD1-8, belonging to three different clonotypes) or were implicated in RBD binding based on SPACE structural clustering with antibodies found in the CoV-AbDab (hRBD 9-16, belonging to eight different clonotypes) as full length human IgG1 (**Extended Data Table 2**, full length sequences in **Supplementary Table 5**). Of these antibodies, 94% (15/16) bound RBD at the tested concentrations with affinities in the low pM to nM range (**Fig. 3d**). Of the RBD binders, 80% (12/15) were also able to neutralise WT SARS-CoV-2. Amongst these, 10 antibodies showed high neutralising capacity, with IC_50_s comparable to the REGEN-COV (Ronapreve) monoclonal antibody cocktail (range approx. 10 – 100 ng/mL) (**Fig. 3e**). As observed for many other antibodies induced by WT-SARS-CoV-2^51,52^, neutralising capacity against Omicron BA.1 SARS-CoV-2 was generally reduced. Nonetheless, it was still quantifiable for five of our antibodies (**Fig. 3e**), and one antibody (hRBD16) retained most of its neutralising capacity (IC_50_ 68 ng/mL against WT compared to 398 ng/mL against Omicron BA.1). Interestingly, hRBD16 is highly homologous to CAB-B37, a memory B cell-derived antibody that is able to neutralise a wide range of SARS-CoV-2 variants^53^, suggesting that hRBD16 might also be broadly neutralising. We used epitope binning by biolayer interferometry to further characterise seven strongly neutralising antibodies and determined that they bound to two distinct competition regions on WT RBD (hRBD3/4/9/12/16 and hRBD5/8 bound to the same regions, respectively) (**Fig. 3f**). Conversely, the differences in neutralising capacity against Omicron BA.1 (hRBD3/4/16 neutralised Omicron BA.1, while hRBD9/12 did not) indicate that although these antibodies bind to the same region on the RBD they interact with different residues.

### Selection of specific binding sites during ASC screening

To take advantage of the multiplexed readout of FACS, we then aimed to select for specific binding properties detectable during the single cell screen. As a proof of concept, we aimed to isolate antibodies that bind to the S1 region of the spike protein but do not bind to the RBD. We encapsulated B cells isolated from PBMCs seven days post the second BNT162b2 dose and stained hydrogel beads with fluorescently labelled S1- and RBD-streptavidin (**Fig. 4a**). We observed a distinct S1-positive but RBD-negative population and sorted and sequenced all antigen-specific IgG- or IgA-positive ASCs (**Fig. 4b**, secretion control and gating strategy in **Extended Data Fig. 7a** and **b**). We selected four sequences that showed the largest S1/RBD FACS signal ratio for expression as full length human IgG1 (highlighted as red squares in **Fig. 4b**, full length sequences in **Supplementary Table 5**). We were able to obtain the binding affinities of three antibodies and found that they bound S1 with affinities in the picomolar range (**Fig. 4c**). One of the antibodies (hS1-1) was able to neutralise WT SARS-CoV-2 with an IC_50_ of 102.9 ng/mL (**Fig 4c, Extended Data Fig. 7c**) but did not show neutralising capacity against the Omicron BA.1 variant (**Supplementary Figure 4**). We verified binding of all antibodies to S1 but not RBD by ELISA (**Fig. 4d**), suggesting that we successfully selected for S1 but not RBD binding in our screen. These results highlight the advantages of the multiplexed, FACS-based readout as it enables us to pre-select relevant candidates (e.g. binders to specific epitopes or multiple viral variants) and focus attention on the downstream characterisation of antibodies with specific properties rather than investing resources in the expression of many potentially unsuitable candidates.

**Fig. 4.**
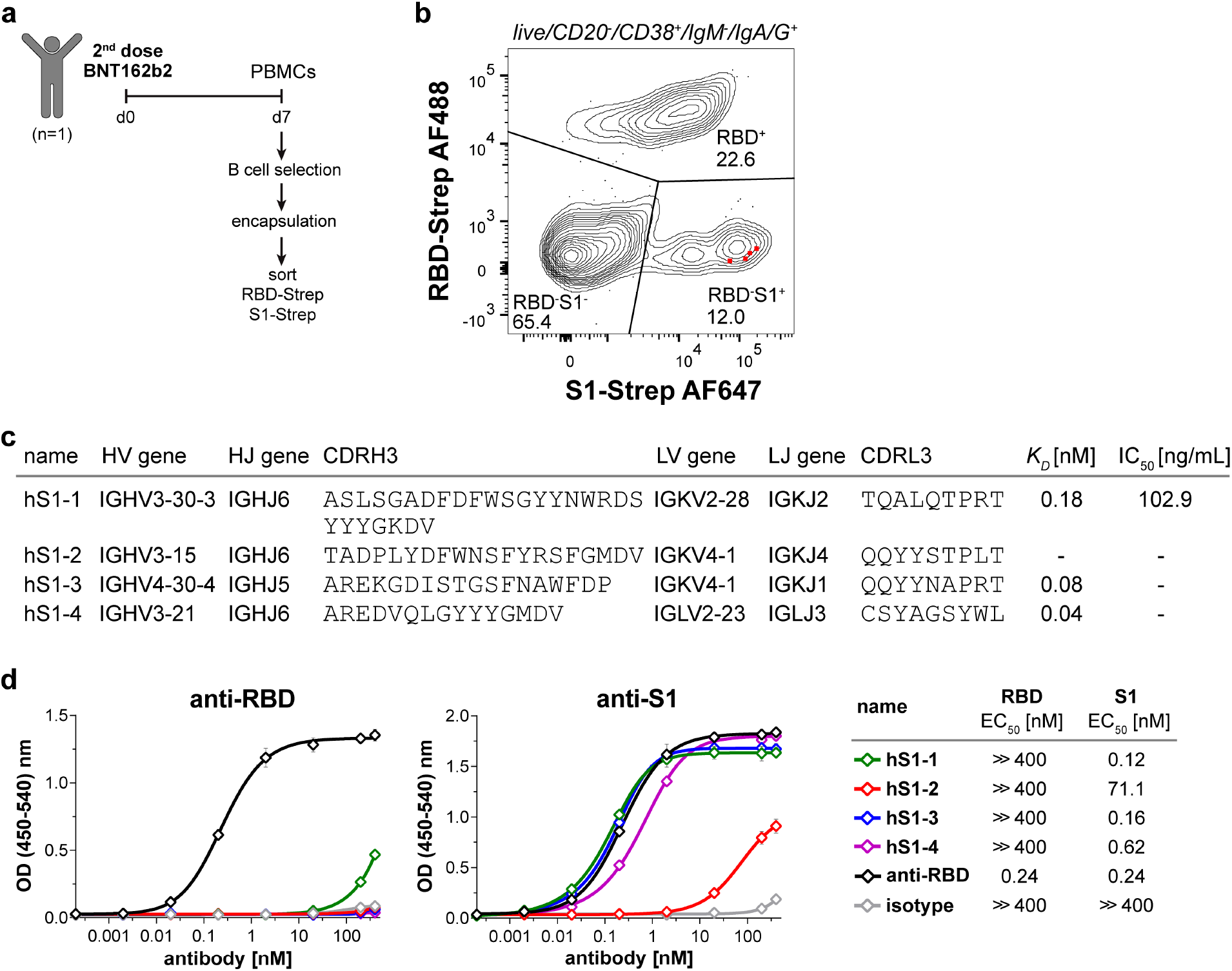
Generation of human anti-SARS-CoV-2 S1 antibodies that do not bind the RBD. (a) Human anti-S1 antibody discovery. B cells were negatively selected from PBMCs 7 days post the second BNT162b2 vaccine dose and then used in the workflow. RBD/S1-specific ASCs were sorted with fluorescently labelled RBD- and S1-streptavidin tetramers. (b) Index sorting of S1-specific human ASCs. Layout for sorting of IgG or IgA secreting ASCs isolated on day 7 post vaccination with fluorescently labelled S1- and RBD-streptavidin tetramers. Based on index sorting, the ratio of S1/RBD fluorescence signal was calculated for all sequenced cells and four sequences that corresponded to events with the highest S1/RBD fluorescence signal ratio were selected for expression (highlighted as red squares). The plot shows 719 events at 5% contour level. (c) Characteristics of human anti-S1 antibodies: variable domain genes (V and J), third complementarity-determining region amino acid sequences (CDR3), equilibrium dissociation constants (K_D_), and half-maximal inhibitory concentrations (IC_50_) against wildtype SARS-CoV-2. Hyphens (-) denote that binding or inhibition was not quantifiable at the tested concentrations. (d) Anti-S1 and -RBD ELISAs of anti-S1 antibodies. An RBD-binding positive control (human IgG1κ, clone AM001414, BioLegend) is shown in black and an isotype control (human IgG1κ, clone QA16A12, BioLegend) is shown in grey. The plots show mean values ± SD of two independent experiments, performed in duplicate. The table shows the median effective concentration (EC_50_), antibody concentration range: 0.0002 – 400 nM.

## Discussion

Our work establishes a fast, simple, versatile, and high-throughput approach for the selection of functional, high affinity antibodies from an underexplored B cell compartment. We generated antibodies directly from ASCs without the need to prepare a DNA library, using straightforward microfluidic equipment and a commercial flow cytometer. Antibodies are screened in the full-length IgG format and functional characterisation can be achieved as quickly as two weeks after obtaining samples by a single researcher without automation. We have illustrated the versatility of this approach by analysing different species (mouse and human) and cells from different tissues (bone marrow plasma cells, splenic ASCs, *ex vivo* stimulated PBMCs, and PBMCs post vaccination).

Our antibody discovery campaigns showcase the ability of our technology to generate high quality murine and human antibodies. The best post-vaccination antibodies we characterised have exceptionally high binding affinities (<10^−12^ M) and neutralising capacities (10 – 100 ng/mL) comparable to a clinical-stage antibody cocktail. Affinities and neutralising capacities were also equivalent to those of memory B cell-derived antibodies isolated by LIBRA-seq after natural infection^54^, highlighting that the results obtained with our technology match those of an established protocol but provide insights into a compartment not yet studied with LIBRA-seq (**Extended Data Figure 8**). Five of our human antibodies also showed neutralising capacity against Omicron BA.1, demonstrating that cross-neutralising activity can be isolated from the plasmablast population after vaccination against WT SARS-CoV-2. Importantly, these binders were isolated from a single human donor without prior screening of serum or plasma for neutralising capacity. In addition, our discovery campaigns were highly efficient, with a true positive rate of >85% for all antibodies and 95% for plasmablast-derived antibodies. In contrast, a recent study performing unbiased sorting of plasmablasts post BNT162b2 booster vaccination yielded on average only 20.3% (42 spike protein binders out of 206 expressed antibodies) of true hits^42^. By enabling antigen-specific sorting, our technology is therefore much more efficient than sorting based on cell surface markers alone.

Leveraging the high throughput of FACS allows screening of >10^7^ hydrogel beads per hour, so even rare antigen-specific ASCs in heterogeneous cell populations can be identified. As highly potent and broadly protective binders can be rare, our high screening capacity increases the likelihood of isolating powerful drug candidates^1^. Another advantage of a flow-cytometry-based readout is that multicolour sorts are a standard procedure, enabling us to test binding to several different antigens or the detection of different antibody isotypes. We have demonstrated sorting with different antigens by selecting antibodies that bind to SARS-CoV-2 spike S1 but not the RBD. Beyond this proof of concept, screening with several antigen variants could be used to identify antibodies against highly conserved epitopes, which increases the likelihood of isolating broadly neutralising antibodies and can help define rational antigens for vaccination^55^. Furthermore, multicolour FACS enables counter selection in the presence of inhibitors or competitors (such as ACE-2)^54^ or screening for orthologue cross-reactivity for testing in animal models^56^. In addition, due to its high throughput, our technology could serve as a fast, economical, and scalable alternative to hybridoma screening. Generation of hybridoma is time-consuming (months) and is usually performed in microtiter plates, which have very low throughput and use more resources than our initial screening format^57^.

The concept of hydrogel-mediated capture of secreted proteins followed by FACS has been proposed before but has not been applied to the selection of functional, recombinant antibodies from natural immune responses^58–60^. Additionally, in contrast to a previously described microdroplet generation method which created polydisperse hydrogel beads of up to 55 µm in diameter^61^, controlled microfluidic encapsulation of cells creates smaller, monodisperse hydrogel beads, which facilitates sorting. The combination of single cell microfluidic encapsulation with FACS also has advantages over methods solely based on microfluidics. Compared to fully microfluidic workflows, we offer a simpler alternative that achieves higher throughput without extensive setup or optimisation. FACS instruments are available in most institutes, can achieve sorting rates more than ten times higher than droplet microfluidic sorters previously used for antibody discovery (200 - 600 Hz)^17,18,62^ and enable more complex, multi-colour sorts. The throughput of our approach is also orders of magnitude higher than that of microchamber-based microfluidic systems (1,000s – 10,000s of cells per chip)^63–65^. While some of these approaches have delivered functional antibodies, they are less accessible to researchers, as the required equipment is prohibitively expensive (e.g. Berkeley Light’s Beacon) or proprietary (e.g. AbCellera’s platform).

Compared to methods that employ cell surface capture combined with FACS^32,33^, we propose a defined system in which we can control the amount of capture reagent independent of cell-surface molecules and without modifying the cell surface. In our system, the hydrogel around the cell captures enough antibodies to be used with both monomeric (OVA) or tetrameric (RBD-strep, S1-strep) antigens without the signal amplification required in previously described methods^33^.

In addition to being a tool for antibody discovery, our technology can offer fundamental insights into immunobiology. By facilitating access to the ASC compartment, we can now look into contrasting the affinities, variant specificities, and biophysical properties of ASC-derived antibodies relative to those of other compartments. In this initial study, we have outlined how our antigen-specific plasmablast sequence profiles can be compared with anti-SARS-CoV-2 antibodies reported in studies on different individuals/B cell compartments to offer finer-grain insights into immunodominance – the phenomenon that underpins convergent viral mutation. This understanding can be used to prioritise candidates tactically, considering both their functional efficacy and their likelihood of withstanding viral natural selection pressures^66^. Finally, the modular design of our capture system (BG-agarose and SNAP-tag protein fusions) allows rapid adaptation to other secreted proteins, for example antibodies from other species^26^ or cytokines. We hypothesize that light-chain mediated capture, as employed in our system, could also enable interrogation of other characteristics of the secreted antibodies, such as glycosylation^67^.

Our technology is currently limited by the throughput of cell encapsulation (up to 10^7^/hour) and low sequencing yield. Cell encapsulation can be further optimised by adjusting the mean number of cells per droplet and by running more microfluidic chips in parallel. While we have used full-length Sanger sequencing for simplicity, our workflow can easily be adapted to next-generation sequencing technologies^68,69^. We aim to increase sequencing output by optimising single cell sequencing protocols and by combining our sorting technology with the LIBRA-seq 10X genomics workflow that uses DNA barcoded antigens^23^ which should also yield single-cell transcriptomic datasets. These improvements will give us the ability to construct sequence-function landscapes for thousands of ASC-derived antibody sequences^70^.

In summary, we present a fast, viable, low-cost tool for the isolation of (human) monoclonal antibodies that can be used without much training not only in industry but also in non-specialist research laboratories. Our technology enabled the discovery of hundreds of antibodies within weeks and had a high recovery rate of true positives (>85%). Through this more efficient hit discovery, resources can be shifted from binder identification to more thorough experimental characterisation and computational assessment^71^. Additionally, the speed and versatility of this method enable rapid adaptation in the face of emerging pathogen variants. As disease outbreaks become more likely in the future^72^, we believe this technology increases our pandemic preparedness by enabling faster access to antibody reagents.

## Acknowledgements

We would like to thank the flow cytometry facility from the School of the Biological Sciences for their support and assistance in this work, K. Stott from the biophysics facility of the Department of Biochemistry for help with BLI measurements, J. Holstein for help with chemical synthesis of BG-agarose, and members of the Hollfelder laboratory for comments on the manuscript. K.F was supported by the MRC, the Cambridge European Trust and St. John’s College Cambridge. M.I.J.R received a Postdoctoral Research fellowship by Boehringer Ingelheim. T.S.K. held an EU H2020 Marie Skłodowska-Curie Individual Fellowship (MSCA-IF 750772). T.N.K. received an AstraZeneca studentship. N.J.M was supported by the MRC (TSF ref. MR/T032413/1), NHSBT (grant ref. WPA15-02), the Wellcome Trust (ISSF ref. 204845/Z/16/Z) and Addenbrooke’s Charitable Trust (grant ref. 900239). This work was supported by the EU Horizon 2020 programme (ERC Advanced Investigator Awards to F.H., 69566), the MRC (grant ref. MC_UU_00025/12, to J.E.D.T), the Medical Research Foundation (MRF-057-0002-RG-THAV-C0798 to J.E.D.T.), and the NIHR Cambridge BRC.

## Author contributions

K.F. and F.H. conceptualised the study. K.F., A.L., T.Y.S., P.P.-G., T.N.K, T.S.K., J.C.Y.-P., R.H., F.L.-M., and P.B. performed experiments. M.I.J.R. developed software and analysed data. K. F. and F.H. wrote the manuscript with input from all authors. F.H., J.E.D.T., M.H, C.M.D and N.J.M. supervised the work.

## Data Availability section

The SI contains amino acid sequences of all antibodies expressed in this study, oligonucleotide sequences for RT-PCR and crystallographic data summarizing structural and refinement statistics. Sequences of experimentally verified antigen-specific antibodies were also deposited at Genbank (accession numbers: OQ208846 - OQ208931). Sequences of anti-SARS-CoV-2 antibodies were deposited in the CoV-AbDab. Crystallographic data were deposited at RCSB PDB with ID <u>8BE1</u>. A file of the microfluidic chip design is available on <u>DropBase</u>.

## Extended Data

**Extended Data Fig. 1.**
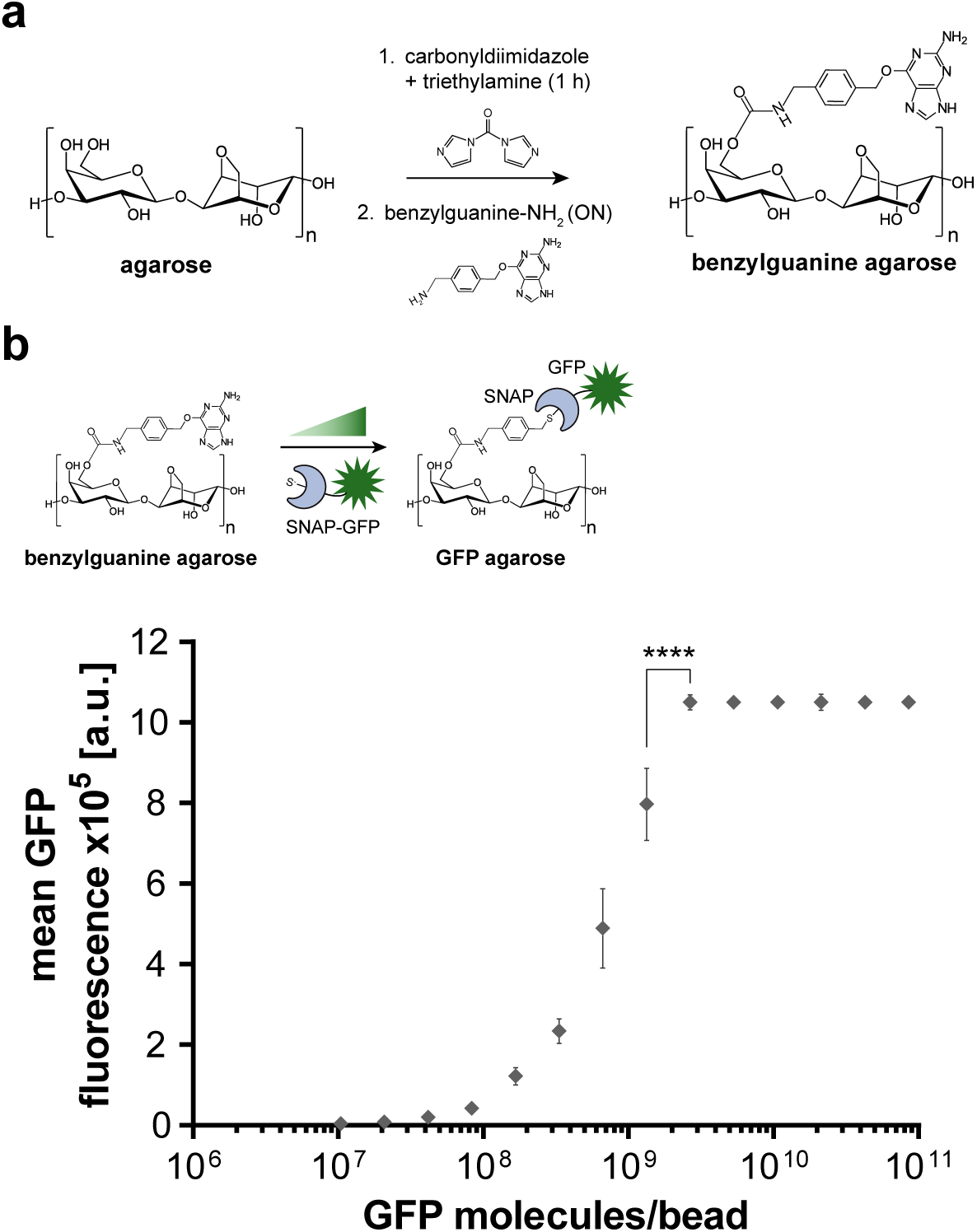
Synthesis and characterisation of benzylguanine (BG)-agarose. (a) Synthesis of BG-agarose. (b) Assessment of 1.5% (w/v) BG-agarose capture valency. BG-agarose beads were incubated with different amounts of GFP-SNAP and the GFP-fluorescence of the beads was analysed by flow cytometry. The fluorescence signal saturates between 1.3 – 2.7 × 10^9^ GFP molecules/bead. Error bar represents SD, at least 18,000 beads were analysed per condition (two tailed p < 0.0001, unpaired t-test).

**Extended Data Fig. 2.**
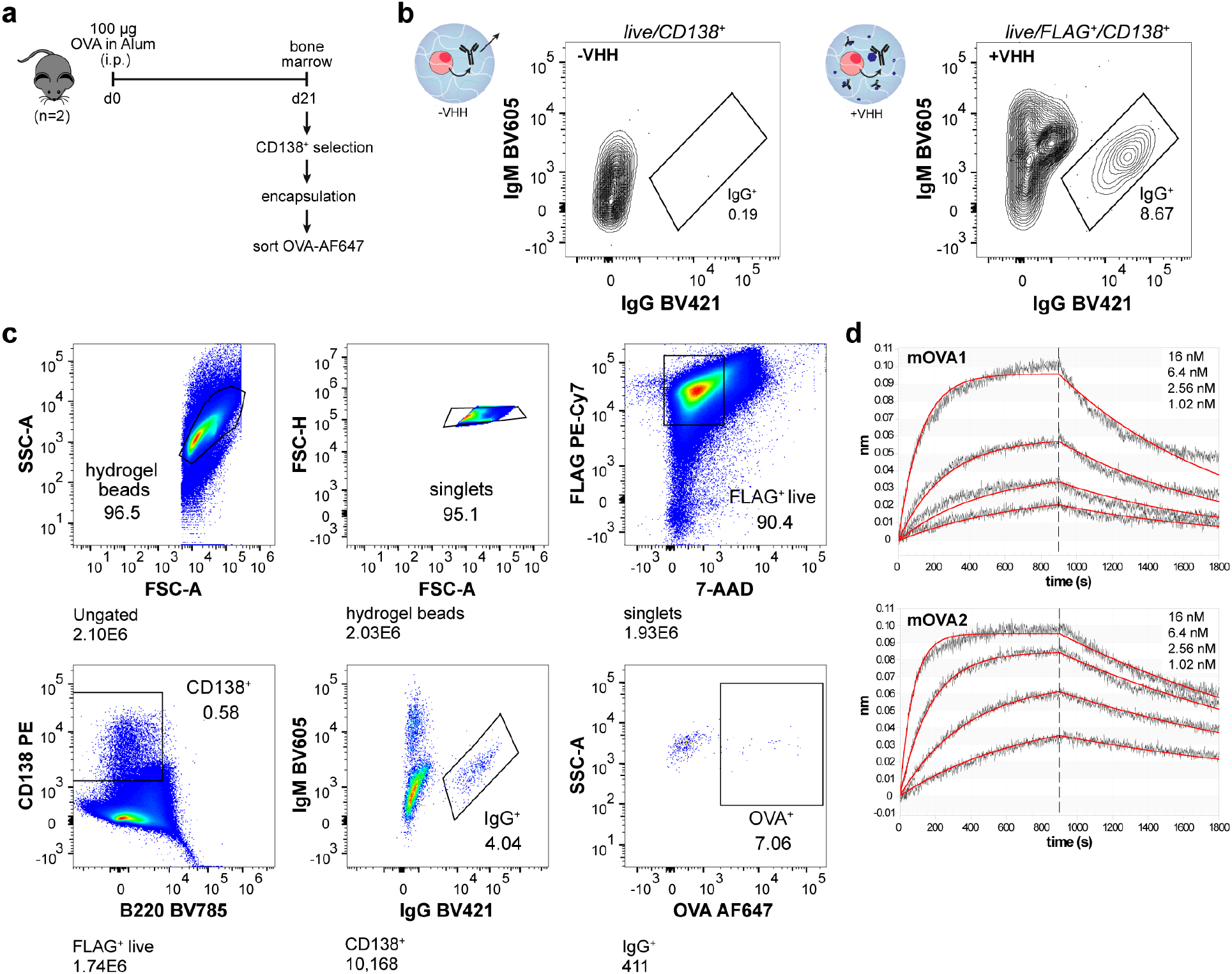
Generation of anti-OVA antibodies from mouse bone marrow plasma cells. (a) Mouse immunisation and analysis scheme. Bone marrow plasma cells (CD138^+^) were magnetically enriched and encapsulated into VHH-functionalised BG-agarose. OVA-specific plasma cells were sorted with fluorescently labelled monomeric OVA (OVA-AF647). (b) Comparison of IgG signal originating from plasma cells in BG-agarose beads incubated with (+VHH) or without VHH-SNAP-FLAG (-VHH) (different experiment than shown in **Figure 1**). The plots show 527 (-VHH) and 1,280 (+VHH) events at 2% contour level. (c) Representative gating strategy for identifying OVA-specific plasma cells from bone marrow. Live plasma cells inside hydrogel beads were gated as live/FLAG^+^/CD138^+^ (VHH-SNAP contains a FLAG tag). (d) Determination of binding affinity of mOVA1 and mOVA2 by biolayer interferometry: mOVA1: K_D_ = 2.36 ± 0.02 nM, χ^2^ = 0.0705, R^2^ = 0.985. mOVA2: K_D_ = 0.684 ± 0.004 nM, χ^2^ = 0.0388, R^2^ = 0.9919. Tested OVA concentrations: 1.02 – 16 nM.

**Extended Data Fig. 3.**
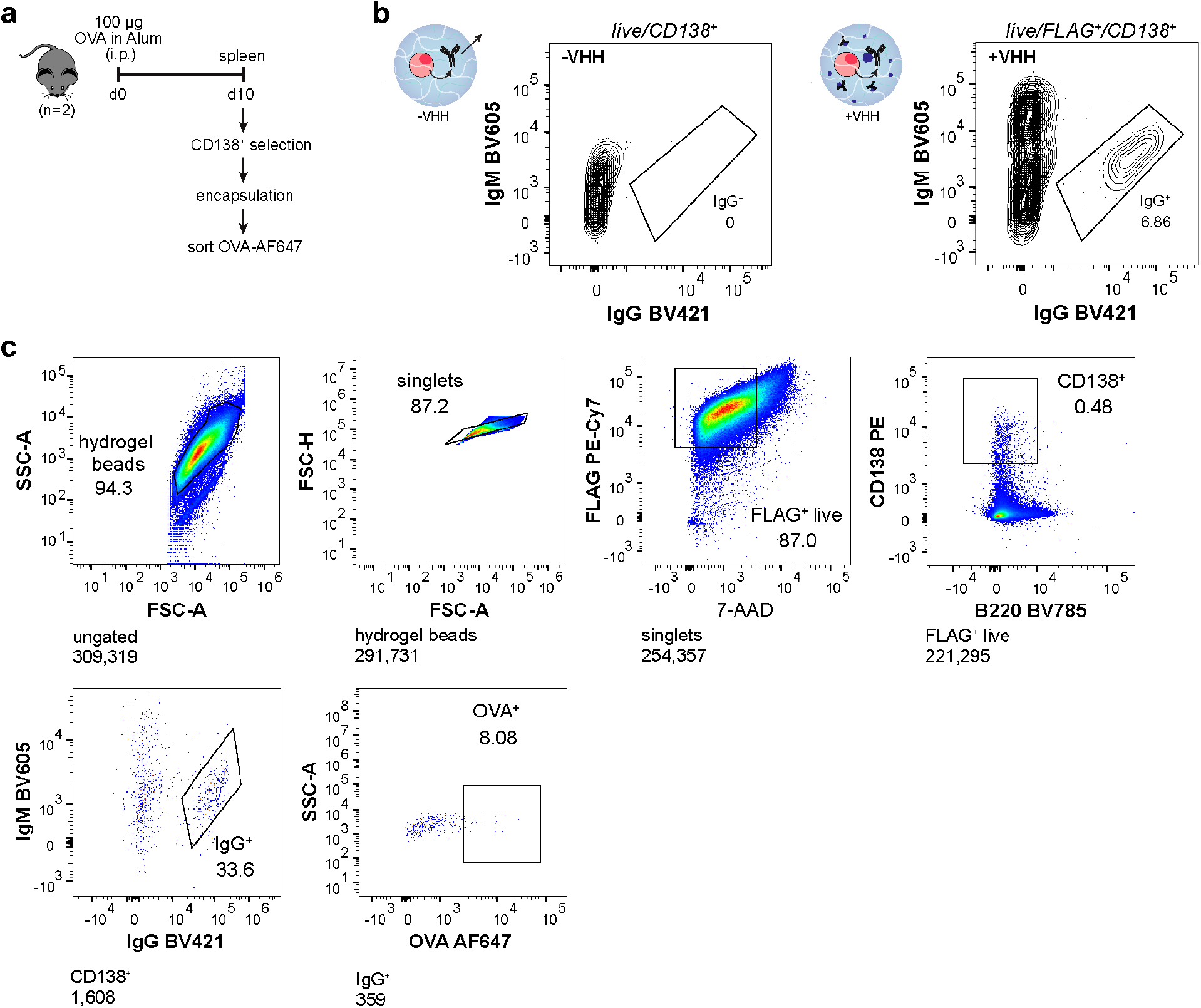
Workflow with spleen-derived ASCs. (a) Mouse immunisation and analysis scheme. (b) Comparison of IgG signal originating from ASCs in BG-agarose beads incubated with (+VHH) or without VHH-SNAP-FLAG (-VHH) (different experiment than shown in **c**). The plots show 3,663 (-VHH) and 2,290 (+VHH) events at 2% contour level. (c) Representative gating strategy for identifying OVA-specific ASCs from spleen. Live ASCs inside hydrogel beads were gated as live/FLAG^+^/CD138^+^ (VHH-SNAP contains a FLAG tag).

**Extended Data Fig. 4.**
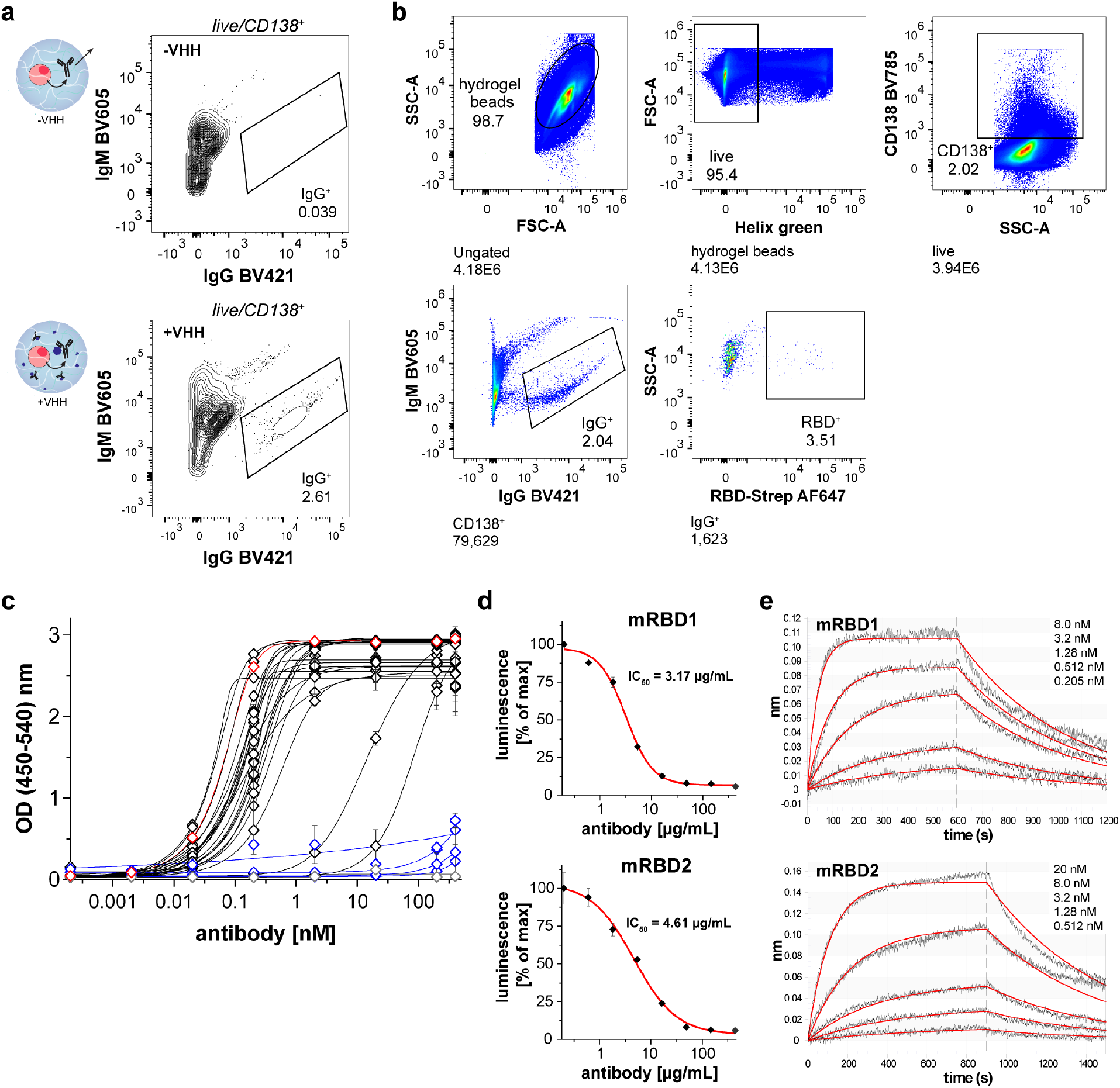
Sorting of murine RBD-specific plasma cells and characterisation of anti-RBD antibodies. (b) Comparison of IgG signal originating from plasma cells in BG-agarose beads incubated with (+VHH) or without VHH-SNAP (-VHH) (different experiment than shown in **Figure 2**). The plots show 2,536 (-VHH) and 14,606 (+VHH) events at 2% contour level. (b) Representative gating strategy for identifying RBD-specific plasma cells from mouse bone marrow (gated as live/CD138^+^/IgM^-^/IgG^+^/RBD^+^). (c) Anti-RBD ELISA. The plot shows representative data of two technical replicates (mean ± SD). Workflow-derived antibodies are shown in black (binders) or blue (non-binders), a commercial positive control (clone 1035753, R&D systems) is shown in red and an isotype control (mouse IgG1k, clone MG1-45, BioLegend) is shown in grey. (d) Neutralisation of SARS-CoV-2 by mRBD1 and mRBD2. Wildtype SARS-CoV-2 (MOI=0.01) was pre-incubated with a 3-fold dilution series of each antibody, then used to infect luminescent reporter cells. Levels of infection after 24 h were quantified as % of maximum luminescence. Mean values ± SD of three technical replicates are shown, representative of three independent experiments. IC_50_, half-maximal inhibitory concentration. (e) Determination of binding affinity of mRBD1 and mRBD2 by biolayer interferometry: mRBD1: K_D_ =0.864 ± 0.07 nM, χ^2^ = 0.0547, R^2^ = 0.991. mRBD2: K_D_ = 4.78 ± 0.04 nM, χ^2^ = 0.1111, R^2^ = 0.9928.

**Extended Data Table 1:**
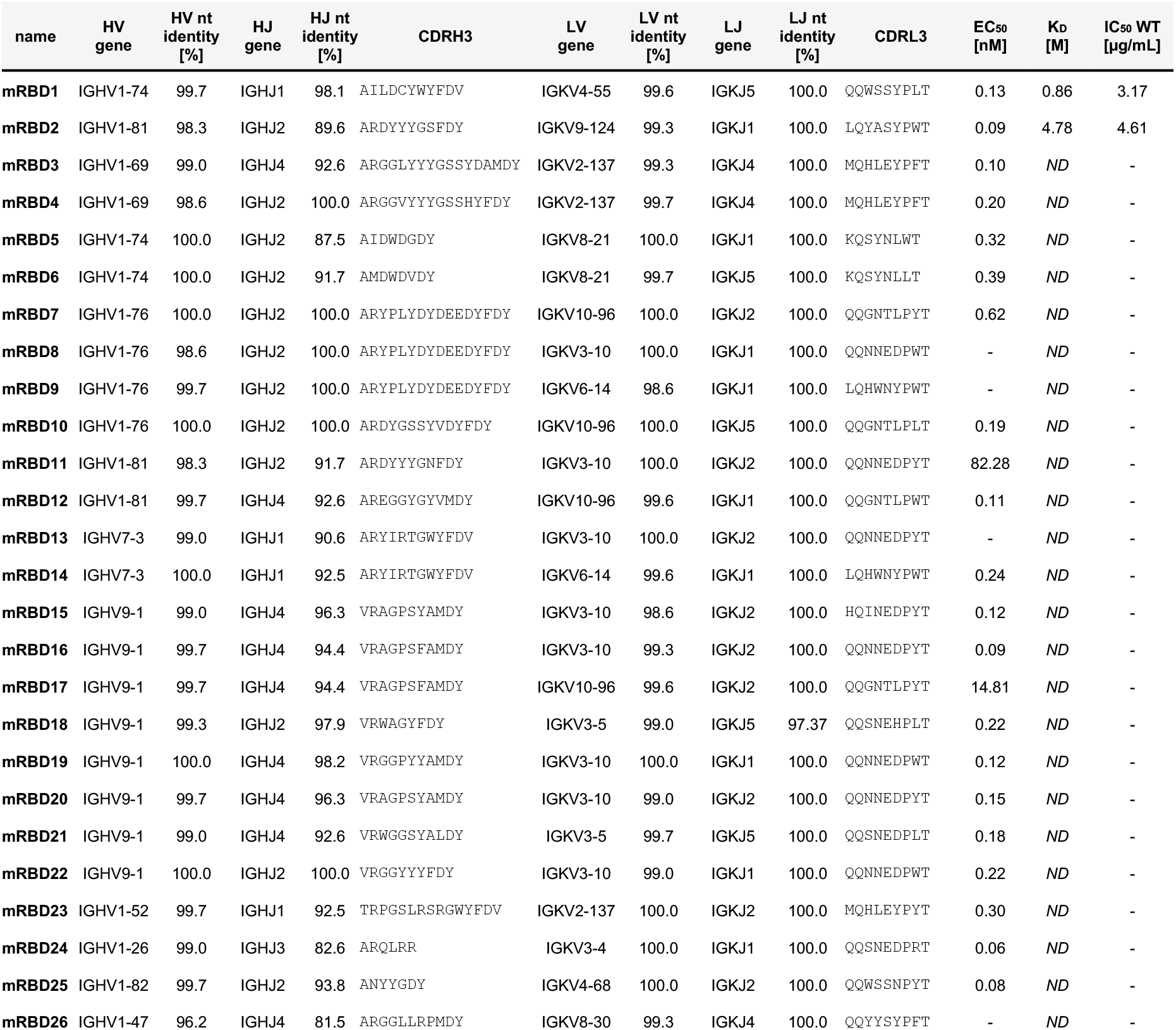
Characteristics of expressed mouse anti-RBD antibodies. The table summarises variable domain sequences (V and J genes), percentage nucleotide identity to germline, third complementarity-determining regions (CDRs), ELISA half maximal effective concentrations (EC_50_), equilibrium dissociation constants (K_D_) and half-maximal inhibitory concentrations (IC_50_) against wildtype SARS-CoV-2. Hyphens (-) denote that binding or inhibition was not quantifiable at the tested concentrations, ND = not determined. Full length sequences of the antibodies can be found it **Supplementary Table 4**.

**Extended Data Fig. 5.**
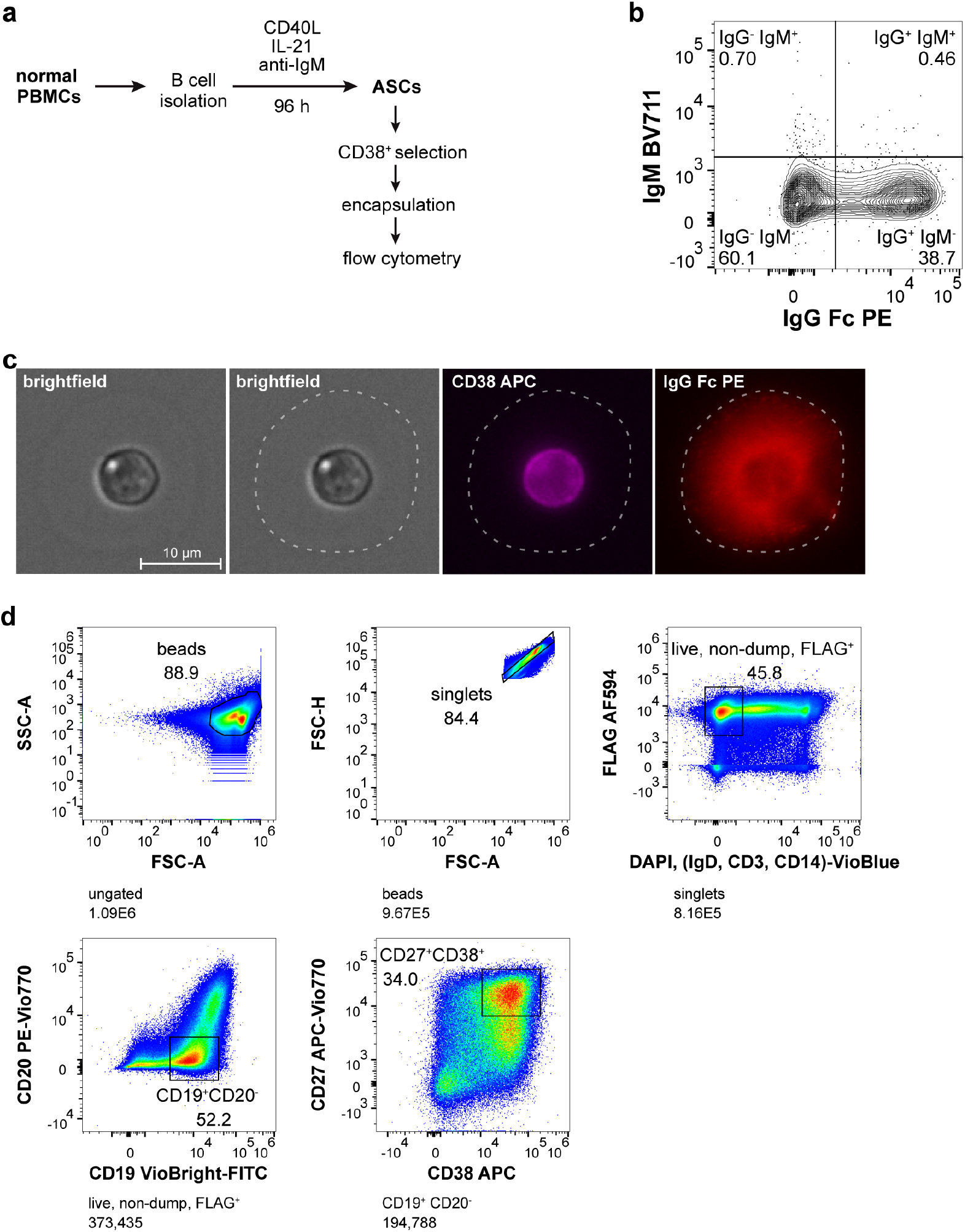
Stimulation of normal PBMCs into ASCs and analysis of antibody secretion by flow cytometry and microscopy. (a) Stimulation conditions and workflow for proof-of-concept in normal PBMCs. To obtain ASCs from normal PBMCs, B cells were stimulated with CD40L, IL-21 and anti-IgM antibody. (b) IgG secretion of stimulated PBMCs encapsulated into hydrogel beads. The plot shows 66,173 events at a contour level of 2%, gating strategy shown in (d). (c) Analysis of encapsulated stimulated PBMCs by epifluorescence microscopy. Hydrogel beads were stained with fluorescently labelled anti-CD38 (APC) and anti-human IgG antibodies (PE). The hydrogel bead boundary is shown as a dotted line. (d) Gating strategy for identifying IgG secreting human ASCs from stimulated B cells. ASCs were gated as DAPI^-^/CD14^-^/CD3^-^/IgD^-^/FLAG^+^/CD19^+^/CD20^-^/CD27^+^/CD38^+^ (VHH-SNAP contains a FLAG tag).

**Extended Data Fig. 6.**
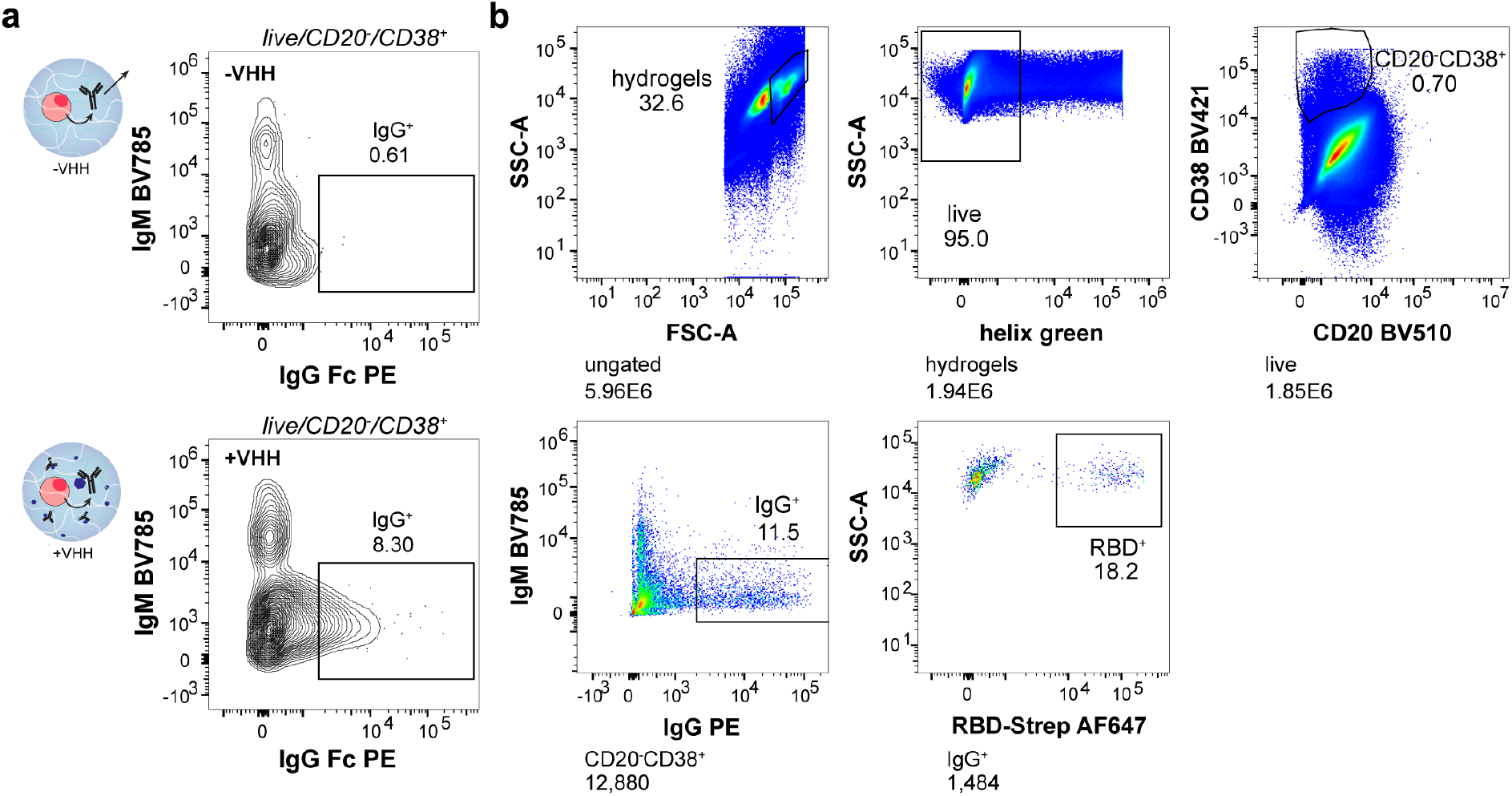
Sorting of SARS-CoV-2 RBD-specific human ASCs. (a) Comparison of IgG signal originating from ASCs in BG-agarose beads incubated with (+VHH) or without VHH-SNAP (-VHH) (different experiment than shown in **Figure 3**). The plots show 1,139 (-VHH) and 1,289 (+VHH) events at 2% contour level. (b) Representative gating strategy for identifying RBD-specific human ASCs (gated as live/ CD20^-^/CD38^+^/IgM^-^/IgG^+^/RBD^+^). Cells in hydrogels could be easily identified by forward and side scatter, therefore the gating is different here compared to other experiments.

**Extended Data Table 2.**
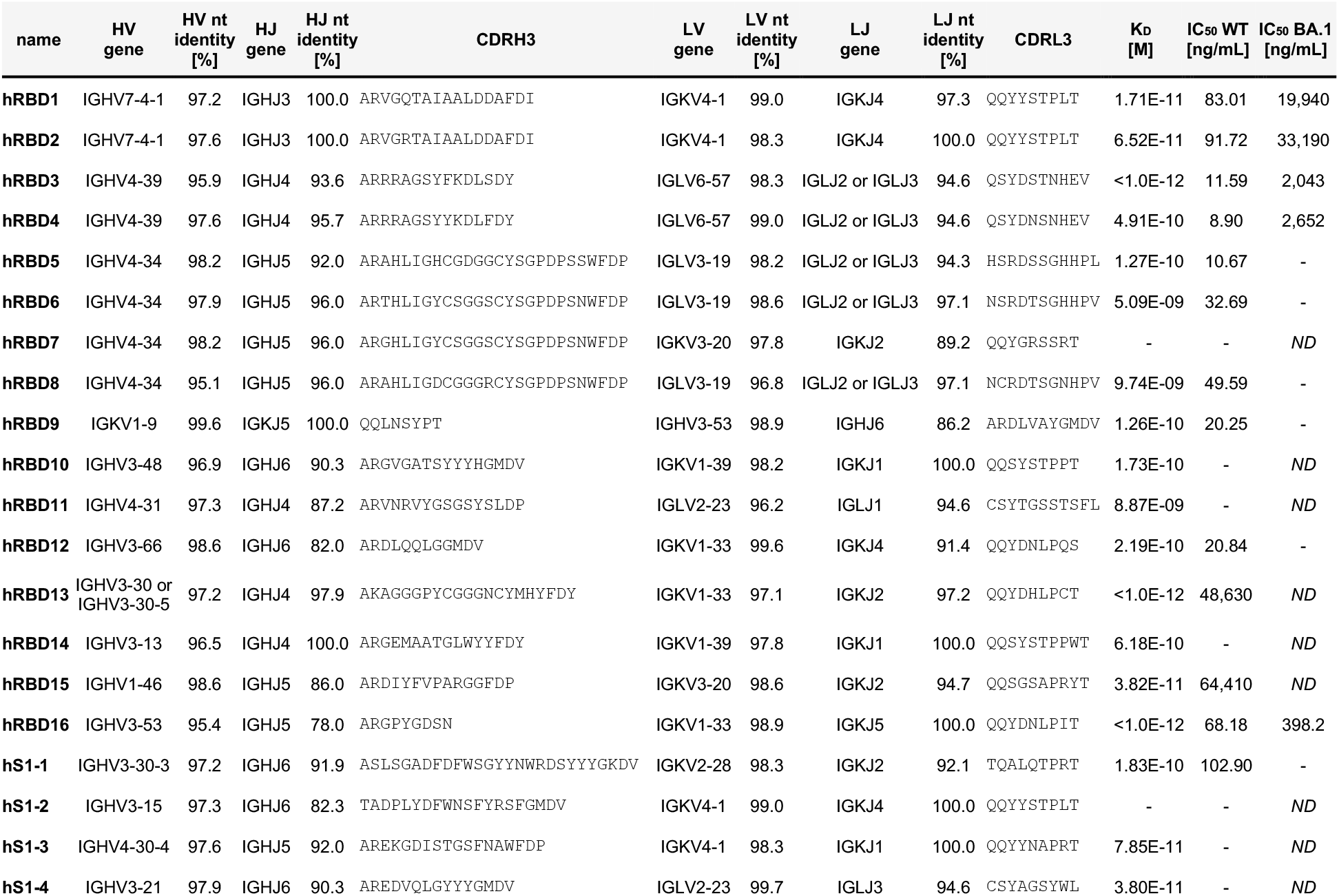
Characteristics of expressed human antibodies. Characteristics of human anti-SARS-CoV-2 antibodies expressed as full-length IgG1: variable domain sequences (V and J genes), percentage nucleotide identity to germline, third complementarity-determining region (CDR3s), equilibrium dissociation constant (K_D_) and half-maximal inhibitory concentrations (IC_50_) against wildtype SARS-CoV-2 and Omicron BA.1. Hyphens (-) denote that binding or inhibition was not quantifiable at the tested concentrations, ND = not determined. Full length sequences of the antibodies can be found it **Supplementary Table 5**.

**Extended Data Fig. 7.**
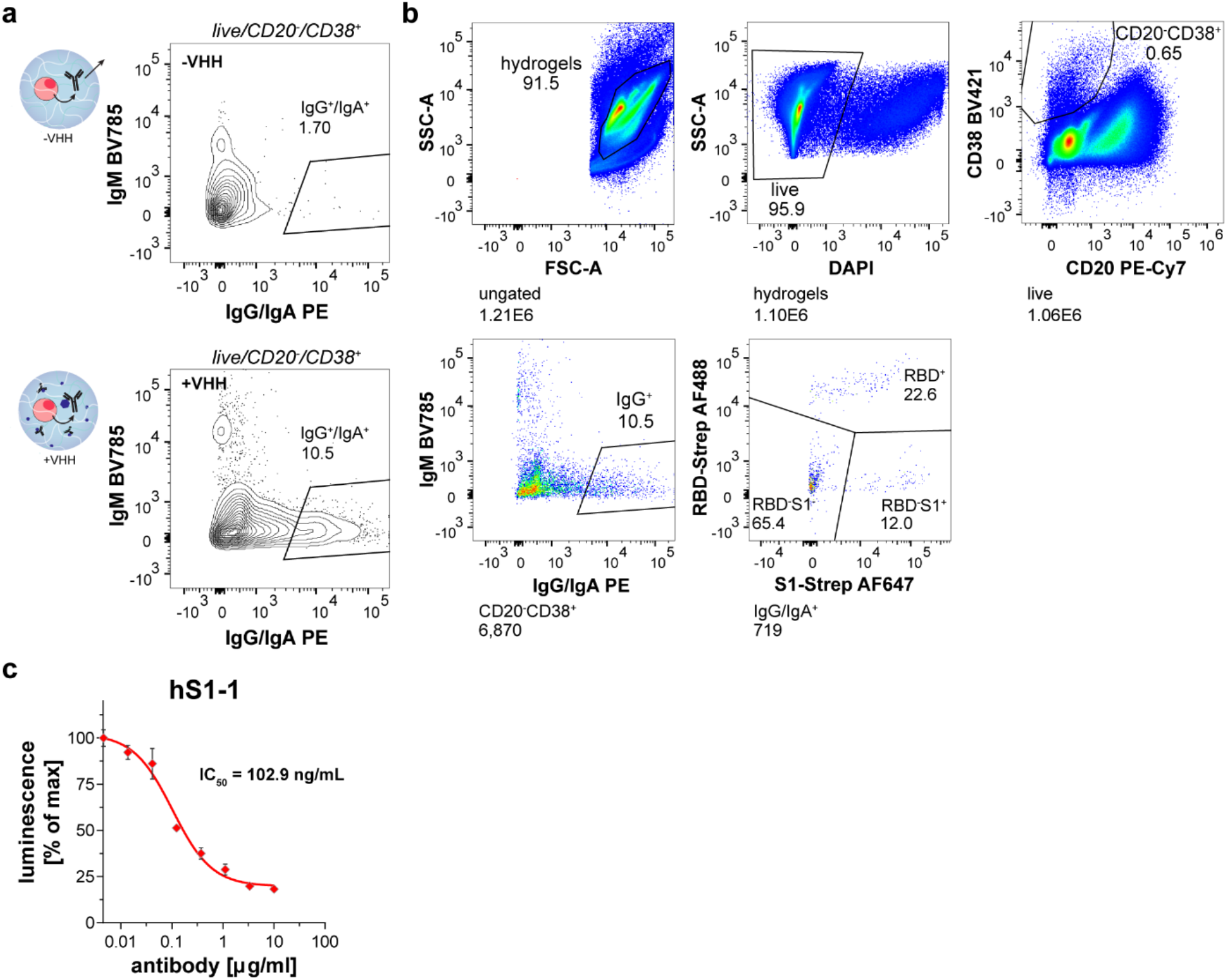
Sorting of SARS-CoV-2 S1-specific human ASCs. (a) Comparison of IgG signal originating from ASCs in BG-agarose beads incubated with (+VHH) or without VHH-SNAP (-VHH). The plots show 823 (-VHH) and 6,870 (+VHH) events at 2% contour level. (b) Representative gating strategy for identifying S1/RBD-specific human ASCs. (c) Neutralisation of SARS-CoV-2 by hS1-1. Wildtype SARS-CoV-2 (MOI=0.01) was pre-incubated with a 3-fold dilution series of hS1-1, then used to infect luminescent reporter cells. Levels of infection after 24 h were quantified as % of maximum luminescence. Mean values ± SD of two technical replicates are shown, representative of two independent experiments. IC_50_, half-maximal inhibitory concentration.

**Extended Data Fig. 8.**
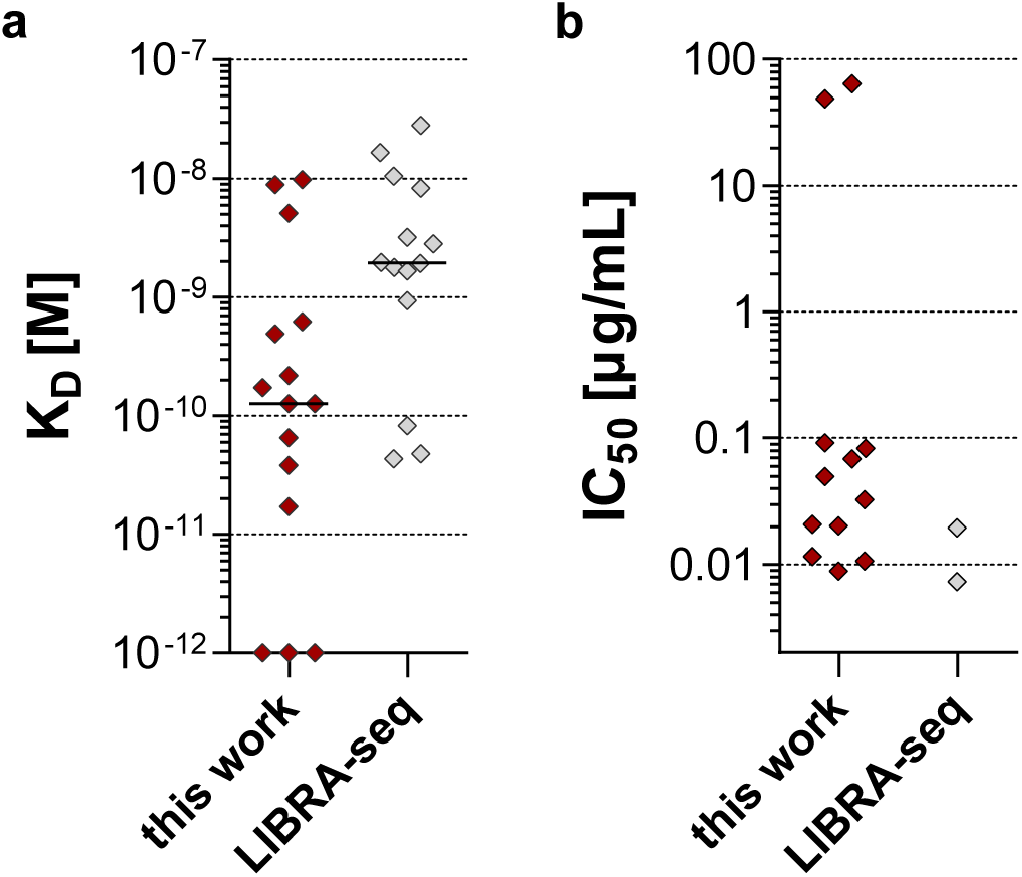
Comparison of human anti-SARS-CoV-2 RBD antibodies isolated in this work with antibodies isolated by LIBRA-seq^54^. (a) Comparison of affinities (K_D_). All characterised RBD binders were included in the comparison. For antibodies isolated in this work, the same K_D_s are depicted in **Fig. 3d**. For antibodies isolated by LIBRA-seq, previously reported values are shown. Horizontal lines denote the median. (b) Comparison of neutralising capacities. Only antibodies tested against authentic SARS-CoV-2 were included in the comparison. For antibodies isolated in this work, the same IC_50_s are depicted in **Fig. 3e**. For antibodies isolated by LIBRA-seq, previously reported values are shown. Different neutralisation assays were used in the two studies.

## Methods

### Ethics statement

Animal experiments were licensed by the UK Home Office according to the Animals Scientific Procedures Act 1986 (License PP6047951) and approved by local ethics committees from the University of Cambridge. Human sample collection and analysis was conducted in accordance with the principles of Good Clinical Practice and following approved protocols of the NIHR National Bioresource. Samples were collected with the written informed consent of all study participants under the NIHR National BioResource-Research Tissue Bank (NBR-RTB) ethics (REC:17/EE/0025).

### Microfluidic methods

The microfluidic chip was designed using AutoCAD (Autodesk) and printed on a high-resolution photomask (Micro Lithography Services). The channel layout of the microfluidic chip (**Supplementary Figure 1**) is available as a .dxf file from <u>DropBase</u>, to manufacture a chip in-house or by outsourcing. Polydimethylsiloxane (PDMS) microfluidic chips were fabricated using standard soft lithography^73^. For replica moulding, approximately 20 gram of PDMS (Sylgard 184, Dow Corning) monomer and curing agent were mixed at a ratio 10:1, and degassed, poured onto the master and solidified overnight at 65 °C. PDMS slabs were cut from the master, then inlets and outlets were created using a 1 mm biopsy punch (kai medical) and the PDMS slab was permanently bonded to glass slides through plasma treatment (Diener Femto). After bonding, microfluidic channels were flushed with 1% (v/v) trichloro(1H,1H,2H,2H-perfluoro-octyl)silane (Sigma-Aldrich) in HFE-7500 (3M Novec) to make them more hydrophobic. Flow rates in microfluidic workflows were controlled using neMESYS 290N syringe pumps (Cetoni) and gas-tight syringes (Hamilton) connected to PTFE tubing (inner diameter 0.5 mm, BOLA). Droplet formation was monitored using a high-speed Mini UX-100 camera (Photron) mounted onto an inverted microscope (Brunel SP981 Microscope).

### Production of VHH-SNAP

Sequences of VHHs (TP1170^26^ as anti-mouse κ VHH, HuFab kappa 1^27,74^ as anti-human κ VHH and HuFab lambda 1^27,74^ as anti-human λ VHH) were obtained as gene fragments (Integrated DNA Technologies, Leuven, Belgium). Expression plasmids based on the pET28a backbone (Novagen) were constructed by Gibson Assembly^75^. Amino acid sequences of all VHH-SNAP constructs can be found in **Supplementary Table 1**. For the capture of mouse antibodies, we focused on anti-κ VHH as 95% of mouse B cells express κ light chains^76^. Proteins were expressed in *E. coli* BL21(DE3) in 2xYT supplemented with 50 µg/mL kanamycin. Cultures were grown at 30 °C, protein expression was induced at OD_600_ 0.6 – 0.8 with 0.25 mM IPTG and proteins were expressed for 5 h at 24 °C. After harvesting, the cell pellet was resuspended in His-wash buffer (50 mM Tris/HCl, pH 7.5, 300 mM NaCl, 20 mM imidazole) and supplemented with protease inhibitor (cOmplete Protease Inhibitor Cocktail, Roche) and benzonase nuclease (Sigma) according to the manufacturer’s instructions. Cells were lysed using 10X BugBuster (Merck) according to the manufacturer’s instructions. Cleared lysate was added to a Ni-NTA column (Ni-NTA affinity resin, Generon) pre-equilibrated with His-wash buffer. Proteins were eluted with 50 mM Tris/HCl, pH 7.5, 300 mM NaCl, 500 mM imidazole. To achieve higher purity, a reverse nickel column purification was performed. Proteins were re-buffered into TEV cleavage buffer (25 mM Tris/HCl, pH 8.0, 150 mM NaCl) using PD-10 columns (GE) and subjected to overnight cleavage at 4 °C with His-tagged TEV protease (Sigma) according to the manufacturer’s instructions. Cleaved proteins were then re-run on a Ni-NTA column to remove the His-tagged TEV protease and the cleaved epitope tag. Flow-through and wash fractions containing the purified VHH-SNAP were collected. The buffer was exchanged to 10% glycerol in PBS using PD-10 desalting columns (GE) and protein samples were aliquoted, snap frozen in liquid nitrogen and stored at -80 °C.

### Production of SARS-CoV-2 RBD-CHis for immunization

Synthetic, codon-optimised DNA encoding for the receptor-binding domain of WT SARS-CoV-2 was cloned into pExp-CHis vector which introduces a C-terminal octa-His-tag (RBD-CHis, sequence in **Supplementary Table 1**). RBD-CHis protein was expressed in *E. coli* BL21(DE3) in 2xYT medium supplemented with 100 µg/ml ampicillin at 37 °C. Protein expression was induced at OD_600_ 0.8 – 1.0 by addition of 0.4 mM IPTG and progressed for 16 h. Cells were pelleted by centrifugation, lysed using Avestin EmulsiFlex C-5 homogenizer in 50 mM Tris-HCl pH 8.0, 5 mM EDTA supplemented with 250 µg/ml of DNaseI. Inclusion bodies were isolated by centrifugation, washed first with 50 mM Tris-HCl pH 8.0, 5 mM EDTA, 1 % Triton-X 100 buffer and then with 50 mM Tris-HCl pH 8.0, 1 M NaCl, 5 mM EDTA buffer. Then inclusion bodies were solubilised in 50 mM Tris-HCl pH 8.0, 5 mM EDTA, 10 mM TCEP and 6 M guanidine hydrochloride. For refolding, the protein solution was added slowly to 50 mM Tris-HCl pH 8.5, 50 mM ethanolamine, 1 M PPS, 2 mM EDTA, 2 mM cysteine and 0.2 mM cystine and incubated for 96 h at 4 °C. The refolded protein was first purified by reverse-phase chromatography using 10 mL Source RPC column and applying a 10-90% acetonitrile/0.1% TFA gradient. The peak fractions were pooled, and the protein was lyophilised. Then the protein was dissolved in 20 mM sodium phosphate pH 7.2, 300 mM NaCl, 0.5 M urea and was further purified by size-exclusion chromatography using Superdex 200 Increase 10/300 GL column (Cytiva) equilibrated in 20 mM sodium phosphate pH 7.2, 300 mM NaCl.

### Production of SARS-CoV-2 RBD for crystallisation

Synthetic DNA encoding for WT SARS-CoV-2 RBD was cloned into pExp-His-Zbasic plasmid to result in a fusion of RBD with N-terminal His-tag and Zbasic fusion partner followed by a TEV protease cleavage site. The fusion protein was expressed in *E. coli* BL21(DE3) in 2xYT media with 100 µg/ml ampicillin. Cells were grown to OD_600_ 0.8 – 1.0 and expression induced with 0.4 mM IPTG. Cells were pelleted by centrifugation after 5 hours of expression at 37 °C and resuspended in 20 mM Tris-HCl pH 8.0, 100 mM NaCl. Before lysis, 250 µg/ml of DNaseI and 1 mM PMSF were added to the cells and cells lysed by sonication. Lysate was cleared by centrifugation, supernatant discarded, and homogenised inclusion bodies were washed twice: first with 20 mM Tris-HCl pH 8.0, 10 mM EDTA, 1 mM TCEP, 1 % Triton-X 100 and then with 20 mM Tris-HCl pH 8.0, 1 M NaCl, 1 mM TCEP. The final pellet was dissolved in 20 mM Tris-HCl pH 8.0, 200 mM NaCl, 10 mM imidazole, 1 mM TCEP, 6 M guanidine-HCl and insoluble material was removed by centrifugation. Solubilised, denatured protein was purified using a PureCube Ni-NTA agarose column. After loading the protein, the column was washed with 10 column volumes (CV) of 20 mM Tris-HCl pH 8.0, 500 mM NaCl, 10 mM imidazole, 6 M deionised urea, 0.5 mM TCEP and protein eluted with 5 CV of 20 mM Tris-HCl pH 8.0, 100 mM NaCl, 250 mM imidazole, 6 M deionised urea, 0.5 mM TCEP. The fusion protein was refolded by diluting it 1:20 in dropwise fashion, with stirring at 4°C, into refolding buffer 100 mM Tris, 500 mM Arg-HCl, 0.5 g/l PEG 3350, 1 M urea, 2 mM cysteine, 0.2 mM cysteine. After 72h of refolding, the solution was diluted 2-fold with 50 mM MES-NaOH pH 6.0 and filtered through a Whatman GF/B microfiber filter. Clarified protein solution was loaded onto HiTrap SP HP cation-exchange column (Cytiva), column washed with 5 CV of 20 mM HEPES-NaOH pH 7.2, 50 mM NaCl and eluted with 25 CV linear gradient to 20 mM HEPES-NaOH pH 7.2, 1000 mM NaCl. Peak fractions were pooled and NHis-TEV protease (produced in-house) was added for digestion of the fusion protein overnight at 4°C. Cleaved protein was passed through a Ni-NTA column to remove the His-Zbasic fusion partner, uncleaved fusion protein and NHis-TEV protease. RBD from the flow-through was concentrated using 10 kDa MWCO Amicon Ultra concentrator and loaded onto HiLoad Superdex 75 pg 16/60 size-exclusion column (Cytiva) equilibrated with 20 mM Tris-HCl pH 7.2, 200 mM NaCl. Pooled peak fractions were concentrated to 15-16 mg/ml using Amicon Ultra concentrators as before, flash frozen in aliquots and stored at –80 °C.

### Production of biotinylated RBD

Synthetic DNA encoding for WT SARS-CoV-2 RBD was cloned into pExp-His-Zbasic-Avi plasmid to result in a fusion of RBD with N-terminal His-Zbasic-TEV fusion partner and appended C-terminal Avi-tag (RBD-Avi, sequence in **Supplementary Table 1**). The Avi-tagged RBD was produced following essentially the same procedure as untagged RBD except for addition of 5 mM MgCl2, 2 mM ATP, 150 μM biotin and 50 μg/mL of biotin ligase (BirA-CHis, produced in-house) during the TEV protease cleavage step for the site-specific biotinylation at the Avi-tag. The reaction proceeded for 16 h at 22°C with gentle mixing.

### Antigen tetramerisation

For the streptavidin-RBD complex formation, a 10X molar excess of biotinylated RBD-Avi protein was incubated with 0.5 mg/ml of streptavidin-AlexaFluor 488 or streptavidin-AlexaFluor 647 (BioLegend) for 1 hour at room temperature. Streptavidin-RBD complexes were separated by size-exclusion chromatography using a Superdex 200 Increase 10/300 GL column (Cytiva) equilibrated in 20 mM sodium phosphate pH 7.2, 200 mM NaCl buffer.

Biotinylated S1 protein was obtained commercially (BioLegend). A 4X molar excess was incubated with streptavidin-Alexa Fluor 647 (0.5 mg/mL, BioLegend) for 30 min at room temperature directly before use.

### Synthesis of benzylguanine agarose

Benzylguanine agarose (BG-agarose) was synthesised by dissolving 500 mg low melting point agarose (#A5030, Sigma), 46 mg carbonyl diimidazole (Sigma) and 236 μL triethylamine (Sigma) in 50 mL anhydrous DMSO (Sigma) and stirring the solution at room temperature under argon atmosphere (adapted from^77^). After 1 h, 250 mg O-[4-(Aminomethyl)benzyl]guanine (Generon) was added and the solution was stirred overnight under argon. The solution was diluted with 450 mL warm distilled water and dialysed against water using snakeskin dialysis tubing (3,500 Da molecular weight cut-off, Thermo Fisher). After six water exchanges, the product was lyophilised yielding a white, spongy material.

### Determination of agarose functionalisation efficiency

BG-agarose (1.5 % (w/v)) was dissolved in PBS, melted by heating to 85 °C using a heated shaker and then cooled to 37 °C. BG-agarose beads (25 µm diameter) were generated by microfluidics using the same microfluidic chip as for cell encapsulation (flow rates: aqueous phase: 6 – 7 μL/min, oil phase: 16 – 18 μL/min). The emulsion was collected on ice to solidify the agarose. BG-agarose beads were demulsified by addition of 800 μL PBS and 200 μL 1H,1H,2H,2H-perfluoro-1-octanol (PFO, Alfa Aesar), filtered through a 50 μm strainer and washed once in 15 mL FACS buffer (1% ultra-low IgG FBS (Thermo Fisher) in PBS). Beads (200,000 per condition) were incubated with serial 1:2 dilutions of GFP-SNAP overnight at room temperature starting from 8.55 × 10^15^ GFP molecules per condition. Beads were washed twice in FACS buffer, and the GFP-fluorescence was analysed on an Attune Nxt flow cytometer (Thermo Fisher). Single beads were gated on SSC-A vs. FSC-A (log-scale) and FSC-H vs FSC-A (log scale). The mean GFP fluorescence and standard deviation for each condition was determined in FlowJo (version 10.7, BD Biosciences) and an unpaired t-test was performed in GraphPad Prism (version 6.00, GraphPad Software) to determine whether there is a significant difference between two conditions.

### Antibody capture on agarose beads

BG-agarose beads (1.5% (w/v) agarose, 25 μM diameter) were generated by microfluidics as described previously and functionalised with 50 µM anti-mouse κ VHH for 1 h at room temperature. After washing twice in FACS buffer, anti-streptavidin antibody (clone 3A20.2, mouse IgG2bκ, BioLegend) was added to 10^6^ beads at a concentration of 0.25 mg/mL and incubated for 1 h at room temperature. GFP-biotin was tetramerised by incubation with APC-streptavidin (BioLegend) for 1 h at 10X molar excess. The tetramer (428 kDa) was purified from excess GFP-biotin (25 kDa) with a 100 kDa cut-off Amicon Ultra centrifugal filter (Merck). Functionalised agarose beads were washed twice and then incubated with 20 nM of the GFP-streptavidin tetramers and anti-mouse IgG2a/b BV421 antibody (1:10 dilution, clone R2-40, BD Bioscience) for 30 min on ice. After two washing steps in FACS buffer, beads were analysed on an Attune Nxt flow cytometer (Thermo Fisher).

### Mouse immunisations

Mice of the C57BL/6 genetic background (8 – 14 weeks old) were immunised with EndoFit Ovalbumin (OVA) (Invivogen) or SARS-CoV-2 RBD. Antigens were prepared as 2 mg/mL sterile-filtered solutions in PBS, mixed with an equal volume of Alhydrogel adjuvant (Invivogen) and incubated on a tube roller for 30 min at 4 °C. Each mouse was injected intraperitoneally with 100 μL of 1 mg/mL antigen in Alum. For confocal experiments and RBD antibody generation, mice were additionally injected intraperitoneally with 100 μL of 0.5 mg/mL OVA or RBD in PBS 21 days post the first immunisation. Spleens were harvested 10 days after the first immunisation and bone marrow cells were harvested 21 days after the first or second immunisation.

### Extraction of splenocytes and bone marrow

Mice were sacrificed by cervical dislocation and subsequent cutting of a femoral artery. A single cell suspension of splenocytes was prepared by mashing the spleens through a 100 μm cell strainer into wash buffer (PBS, 0.1% FBS, 2 mM EDTA). Splenocytes were washed in ice-cold buffer and then filtered again through a 70 μm strainer. For bone marrow extraction, femur and tibias were removed from both legs, the edges of the bones were cut open with scissors and the bone marrow was flushed out using a 23-gauge needle and a syringe filled with wash buffer. A single cell suspension was prepared by filtering through a 100 μm cell strainer. Red blood cells of both spleen and bone marrow were lysed by incubation in 1 mL RBC lysis buffer (155 mM NH_4_Cl, 12 mM NaHCO_3_, 0.1 mM EDTA) for 5 min at room temperature. Cells were washed once in wash buffer and used for ASC enrichment. All centrifugation steps were carried out at 300 x g for 5 min at 4 °C.

### Mouse ASC enrichment

Plasma cells were enriched using direct magnetic labelling with anti-CD138 (syndecan-1) microbeads (Miltenyi Biotec) according to the manufacturer’s instructions. Enriched cells were washed once in wash buffer and an Fc blocking step was performed using TruStain FcX (anti-mouse CD16/32) antibody (BioLegend) according to the manufacturer’s instruction.

### Mouse ASC encapsulation

BG-agarose (3 – 6 % (w/v)) was dissolved in PBS, melted by heating to 85 °C using a heated shaker and then cooled to 37 °C. For FACS experiments following OVA immunisation, 6% BG-agarose was mixed with anti-mouse κ VHH-SNAP prior to encapsulation to obtain a 3% BG-agarose solution and a final VHH-SNAP concentration of 100 μM. BG-agarose can be functionalised with VHH-SNAP before or after encapsulation, but in our hands, VHH addition after hydrogel bead formation and demulsification lead to a slightly better signal-to-noise ratio. For all other experiments, a 3% (w/v) BG-agarose solution was therefore used directly without prior VHH-SNAP addition. ASCs obtained through magnetic selection (on average >2 × 10^6^ cells per spleen and >1.5 × 10^6^ cells for bone marrow) were washed, resuspended in encapsulation buffer (PBS, 10% ultra-low IgG FBS, 18% OptiPrep (Sigma Aldrich), 2 mM EDTA, 0.1% Pluronic F-68) at a concentration of 3 × 10^7^ cells/mL and filtered through a 40 μM strainer. Cells were then mixed with an equal volume of 3% BG-agarose in PBS. The solution was aspirated into microfluidic tubing and encapsulated with flow rates of around 6 – 7 μL/min for the aqueous phase (cells in agarose) and 16 – 18 μL/min for the oil phase (mean number of cells per droplet (λ) = 0.12). During encapsulation, the microfluidic chip was heated to 37 °C using a heated glass microscope plate (Bioscience Tools) although this is not strictly necessary as the low-melting point agarose will remain liquid for at least one hour. We routinely ran two microfluidic chips in parallel, enabling encapsulation of up to 1.26 × 10^7^ cells per hour. Alternatively, the mean number of cells per droplet (λ) can be increased up to a value of 0.3. After encapsulation, the emulsion was collected on ice to solidify the agarose. Hydrogel beads were demulsified by addition of 800 μL PBS and 200 μL PFO, filtered through a 50 μm strainer and washed once in 15 mL cold FACS buffer. If VHH-SNAP had not been added prior to encapsulation, the hydrogel beads were resuspended in 50 µM anti-mouse κ VHH-SNAP in PBS, incubated for 15 min at room temperature and then washed once in FACS buffer. For IgG secretion control experiments, no VHH-SNAP was added. Hydrogel beads were then resuspended in pre-warmed medium (RPMI supplemented with 10% ultra-low IgG FBS, 20 mM HEPES, 50 μM ß-mercaptoethanol, 1 mM sodium-pyruvate, 2 mM L-glutamine, 100 U penicillin/streptomycin) and incubated for 1 h 45 min at 37 °C, 5% CO_2_. All centrifugation steps were carried out at 300 x g for 10 min at 4 °C.

### Staining of encapsulated mouse ASCs for FACS

After incubation, hydrogel beads were washed once in FACS buffer. For OVA experiments, hydrogel beads were stained with anti-CD138 PE (1:80 dilution, clone 281-2, BioLegend), anti-FLAG PE-Cy7 (1:200 dilution, clone L5, BioLegend), OVA-Alexa Fluor 647 (3 μg/mL, Thermo Fisher), anti-IgG1 BV421 (1:25, clone RMG1-1, BioLegend), anti-IgG2a/2b BV421 (1:100, clone R2-40, BD), anti-IgG2a BV421 (also binds IgG2c, 1:80 dilution, clone RMG2a-62, BioLegend), anti-IgG3 BV421 (1:100 dilution, clone R40-82, BD), anti-IgM BV605 (1:40, clone RMM-1, BioLegend), anti-CD45R/B220 BV785 (1:50, clone RA3-6B2, BioLegend) for 30 min on ice and then washed once in FACS buffer. Shortly before acquisition, 5 μL of 7-AAD (50 µg/mL, BioLegend) or helix green (1 nM final concentration, BioLegend) were added to each tube. To simplify the panel, the anti-FLAG and anti-B220 antibodies were omitted in the RBD experiments. Hydrogel beads were stained with strep-RBD Alexa Fluor 647 (40 nM final concentration), anti-IgG1 BV421 (1:25, clone RMG1-1, BioLegend), anti-IgG2a/2b BV421 (1:100, clone R2-40, BD), anti-IgG2a BV421 (also binds IgG2c, 1:160 dilution, clone RMG2a-62, BioLegend), anti-IgG3 BV421 (1:100 dilution, clone R40-82, BD), anti-IgM BV605 (1:40, clone RMM-1, BioLegend), anti-CD138 BV785 (1:80, clone 281-2, BioLegend) for 30 min on ice and then washed once in FACS buffer. Shortly before acquisition, helix green (1 nM final, BioLegend) was added to each tube.

### Analysis of encapsulated mouse ASCs by confocal microscopy

Hydrogel beads were stained with OVA-Alexa Fluor 555 (1.5 μg/mL, Thermo Fisher), anti-CD138 Alexa Flour 647 (1:80 dilution, clone 281-2, BioLegend), donkey anti-mouse IgG Alexa Fluor Plus 405 (1:200 dilution, #A48257, Thermo Fisher) for 30 min on ice and then washed once in FACS buffer. For confocal imaging, 5 µL of encapsulated cells were pipetted into microscopy compatible 35 mm dishes (ibidi, #81156) and overlayed with 500 µL anti-evaporation oil (ibidi, # 50051). Imaging was performed on a Leica TCS SP8 confocal microscope. Fluorophores were excited with a 405 nm, a 552 nm, and a 638 nm laser, respectively. Images were exported and analysed with the open-source software Fiji^78^.

### Sorting of encapsulated mouse ASCs

Before the sort, hydrogel beads were filtered through a 50 μm strainer. Hydrogel beads were sorted on a four laser BD FACSAria III using a 100 μm nozzle into 96 well plates (Frame Star, 4titude) that contained 4 μL of lysis buffer (3 U/μL RNAseOUT RNase inhibitor, 1 mM DTT in 0.5X PBS^79^). Plates were spun down for 30 s at 400 x g immediately after the sort, placed on dry ice and then stored at -80 °C until processing. Hydrogel beads were gated to contain live ASCs (CD138^+^) that were IgG positive but IgM negative. The antigen signal (OVA or RBD) of hydrogel beads that were IgG^-^/IgM^-^ were used as guidance for setting the antigen positive gate. For OVA experiments, an additional anti-FLAG antibody was used to identify cells encapsulated into the functionalised BG-agarose hydrogel as the VHH-SNAP construct contained a FLAG-tag. For the OVA IgG secretion controls, cells in hydrogels beads were gated as live/FLAG^+^/CD138^+^ for the sample to which VHH-SNAP-FLAG was added and as live/CD138^+^ for the sample without VHH-SNAP-FLAG addition (therefore also FLAG-).

### PBMC isolation from peripheral blood

Peripheral blood (40 mL) was collected from two study participants in lithium heparin tubes. Peripheral blood mononuclear cells (PBMCs) were isolated by density gradient centrifugation using Histopaque-1077 (Sigma). Blood was diluted 1:1 with PBS (room temperature) and 35 mL of blood were layered onto 15 mL of Histopaque and centrifuged (acceleration: 1, deceleration: 0, 800 x g for 20 min at room temperature). The buffy coat containing the PBMCs was removed and the PBMCs were washed twice in ice cold wash buffer (0.1% FBS, 2 mM EDTA in PBS). PBMCs were counted, and one half was used directly for encapsulation experiments while the other half was resuspended in FBS and then mixed 1:1 with freezing medium (50% FBS, 10% DMSO in RPMI) by slow, dropwise addition. PBMCs were put into a freezing container (cooling rate: -1°C/minute) and stored at -80 °C before transfer to liquid nitrogen. Unless stated otherwise, all centrifugation steps were carried out at 400 x g for 10 min at 4 °C.

### PBMC stimulation

Normal cryopreserved human PBMCs (10 million cells/vial, Lonza, 4W-270) were thawed quickly at 37 °C and added dropwise to 10 mL of pre-warmed PBMC medium (RPMI 1640 supplemented with 10% FBS, 2 mM GlutaMAX, 1 mM sodium pyruvate, 10 mM HEPES pH 7.4, 1X MEM non-essential amino acids (Sigma), 50 μM ß-mercaptoethanol, 0.25 μg/mL amphotericin B, 100 U/mL penicillin/streptomycin) supplemented with 100 μg/mL DNase (Stemcell technologies). Cells were spun down at 500 x g for 10 min at 4 °C, resuspended in 2 mL of medium and rested for 1 h in the incubator (37 °C, 5% CO_2_). During this time, stimulation medium (2X concentration) was prepared by adding 400 ng/mL HA-tagged rhCD40L (R&D Systems) and 100 ng/mL anti-HA tag antibody (clone 543851, R&D Systems) to PBMC medium and incubating for 15 min at room temperature to crosslink the proteins. After incubation, 100 ng/mL human IL-21 (Peprotech) and 20 μg/mL anti-IgM antibody (polyclonal F(ab’)_2_ fragment goat anti-human, AB_2337553, Jackson ImmunoResearch) were added. After incubation, cells were washed once in MACS buffer and B cells were negatively selected with the human pan-B cell isolation kit (Miltenyi) according to the manufacturer’s instructions. Enriched B cells were spun down and resuspended in PBMC medium supplemented with 80 μg/mL apotransferrin (Sigma) at 10^6^ cells/mL. Cells were then mixed 1:1 (v/v) with stimulation medium and incubated for 96 h at 37 °C, 5% CO_2_.

### Encapsulation of stimulated PBMCs

A 6% (w/v) BG-agarose solution in PBS was heated to 85 °C using a heated shaker for 10 min and then cooled to 37 °C. The agarose was then mixed 1:1 (v/v) with anti-human κ VHH-SNAP (final concentration 100 μM) and left at 37 °C for 15 min before use. As for the mouse experiments, BG-agarose can be functionalised with VHH-SNAP before or after encapsulation, but in our hands, VHH addition after hydrogel bead formation and demulsification lead to a slightly better signal-to-noise ratio. After stimulation, cells were washed once in MACS buffer and CD38^+^ cells were positively selected using the CD38 MicroBead kit (Miltenyi) according to the manufacturer’s instructions. CD38^+^ cells were washed once and a blocking step using FcR Blocking Reagent (Miltenyi) was performed according to the manufacturer’s instructions. Cells were then resuspended in encapsulation buffer (PBS, 10% ultra-low IgG FBS, 18% Optiprep, 2 mM EDTA, 0.1% Pluronic F-68) and filtered through a 40 μM strainer to obtain a concentration of 3 × 10^7^ cells/mL. Cells were then mixed with an equal volume of VHH-functionalised 3% BG-agarose in PBS. Cells were encapsulated into agarose as previously described and the emulsion was collected on ice to solidify the agarose. Hydrogel beads were demulsified by addition of 800 μL PBS and 200 μL PFO. Beads were filtered through a 50 μm strainer and washed once in 15 mL cold FACS buffer. Afterwards, hydrogel beads were resuspended in pre-warmed PBMC medium (supplemented with 10% ultra-low IgG FBS instead of normal FBS) and incubated for 1 h 45 min at 37 °C, 5% CO_2_. All centrifugation steps were carried out at 300 x g for 10 min at 4 °C.

### Staining of encapsulated normal PBMCs for FACS and microscopy

After incubation, hydrogel beads were washed once in FACS buffer and then stained with anti-CD14 VioBlue (1:50 dilution, clone TÜK4, Miltenyi), anti-CD3 VioBlue (1:50 dilution, clone BW264/56, Miltenyi), anti-IgD VioBlue (1:50 dilution, clone IgD26, Miltenyi), anti-CD19 VioBright-FITC (1:50 dilution, clone LT19, Miltenyi), anti-CD27 APC-Vio770 (1:50 dilution, clone M-T271, Miltenyi), anti-CD20 PE-Vio770 (1:50 dilution, clone LT20, Miltenyi), anti-CD38 APC (1:50 dilution, clone IB6, Miltenyi), anti-IgG Fc PE (1:50 dilution, clone HP6017, BioLegend, discontinued), anti-IgM BV711 (1:20 dilution, clone MHM-88, BioLegend) and anti-FLAG Alexa Fluor 594 (1:200 dilution, clone L5, BioLegend). The sample was incubated on ice for 30 min and washed once in FACS buffer. DAPI (1:500,00) was added shortly before analysis on an Attune NxT flow cytometer. Hydrogel beads were examined for APC fluorescence (anti-CD38) and PE fluorescence (IgG secretion) under an EVOS FL (Thermo Fisher) using filters for Cy5 (ex: 628/40 nm, em: 692/40 nm) and RFP (ex: 531/40 nm, em: 593/40 nm), respectively.

### PBMC encapsulation and sort post vaccination

Blood was collected from study participants 7 – 9 days post the second BNT162b2 vaccine dose. Freshly isolated PBMCs were used directly for B cell enrichment. Cryopreserved PBMCs were thawed quickly at 37 °C and added dropwise to 10 mL of pre-warmed medium (RPMI 1640 supplemented with 10% ultra-low IgG FBS, 2 mM GlutaMAX, 1 mM sodium pyruvate, 10 mM HEPES pH 7.4, 100 U/mL penicillin/streptomycin) with 100 μg/mL DNase (Stemcell technologies). Cells were spun down (400 x g, 10 min, 4 °C) and washed once in MACS buffer. B cells were negatively enriched with the human pan-B cell isolation kit (Miltenyi) according to the manufacturer’s instructions. On average 5 – 8 × 10^6^ cells were obtained per purification. Enriched B cells were spun down and a blocking step using FcR Blocking Reagent (Miltenyi) was performed according to the manufacturer’s instructions. Cells were then resuspended in encapsulation buffer and filtered through a 40 μM strainer to obtain a concentration of 3 × 10^7^ cells/mL. Cells were then mixed with an equal volume of 3% BG-agarose in PBS and encapsulated and demulsified as previously described for mouse ASC encapsulation (λ = 0.12, two microfluidic chips run in parallel). Hydrogel beads were filtered through a 50 μm strainer and washed once in 15 mL cold FACS buffer. Beads were resuspended in PBS with anti-human VHH-SNAPs (anti-κ and anti-λ, 50 µM of each) and incubated for 15 min at room temperature. Hydrogel beads were then washed once in cold FACS buffer and resuspended in pre-warmed medium supplemented with 10 ng/mL human IL-6 followed by incubation for 1 h 45 min at 37 °C, 5% CO_2_.

### Staining and sorting of encapsulated post-vaccine PBMCs

For sorts with RBD, hydrogel beads were washed once in FACS buffer and stained using 40 nM strep-RBD Alexa Fluor 647, anti-IgG Fc PE (1:100 dilution, clone HP6017, BioLegend (discontinued)), anti-IgM BV785 (1:50 dilution, clone MHM-88, BioLegend), anti-CD38 BV421 (1:100 dilution, clone HB-7, BioLegend), anti-CD20 BV510 (1:50 dilution, clone 2H7, BioLegend) for 30 min on ice. Beads were washed once in FACS buffer and shortly before acquisition, helix green (1 nM final, BioLegend) was added to the sample. For sorts with RBD and S1, hydrogel beads were washed once in FACS buffer and then stained using 40 nM strep-S1 Alexa Fluor 647, 40 nM strep-RBD Alexa Fluor 488, anti-IgG Fc PE (1:100 dilution, clone HP6017, BioLegend (discontinued)), anti-IgA PE (1:50 dilution, clone IS11-8E10, Miltenyi), anti-IgM BV785 (1:50 dilution, clone MHM-88, BioLegend), anti-CD38 BV510 (1:50 dilution, clone HB-7, BioLegend), anti-CD20 PE-Cy7 (1:50 dilution, clone 2H7, BioLegend) for 30 min on ice. Hydrogel beads were washed once in FACS buffer and shortly before acquisition, DAPI (1:50,000 dilution, Thermo Fisher) was added to the sample.

Beads were (index) sorted on a four laser BD FACS Aria III using a 100 μm nozzle into 96 well plates (Frame Star, 4titude) that contained 6 μL of lysis buffer (5% PEG 8000, 0.1% Triton-X-100, 0.5 U/μL RNAseOUT, 0.5 μM OligoT30VN, 0.5 μM random hexamers, 0.5 mM/each dNTPs). The antigen signal of hydrogel beads that showed neither IgG/IgA nor IgM signal was used as guidance for setting the antigen-positive gate. Plates were spun down for 30 s at 400 x g immediately after the sort, placed on dry ice and then stored at -80 °C until processing.

### Reverse transcription, PCR & sequencing of mouse antibody variable regions

Plates were thawed on ice for 5 min and spun down at 400 x g for 1 min at 4 °C. For lysis and reverse transcription, a combination of previously published protocols^79,80^ was used together with the template switch oligo from the 10X Chromium Next GEM Single Cell V(D)J Reagent Kits v1.1 (without barcode and UMI). This protocol uses the terminal transferase activity of the Maxima H Minus reverse transcriptase to aid incorporation of a template switch oligo sequence into the nascent cDNA which can then serve as a universal forward primer binding site. Primer sequences can be found in **Supplementary Table 2**. Per well, 7 μL of lysis and annealing mix (0.6% IGEPAL, 5.5 mM random hexamer primers, 3.99 mM anchored oligo-dT, 0.9 U/μL RNaseOUT) were added and the plate was incubated at 65 °C for 5 min. Plates were placed on ice for 5 min and 10 μL of RT mix were added to each well (15% PEG 8000, 2X Maxima RT buffer, 2 mM dNTPs, 4 μM template-switch oligo (10X_TSO), 4 U/μL Maxima H Minus reverse transcriptase). Plates were quickly spun down and transferred to a thermocycler for reverse transcription (90 min at 42 °C, 5 cycles of 2 min at 50 °C followed by 2 min at 42 °C). Plates containing cDNA were stored at -20 °C. Leader peptide and variable regions of both heavy chain and κ light chain were then amplified in two rounds of semi-nested PCR. Reverse primer sequences from the 10X Chromium Next GEM Single Cell V(D)J Reagent Kits v1.1 were used. For the first PCR, 8 μL of cDNA reaction mix was added to 17 μL of PCR mix using KAPA HiFi HotStart polymerase and KAPA HiFi Fidelity Buffer according to the manufacturer’s instructions. The forward primer (0.3 μM final concentration, fw_TSO_PCR1) was designed to bind the PCR handle contained in the template switch oligo and reverse primers bound inside the constant regions of the immunoglobulins (final concentrations: 0.3 μM mIgG12abc, 0.1 μM mIgG2b, 0.1 μM mIgG3-1, 0.1 μM mIgG3-2, 0.2 μM mkappa). The PCR was performed at 95 °C for 3 min, followed by 20 cycles of 20 s at 98 °C, 30 s at 67 °C and 70 s at 72 °C and 5 min final elongation at 72 °C. From the first PCR reaction, 5 μL were transferred to the second PCR (final volume 25 μL) for 22 amplification cycles. For OVA experiments, the same PCR conditions and final subtype primer concentrations as for the first PCR were used. DNA obtained from anti-OVA hits were then cut out from agarose gels and amplicons from heavy and light chains were purified separately by gel extraction (Zymoclean Gel DNA Recovery Kit, Zymo Research).

For RBD experiments, PCRs were split after the first PCR and heavy and light chains were amplified separately using final primer concentrations of 0.3 μM for the reverse κ primer and the same conditions for the heavy chain primers as for the first PCR. Amplicons from anti-RBD hits were purified by SpeedBead (Sigma) purification. Following purification, all DNA sequences were obtained by Sanger sequencing. We found that sequencing efficiency was suboptimal with this protocol and recommend following a published protocol which does not employ a template switch oligo^79^. We also advise to decrease the amount of PFO for demulsification or use chemical-free breaking of the droplet emulsion using an antistatic gun as PFO can impact the efficiency of enzymatic reactions^81^.

### Reverse transcription, PCR & sequencing of human antibody variable regions

For human antibody sequencing, the reverse transcription in each well was performed based on a modified Smart-seq3 protocol using a template-switch approach^82^. Specifically, all volumes for the lysis and reverse transcription step were doubled, and random hexamers (0.5 μM final concentration) were added to the lysis buffer mix. All other steps were performed according to the published protocol. All oligonucleotides are listed in **Supplementary Table 3**.

For human anti-RBD antibodies, 12 μL of PCRI mix (final concentrations in KAPA HiFi Fidelity Buffer: 0.3 mM/each dNTPs, 0.5 mM MgCl_2_, 0.5 μM fw_Read1/2 primer and 0.1 μM of each reverse primer (rv_IgG_HC_PCR1, rv_kappa_PCR1, rv_lambda_PCR1), 0.02 U/μL KAPA HiFi HotStart polymerase) were added to each well after reverse transcription. PCR reactions were performed at 95 °C for 3 min, followed by 20 cycles of 20 s at 98 °C, 30 s at 65 °C and 2 min 30 s at 72 °C and a 5 min final elongation step at 72 °C. For the semi-nested second PCR, 5 μL were transferred from the first PCR and PCRs were performed using KAPA HiFi HotStart polymerase and KAPA HiFi Fidelity Buffer according to the manufacturer’s instructions. Because we initially planned to use Illumina sequencing^68^, we performed two sets of second PCRs, one that introduces the full-length Illumina Read 1 or Read 2 at the 5’ end and Read 2 or Read 1 at the 3’ end. For this reaction, fw_R1_PCR2 was combined with rv_R2 primers and fw_R2_PCR2 was combined with rv_R1 primers at 0.3 μM final concentration each in 25 µL. PCR reactions were performed as previously but with 22 amplification cycles. PCR products were purified using SpeedBeads (Sigma) and eluted with 15 μL of nuclease-free water. For addition of barcode indices and Illumina P5/P7 adapters, a third PCR was performed. For PCRs with 5’ Read 1, i5 primers (P5_1 - P5_96, used as well barcodes) were used as forward primers and i7 primers (P7_1 - P7_20, used as plate barcodes) were used as reverse primers. For PCRs with 5’ Read 2, i5 primers were used as reverse primers and i7 primers as forward primers. PCRs were performed using KAPA HiFi HotStart polymerase and KAPA HiFi Fidelity Buffer according to the manufacturer’s instruction in 25 μL volume with 0.3 μM of primers and 2 μL of PCR 2 as template with annealing at 65 °C and 22 cycles of amplification. PCR products were purified using SpeedBeads (Sigma) and sequenced by Sanger sequencing.

We found that this protocol did not yield high sequencing efficiencies (≈ 17%) and therefore decided to amplify the RBD/S1 hits with a different primer set which does not rely on template switching^83^. The reverse transcription in each well was performed in the same way as for the RBD sort (with template switching) but with PCR primers designed to bind inside the antibody sequence and the PCR reactions are therefore independent of template switching. After reverse transcription, the cDNA was diluted by addition of 27.4 µL of nuclease-free water. For the first PCR reaction, 4 µL of diluted cDNA were mixed with 21 µL of PCRI mix (final concentrations in KAPA HiFi Fidelity Buffer: 0.3 mM/each dNTPs, 0.5 mM MgCl2, 0.2 µM oPR_IGHV mix, 0.2 µM oPR_IGKV mix, 0.2 µM oPR_IGLV mix, 0.2 µM oPR_1st_rv mix, 0.02 U/μL KAPA HiFi HotStart polymerase). PCR reactions were performed at 95 °C for 3 min, followed by 50 cycles of 30 s at 98 °C, 30 s at 62 °C and 45 s at 72 °C and a 5 min final elongation step at 72 °C. Three separate, semi-nested second PCRs were performed for amplification of heavy chain and κ and λ light chains. For the second PCR, 4 µL of the first PCR was used directly and mixed with 21 µL of PCRII mix (final concentration in KAPA HiFi Fidelity Buffer: 0.3 mM/each dNTPs, 0.3 µM fw primer mix (oPR_IGHV fw mix, oPR_IGKV mix or oPR_IGLV mix) with the respective 0.3 µM rv primer mix (IgG/IgA rv mix, 3’ Cκ 494 (κ) or 3’ XhoI Cλ (λ)), 0.02 U/μL KAPA HiFi HotStart polymerase). PCR conditions were the same as for the first PCR. PCR products were purified using SpeedBeads (Sigma) and sequenced by Sanger sequencing, leading to an increase in matched heavy and light chain sequences (from 17% to over 40%). We advise to decrease the amount of PFO for demulsification or use chemical-free breaking of the droplet emulsion using an antistatic gun as PFO can impact the efficiency of enzymatic reactions^81^. Further optimisation of sequencing conditions is likely to increase the percentage of recovered sequences.

### Bioinformatic analysis

Antibody sequences were annotated using IMGT/V-Quest from the IMGT database^84^. Output files contain information such as gene segment usage, CDR and framework annotations and nucleotide and amino acid mutation statistics. All clonotyping was performed using an in-house script, with antibodies greedily clustered if they had a common IGHV gene assignment, the same length CDR3, and within 80% CDR3 amino acid identity of the cluster representative. The Coronavirus Antibody Database (CoV-AbDab^35^, February 2022 version), was downloaded as a set of anti-coronavirus reference antibodies with functional annotations. Clonotyping was then applied to functionally cluster input files containing each set of ASC-derived antibodies concatenated with all CoV-AbDab antibodies with a complete IGHV and CDRH3 label. An alternative clustering method based on grouping antibodies by similar predicted 3D structure (SPACE) was also applied as per the original paper^50^,using a threshold structural similarity of 0.75 Å. This was performed over each set of ASC-derived antibodies coupled with the subset of antibodies in CoV-AbDab with full variable heavy (VH) and variable light (VL) chain sequences (as sequence coverage over all six CDRs is required by SPACE). Statistical analysis was performed in Graph Pad Prism (version 6.00, GraphPad Software).

### Cloning of antibodies

For mouse antibodies, primers were designed to amplify the variable regions including the leader peptides with the forward primer binding to the leader peptide and the reverse primer binding in the constant region of the antibody chain. To generate PCR amplicons, Q5 Polymerase High-Fidelity 2X Master Mix (New England Biolabs) was used according to the manufacturer’s protocol using the second sequencing PCR as template. For anti-OVA antibodies, variable regions and leader peptides were cloned into pVITRO1 (Invivogen) plasmids that contained the constant regions for mouse IgG1 κ heavy and light chains using Gibson Assembly^75^. For anti-RBD antibodies, standard restriction cloning using the Type IIS restriction enzyme BsmBI-v2 (NEB) was employed to insert the leader peptides and variable regions into pVITRO1 (mouse IgG1 κ). Mouse antibodies were excluded from further testing if the plasmid sequencing did not match the expected sequence. For human antibodies, primers were designed to amplify the variable regions excluding the leader peptides. To generate PCR amplicons, Q5 Polymerase High-Fidelity 2X Master Mix (New England Biolabs) was used according to the manufacturer’s protocol using the second or third PCR as template. Variable regions were cloned into pVITRO1 plasmids that already contained the constant regions for human IgG1 heavy and light chains (κ or λ, Addgene plasmids #61883 and #50366)^85^ using Gibson Assembly^75^. Amino acid sequences of the variable regions of the antibodies can be found in **Supplementary Table 4** and **5**.

### Antibody expression and purification

FreeStyle 293-F cells were maintained in shaking flasks in Freestyle 293 Expression Medium at 0.4 – 2 × 10^6^ cells/mL (incubator conditions: 37 °C, 70% humidity, 8% CO_2_, 125 rpm). Per 10^6^ cells, 1.2 μg of DNA was mixed with 2.4 μL of 1 mg/mL polyethylenimine (linear PEI, M_W_ 25,000, Polysciences) in one tenth of the final culture volume of medium. The mixture was vortexed for 15 s followed by 15 min incubation at room temperature and then added to a culture of 10^6^ cells/mL. Valproic acid (VPA, 2-propyl-pentanoic acid, Sigma) was added 4 h post transfection to a final concentration of 3.5 mM. The culture supernatant was harvested one week after transfection by centrifugation for 15 min at 4000 x g, sterile filtered and stored at 4 °C until purification. The supernatant was added to a gravity column containing 0.5 mL of Pierce Protein G Agarose (Thermo Fisher) pre-equilibrated with PBS. The column was washed with ten column volumes of PBS and the antibodies were eluted in several fractions of 250 μL with elution buffer (10 mM glycine, pH 2.7) into collection tubes containing 12.5 μL of neutralisation buffer (1 M Tris-HCl, pH 9.0). The fractions containing protein (measured by absorbance at 280 nm) were pooled and the buffer was exchanged to PBS with PD-10 desalting columns (GE). Purity of the proteins was confirmed by SDS-PAGE. The recombinant antibodies were stored at 4 °C. Mouse antibodies were excluded from further testing if it was not possible to express the antibody in a single attempt.

### ELISA

Microtiter plates (384 well Nunc MaxiSorp, flat bottom, Thermo Fisher) were coated with 20 μL of 2 μg/mL RBD or S1 in PBS overnight at 4 °C. Plates were washed twice with 90 μL PBS-T and incubated for 1 h at room temperature with 40 μL of blocking buffer (2.5% BSA in PBS-T). Plates were washed twice with PBS-T. Antibody samples were diluted in 1% BSA in PBS-T (final concentrations 400 nM, 200 nM, 20 nM, 2 nM, 0.2 nM, 0.02 nM, 0.002 nM, 0.0002 nM) and 20 μL of sample was added per well followed by incubation at room temperature for 1 h. Commercial anti-RBD antibodies (mouse IgG1κ, clone 1035753, R&D systems or human IgG1κ, clone AM001414, BioLegend) and isotype controls (mouse IgG1κ, clone MG1-45, BioLegend or human IgG1κ, clone QA16A12, BioLegend) were used as controls. Plates were incubated for 1 h at room temperature, washed four times with PBS-T and 20 μL of polyclonal HRP-conjugated detection antibodies (goat anti-mouse, #G-21040, Thermo Fisher at 1:1,000 dilution or goat anti-human, #ab7153, Abcam at 1:10,000 dilution) were added to each well and incubated for 30 min (anti-mouse) or 45 min (anti-human) at room temperature. The plate was washed three times with PBS-T and one time with PBS. TMB (3,3’,5,5’-Tetramethylbenzidine, Thermo Fisher) solution (20 μL) was added per well and the enzymatic reaction was stopped by addition of 20 μL of 0.16M H_2_SO_4_ per well after 5 min. Absorbance was read at 450 nm and 540 nm in a plate reader (Infinite 200 PRO Tecan). For data analysis, the absorbance at 540 nm was subtracted from the signal at 450 nm. To obtain EC_50_s, titration curves were plotted as absorbance vs log (antibody concentration), then analysed by non-linear regression using the Sigmoidal, 5PL, X is log(concentration) function in GraphPad Prism^86^. For the purpose of visualisation, mouse antibodies with no quantifiable ELISA signal at the tested concentrations were assigned an arbitrary EC_50_ of 1,000 nM. Mouse antibodies that did show binding to RBD under these conditions were also tested in an anti-streptavidin ELISA (under the same conditions as described previously except that plates were coated with 20 μL of 2 μg/mL streptavidin). Under the tested conditions, no binding to streptavidin was observed.

### Determination of antibody binding affinities

Binding affinities were determined using the Octet RED96 biolayer interferometry system. Mouse antibodies (7.5 μg/mL anti-OVA and 5 μg/mL anti-RBD) were captured on pre-conditioned anti-mouse IgG Fc-capture biosensors (AMC, Sartorius) for 600 s. Human antibodies (5 μg/mL or 7.5 µg/mL) were captured on pre-conditioned anti-human IgG Fc-capture biosensors (AHC, Sartorius) for 600 s. For double-referencing, control sensors were not loaded with antibody but incubated in buffer instead. Biosensors were then probed with different concentrations of the corresponding antigens. All dilutions were performed in kinetics buffer (0.1% BSA, 0.02% Tween in PBS). Sensors were regenerated with 10 mM glycine pH 1.7. Data were analysed using the Octet data analysis software (version 11.0.0.4) using a 1:1 binding model. For the purpose of visualisation, one human antibody with no quantifiable K_D_ at the tested concentrations was assigned an arbitrary K_D_ value of 10^−7^ M. Individual binding curves of human RBD and S1 binders can be found in **Supplementary Figure 2**.

### Live virus neutralisation assays

The SARS-CoV-2 viruses used in this study were a wildtype (lineage B) isolate (SARS-CoV-2/human/Liverpool/REMRQ0001/2020), a kind gift from Ian Goodfellow (University of Cambridge), isolated by Lance Turtle (University of Liverpool) and David Matthews and Andrew Davidson (University of Bristol)^87,88^, and an Omicron (lineage B.1.1.529) variant, a kind gift from Ravindra Gupta^89^. Half-maximal inhibitory concentrations (IC_50_s) for indicated monoclonal antibodies were measured essentially as previously described^37–39^. In brief, luminescent reporter cells expressing SARS-CoV-2 Papain-like protease-activatable circularly permuted firefly luciferase (FFluc) were seeded in flat-bottomed 96-well plates. HEK293T-ACE2-30F-PLP2 cells (clone B7, available from the National Institute for Biological Standards and Control (NIBSC), catalogue number 101062) were used to test murine antibodies^37^, and A549-ACE2-TMPRSS2-30F-PLP2 cells (clone E8) were used to test human antibodies^89^. The next day, SARS-CoV-2 viral stock (MOI=0.01) was pre-incubated with a 3-fold dilution series of each antibody for 1 h at 37 °C, then added to the cells. 24 h post-infection, cells were lysed in Bright-Glo Luciferase Buffer (Promega) diluted 1:1 with PBS and 1% NP-40, and FFluc activity was measured by luminometry. Experiments were conducted in duplicate or triplicate, as indicated. To obtain IC_50_s, titration curves were plotted as FFluc vs log (antibody concentration), then analysed by non-linear regression using the Sigmoidal, 4PL, X is log(concentration) function in GraphPad Prism. IC_50_s were quantified when (1) at least 50% inhibition was observed at the highest antibody concentration tested (100 μg/mL), and (2) a sigmoidal curve with a good fit was generated. For purposes of visualisation and ranking, human antibodies with no quantifiable neutralising capacity were assigned an arbitrary IC_50_ of 1,000 μg/mL. Individual neutralisation curves for human antibodies can be found in **Supplementary Figure 3**.

### Epitope binning by biolayer interferometry

For in-tandem assays of mouse antibodies, biotinylated RBD (10 μg/mL) was immobilised on streptavidin biosensors (Sartorius) for 300 s. The biosensors were then probed with 100 nM of the first antibody for 600 s followed by 100 nM of the second antibody for 300 s. Independent binding of both clones was verified with a buffer control. For classical sandwich assays of human antibodies, 10 µg/mL of the first antibody was immobilised on anti-human IgG Fc-capture biosensors for 600 s. After blocking of free capture sites on the biosensors with an isotype control (10 µg/mL, human IgG1κ, clone QA16A12, BioLegend) for 600 s, sensors were incubated with 20 nM RBD for 300 s. Biosensors were then probed with the second antibody (10 µg/mL) for 300 s. All dilutions were performed in kinetics buffer (0.1% BSA, 0.02% Tween in PBS). Data were analysed using the Octet data analysis software (version 11.0.0.4). Individual binding curves of human sandwich assays can be found in **Supplementary Figure 4**.

### Purification of mRBD2 Fab and RBD-Fab complex

DNA encoding for the heavy and light chains of the Fab region of mRBD2 was cloned into a bi-cistronic expression cassette in the modified pExp vector. The light chain was under the control of a T7 promoter followed by a short intergenic region containing a favourable ribosomal binding site and the heavy chain with an added C-terminal His-tag. The pExp-Fab expression vector was co-transformed with plasmid pRARE2 (extracted from Rosetta2 (DE3) cells, Novagen) into Shuffle T7 Express cells (NEB). The cells were grown in 2xYT media supplemented with 100 µg/ml ampicillin and 15 µg/ml of chloramphenicol to OD_600_ 0.8 – 1.0 and then protein production was induced with 0.4 mM IPTG and proceeded for 16 h at 20 °C. The cells were harvested, resuspended in lysis buffer containing 20 mM HEPES-NaOH pH 7.2, 300 mM NaCl, 20 mM imidazole, supplemented with 250 µg/ml of DNaseI and 1 mM PMSF and then lysed by sonication. Lysate was cleared by centrifugation and protein was purified by Ni-affinity chromatography using PureCube Ni-NTA agarose resin. After washing the column with 20 CV of 20 mM HEPES-NaOH pH 7.2, 300 mM NaCl, 20 mM imidazole, the protein was eluted in 5 CV of 20 mM HEPES-NaOH pH 7.2, 300 mM NaCl, 250 mM imidazole. Next, the buffer of the eluted protein was exchanged to 20 mM HEPES-NaOH pH 7.2, 20 mM NaCl using a HiPrep 26/10 Desalting column (Cytiva) and the Fab was loaded onto HiTrap Capto SP ImpRes column (Cytiva). The protein was eluted in a linear gradient of 20 – 1000 mM NaCl buffer containing 20 mM HEPES-NaOH pH 7.2. Peak fractions were combined, the protein was concentrated using 30 kDa MWCO Amicon Ultra concentrators and further purified by size-exclusion chromatography using HiLoad Superdex 75 pg 16/60 column (Cytiva) equilibrated in 20 mM Tris-HCl pH 7.2, 250 mM NaCl. For the RBD-Fab complex formation, an excess of RBD was incubated with concentrated purified Fab for 1 h at 22 °C and then the complexes were separated by size exclusion chromatography using HiLoad Superdex 75 pg 16/60 column (Cytiva) equilibrated in 20 mM Tris-HCl pH 7.2, 250 mM NaCl. The RBD-Fab protein complex in peak fractions was concentrated to 10 mg/ml using 30 kDa MWCO Amicon Ultra concentrators and used for crystallization trials.

### RBD-mRBD2 Fab crystallisation and structure determination

Crystallisation of the RBD-Fab complex was performed in sitting drops set up using the Mosquito liquid dispensing robotic system (SPT Labtech) by mixing 200 nL of protein solution with 200 or 100 nL of precipitant in 96-well 2-drop MRC crystallization plates. The crystals appeared in several conditions of the Ligand-Friendly Screen (Molecular Dimensions) after 3 – 8 days at 16 °C, they were cryo-cooled in liquid nitrogen without the need for additional cryoprotectant. The crystals resulting in the presented structural data grew in the condition containing 100 mM Bis-Tris propane pH 6.5, 200 mM potassium thiocyanate, 20 % w/v PEG 3350, 10 % v/v ethylene glycol. X-ray diffraction data were collected at Diamond Light Source and processed by auto-processing pipelines using autoPROC and Staraniso (Global Phasing Ltd). The structure was solved using molecular replacement using structures of the RBD (PDB: 6YM0) and a Fab fragment (PDB: 3BGF) as search models. The structure was refined with phenix.refine^90^ and autoBuster (Global Phasing Ltd.) with manual fixing using Coot^91^. The final structure (PDB: 8BE1) contains two RBD:Fab complexes per asymmetric unit. As the complex made from chains D, E and F is likely affected by crystal packing (ca. 14° shift in the RBDs relative to the Fab, **Supplementary Figure 2**) we used the other complex for analysis. Crystallographic data collection, data reduction and refinement statistics are shown in **Supplementary Table 6**.

### Data analysis

Sanger sequencing reads were analysed using Geneious Prime (version 2019.2.1, Biomatters). Flow cytometry data were analysed using FlowJo (version 10.7, BD Biosciences). Statistical analyses were performed using GraphPad Prism (version 6.00, GraphPad Software).

