## Supplementary Information for "Microfluidics-enabled fluorescence-activated cell sorting of single pathogen-specific antibody secreting cells for the rapid discovery of monoclonal antibodies"

**Supplementary Figures**

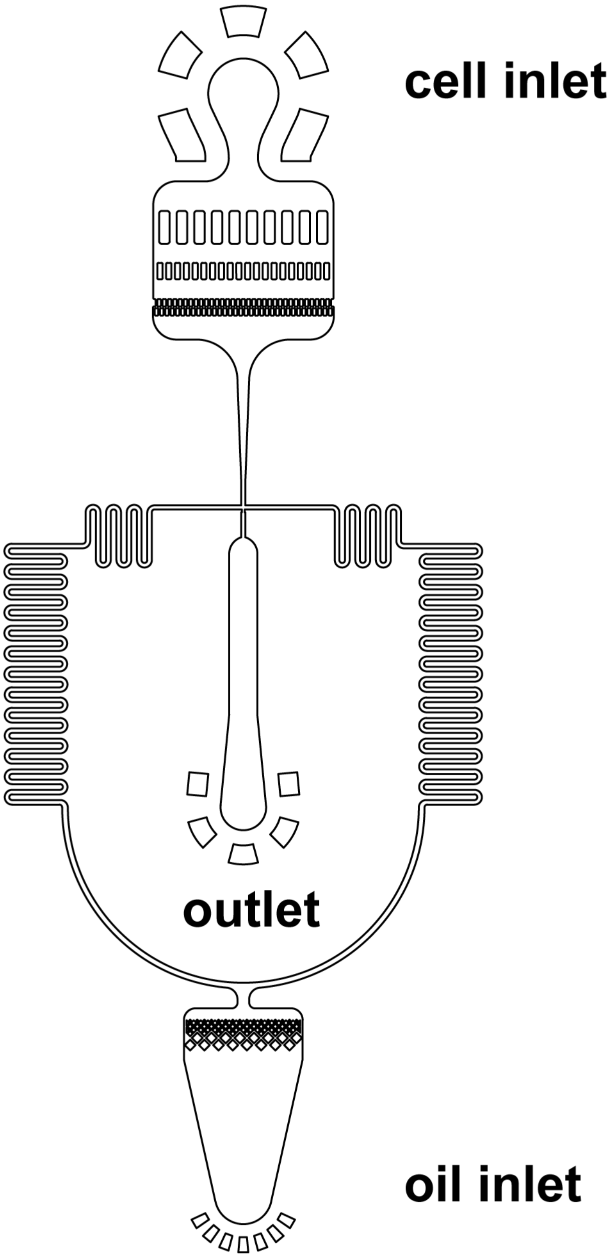

**Supplementary Figure 1 Design of microfluidic chip.** The channel layout is available as a .dxf file from https://openwetware.org/wiki/DropBase.

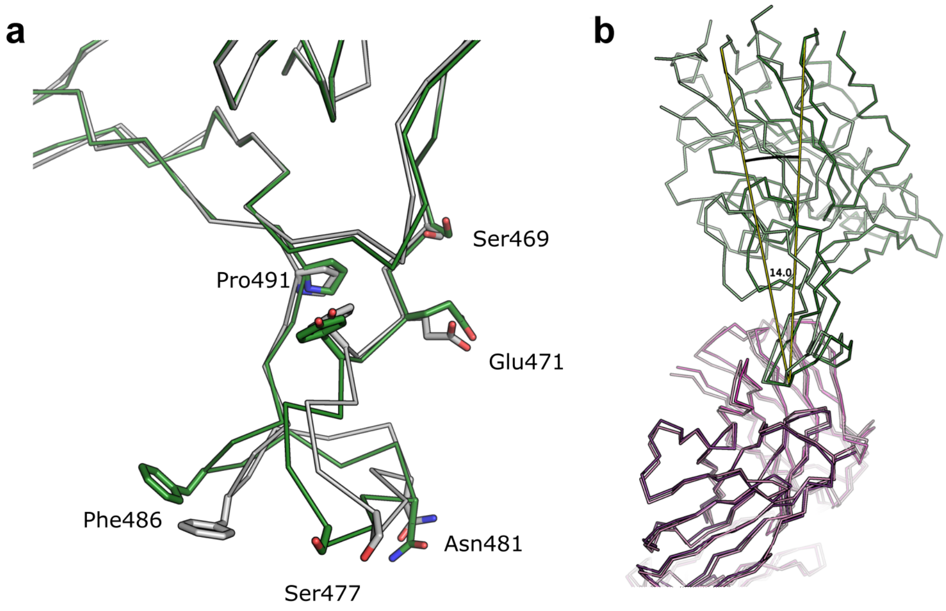

**Supplementary Figure 2 Conformational changes in RBD and in the RBD:Fab complexes.** (a) Superposition of RBD (chain C, green) from our Fab complex and RBD in an ACE2 complex (PDB: 6m0j, grey), showing how the loop is shifted by a few ångströms in the Fab complex with some of the sidechains from the moved loop shown as sticks. (b) Superposition of the two Fab fragments (dark and light purple) found in the asymmetric unit of our structure highlighting the circa 14° tilt of RBD relative to the Fab fragment in the second complex (RBD chain F, light green), in comparison with the other RBD (which is used in the analysis of the main paper; dark green), most likely due to crystal contacts.

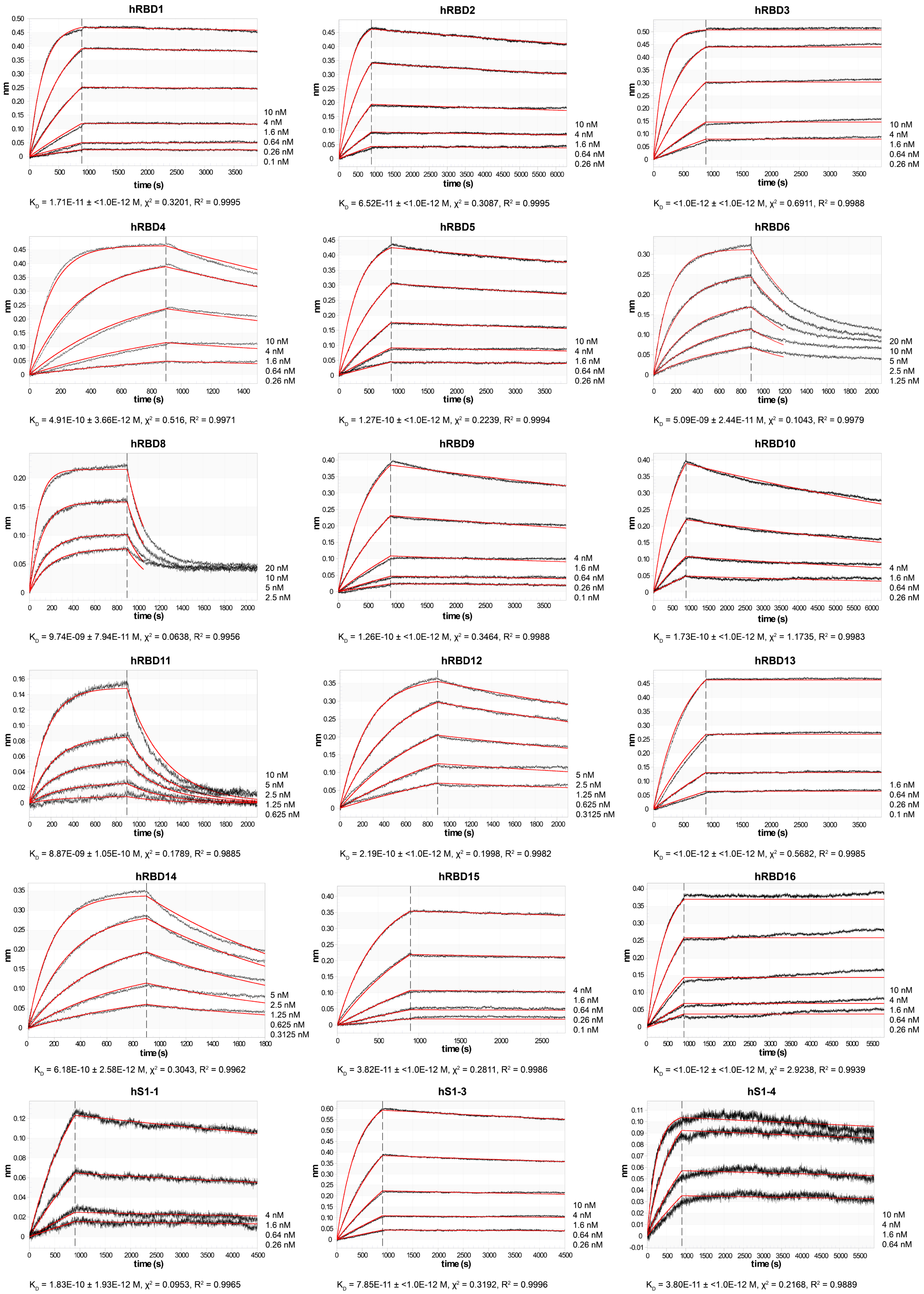

**Supplementary Figure 3 Determination of binding affinities of human anti-RBD and anti-S1 antibodies by biolayer interferometry.** The dissociation constant ($K_D$) as well as fitting parameters ($\chi^2$, $R^2$) are given below the respective sensorgrams.

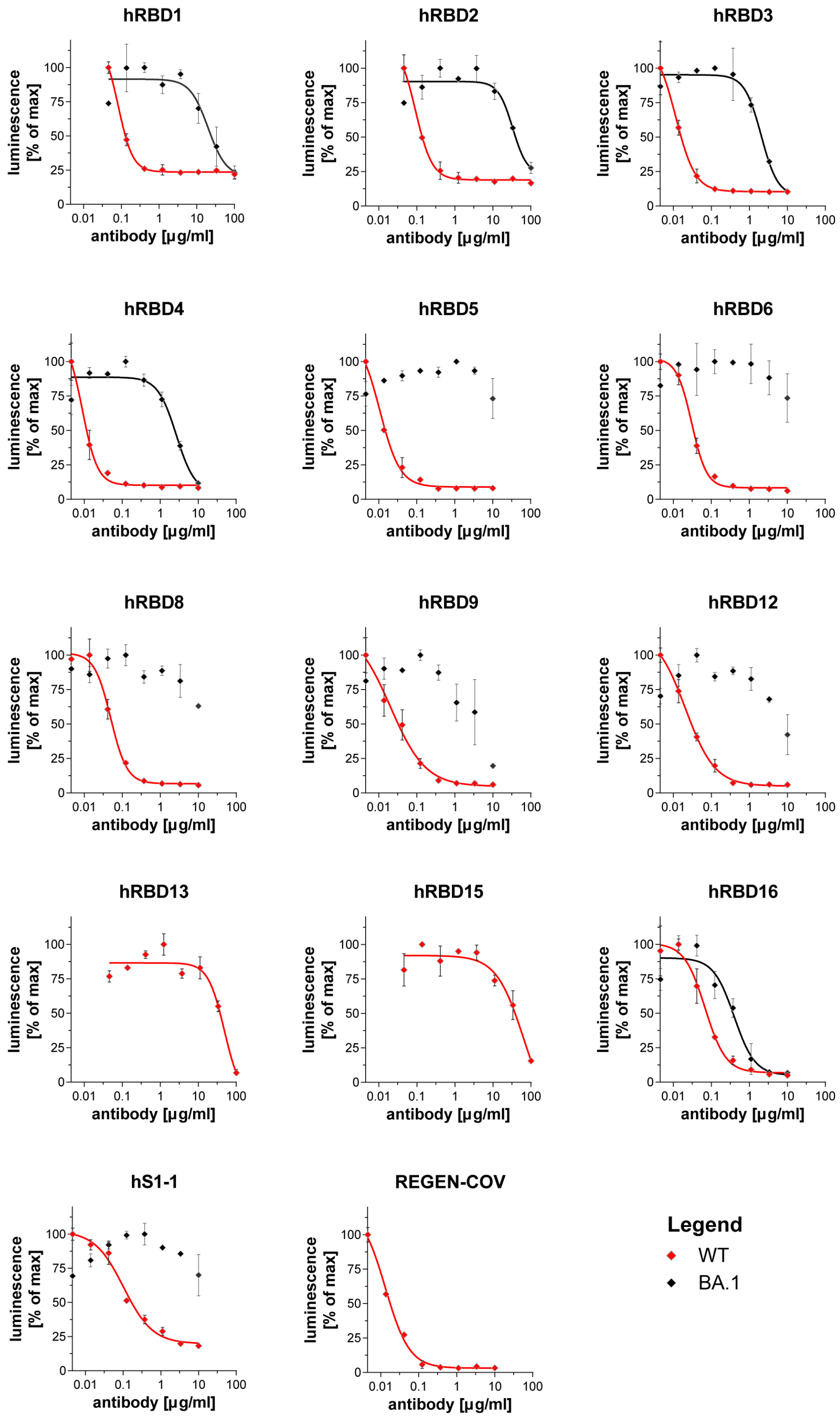

**Supplementary Figure 4 Neutralisation of SARS-CoV-2 by human antibodies.** Wildtype or Omicron BA.1 SARS-CoV-2 (MOI=0.01) was pre-incubated with a 3-fold dilution series of each antibody or the REGEN-COV (Ronapreve) antibody cocktail, then used to infect luminescent reporter cells. Levels of infection after 24 h were quantified as % of maximum luminescence. Mean values ± SD of two technical replicates are shown. $IC_{50}$s calculated from these titration curves are depicted in **Fig. 3e**. The titration curve for hS1-1 against WT SARS-CoV-2 is also shown in **Extended Data Fig. 7c**.

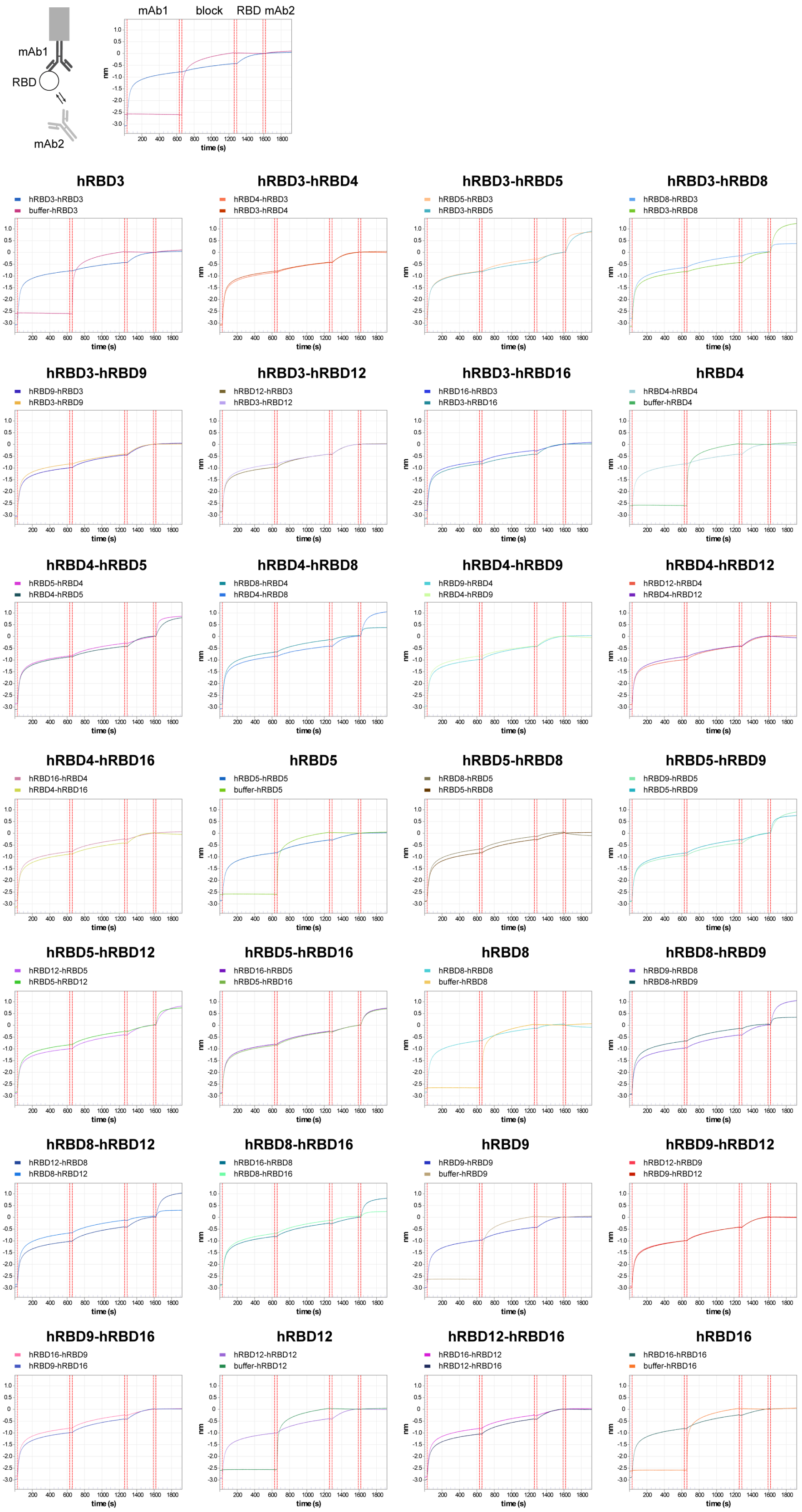

**Supplementary Figure 5 Epitope binning of selected human antibodies by biolayer interferometry.** Antibodies were immobilised on anti-human IgG Fc-capture biosensors followed by blocking of free capture sites on the biosensors with a human IgG1 isotype control and incubation with RBD. Biosensors were then probed with the second antibody.

### Supplementary tables

**Supplementary Table 1 Amino acid sequences of expressed proteins (VHH-SNAP, RBD)**

| name | sequence |
|---|---|
| **His-TwinStrep-SNAPf-XTEN-anti-mouse κ VHH (TP1170)-FLAG** | MQLSHHHHHHSSGLVPRGSHMASWSHPQFEKGGGSGGGSGGSAWSHPQFEKGSENLYFQSGGSMDKDCEMKRTTLDSPLGKLELSGCEQGLHRIIFLGKGTSAADAVEVPAPAAVLGGPEPLMQATAWLNAYFHQPEAIEEFPVPALHHPVFQQESFTRQVLWKLLKVVKFGEVISYSHLAALAGNPAATAAVKTALSGNPVPILIPCHRVVQGDLDVGGYEGGLAVKEWLLAHEGHRLGKPGLGSGSETPGTSESATPEGSQVQLVESGGGWVQPGGSLRLSCAASGFTFSDTAMMWVRQAPGKGREWVAAIDTGGGYTYYADSVKGRFTISRDNAKNTLYLQMNSLKPEDTARYYCAKTYSGNYYSNYTVANYGTTGRGTLVTVSSHMGLGSDYKDDDDK |
| **His-TwinStrep-SNAPf-XTEN-anti-human κ VHH-FLAG** | MQLSHHHHHHSSGLVPRGSHMASWSHPQFEKGGGSGGGSGGSAWSHPQFEKGSENLYFQSGGSMDKDCEMKRTTLDSPLGKLELSGCEQGLHRIIFLGKGTSAADAVEVPAPAAVLGGPEPLMQATAWLNAYFHQPEAIEEFPVPALHHPVFQQESFTRQVLWKLLKVVKFGEVISYSHLAALAGNPAATAAVKTALSGNPVPILIPCHRVVQGDLDVGGYEGGLAVKEWLLAHEGHRLGKPGLGSGSETPGTSESATPEGSQVQLQESGGGLVQPGGSLRLSCAASGRTISRYAMSWFRQAPGKEREFVAVARRSGDGAFYADSVQGRFTVSRDDAKNTVYLQMNSLKPEDTAVYYCAIDSDTFYSGSYDYWGQGTQVTVSSHMGLGSDYKDDDDK |
| **His-TwinStrep-SNAPf-XTEN-anti-human λ VHH-FLAG** | MQLSHHHHHHSSGLVPRGSHMASWSHPQFEKGGGSGGGSGGSAWSHPQFEKGSENLYFQSGGSMDKDCEMKRTTLDSPLGKLELSGCEQGLHRIIFLGKGTSAADAVEVPAPAAVLGGPEPLMQATAWLNAYFHQPEAIEEFPVPALHHPVFQQESFTRQVLWKLLKVVKFGEVISYSHLAALAGNPAATAAVKTALSGNPVPILIPCHRVVQGDLDVGGYEGGLAVKEWLLAHEGHRLGKPGLGSGSETPGTSESATPEGSQVQLQDSGGGLVQPGGSLRLSCAASGFTLDYYAIGWFRQAPGKEREGVSCISSSDGSSDGYTYYADSVKGRFTISRDNAKNTVYLQMNNLKPEDTAVYYCAARLFGVATTDPSYFDSWGQGTQVTVSSHMGLGSDYKDDDDK |
| **RBD-CHis** | MNITNLCPFGEVFNATRFASVYAWNRKRISNCVADYSVLYNSASFSTFKCYGVSPTKLNDLCFTNVYADSFVIRGDEVRQIAPGQTGKIADYNYKLPDDFTGCVIAWNSNNLDSKVGGNYNYLYRLFRKSNLKPFERDISTEIYQAGSTPCNGVEGFNCYFPLQSYGFQPTNGVGYQPYRVVVLSFELLHAPATVCGSSPHHHHHHHH |
| **RBD-Avi** | MNHHHHHHHHHSGTSGVDNKFNKERRRARREIRHLPNLNREQRRAFIRSLRDDPSQSANLLAEAKKLNDAQPKSGTENLYFQGSGTNLCPFGEVFNATRFASVYAWNRKRISNCVADYSVLYNSASFSTFKCYGVSPTKLNDLCFTNVYADSFVIRGDEVRQIAPGQTGKIADYNYKLPDDFTGCVIAWNSNNLDSKVGGNYNYLYRLFRKSNLKPFERDISTEIYQAGSTPCNGVEGFNCYFPLQSYGFQPTNGVGYQPYRVVVLSFELLHAPATVCGPGSSGLNDIFEAQKIEWHEA |

**Supplementary Table 2 Oligonucleotide sequences for RT-PCR of antibody variable genes from mouse.** The following modifications were used: riboguanosine (rG).

| name | sequence |
|---|---|
| **10X_TSO** | CTACACGACGCTCTTCCGATCTTTTCTTATATrGrGrG |
| **fw_TSO_PCR1** | AATGATACGGCGACCACCGAGATCTCTACACGACGCTCTTCC GATCT |
| **rv_mIgG12abc _PCR1** | GGACAGGGATCCAGAGTTCCA |
| **rv_mIgG2b_PCR1** | AGGTGACGGTCTGACTTGGC |
| **rv_mIgG3-1_PCR1** | GCTGGACAGGGCTCCATAGTT |
| **rv_mIgG3-2_PCR1** | GGCACCTTGTCCAATCATGTTCC |
| **rv_mkappa_PCR1** | ATGTCGTTCATACTCGTCCTTGGT |
| **fw_PCR2** | AATGATACGGCGACCACCGAGATC |
| **rv_mIgG12ac_PCR2** | CCCTTGACCAGGCATCC |
| **rv_mIgG2b_PCR2** | AGGTCACGGAGGAACCAGTTG |
| **rv_mIgG3-1_PCR2** | GGCATCCCAGTGTCACCGA |
| **rv_mIgG3-2_PCR2** | AGAAGATCCACTTCACCTTGAAC |
| **rv_mkappa_PCR2** | GAAGCACACGACTGAGGCAC |

**Supplementary Table 3 Oligonucleotide sequences for RT-PCR of antibody variable genes from human.** Indices (i5 and i7) from the Illumina xGen UDI Primer Pair list were used. For i5 primers, indices 1 – 96 were used and for i7 primers indices 1 – 20 were used. The following modifications were used: 5' biotin modification (5Biosg), phosphorothioated DNA bases (*), riboguanosine (rG).

| **name** | **sequence** |
|---|---|
| ***reverse transcription*** | |
| **Oligo dT** | 5Biosg-TTTTTTTTTTTTTTTTTTTTTTTTTTTTTTVN |
| **Smartseq3_N8_TSO** | 5Biosg-AGAGACAGATTGCGCAATGNNNNNNNNrGrGrG |
| ***PCRs RBD sort*** | |
| **fw_Read1/2** | AGATGTGTATAAGAGACAGATTGCGCAA*T*G |
| **rv_IgG_HC_PCR1** | TTGTCCACCTTGGTGTTGC*T*G |
| **rv_kappa_PCR1** | GTTTCTCGTAGTCTGCTTTGC*T*C |
| **rv_lambda_PCR1** | GGTGCTCCCTTCATGCG*T*G |
| **fw_R1_PCR2** | TCGTCGGCAGCGTCAGATGTGTATAAGAGACAGATTGCGC AATG |
| **rv_ R2_IgG_HC PCR2** | GTCTCGTGGGCTCGGAGATGTGTATAAGAGACAGAGTCCT TGACCAGGCAGCC |
| **rv_R2_kappa PCR2** | GTCTCGTGGGCTCGGAGATGTGTATAAGAGACAGAGATGG CGGGAAGATGAAGAC |
| **rv_R2_lambda PCR2** | GTCTCGTGGGCTCGGAGATGTGTATAAGAGACAGGGAGGG CGGGAACAGAGTGAC |
| **fw_R2_PCR2** | GTCTCGTGGGCTCGGAGATGTGTATAAGAGACAGATTGCG CAATG |
| **rv_R1_IgG_HC_PCR2** | TCGTCGGCAGCGTCAGATGTGTATAAGAGACAGAGTCCTT GACCAGGCAGCC |
| **rv_R1_kappa_PCR2** | TCGTCGGCAGCGTCAGATGTGTATAAGAGACAGAGATGGC GGGAAGATGAAGAC |
| **rv_R1_lambda_PCR2** | TCGTCGGCAGCGTCAGATGTGTATAAGAGACAGGGAGGGC GGGAACAGAGTGAC |
| **i5 primers**<br>P5-*i5 (example)*-anneal | AATGATACGGCGACCACCGAGATCTACAC-*ATATGCGC*-TCGTCGGCAGCGTCAGATG |
| **i7 primers**<br>P7-*i7 (example)*-anneal | CAAGCAGAAGACGGCATACGAGAT-*CTGATCGT*-GTCTCGTGGGCTCGGAGATG |

| *PCRs RBD/S1 sort* | |
|---|---|
| ***oPR_IGHV mix*** | |
| oPR-IGHV-1 | ATGGACTGGACCTGGAGCATCC |
| oPR-IGHV-2 | ATGGACTGGACCTGGAGGATCCTC |
| oPR-IGHV-3 | ATGGACTGGACCTGGAGGGTCTTC |
| oPR-IGHV-4 | ATGGACTGGATTTGGAGGGTCCTCTTC |
| oPR-IGHV-5 | ATGGACACACTTTGCTACACACTCCTGC |
| oPR-IGHV-6 | ACTTTGCTCCACGCTCCTGC |
| oPR-IGHV-7 | GGCTGAGCTGGGTTTTCCTTGTTG |
| oPR-IGHV-8 | GGCTCCGCTGGGTTTTCCTTGTTG |
| oPR-IGHV-9 | CACCTGTGGTTCTTCCTCCTGCTG |
| oPR-IGHV-10 | ATGAAACACCTGTGGTTCTTCCTCCTCC |
| oPR-IGHV-11 | ACATCTGTGGTTCTTCCTTCTCCTGGTG |
| oPR-IGHV-12 | GCCTCTCCACTTAAACCCAGGCTC |
| oPR-IGHV-13 | ATGTCTGTCTCCTTCCTCATCTTCCTGC |
| oPR-IGHV-14 | ATGGAGTTGGGGCTGAGCTGG |
| oPR-IGHV-15 | ATGGGGTCAACCGCCATCCTC |
| ***oPR_IGKV mix*** | |
| oPR-IGKV-1 | ATGAGGCTCCTTGCTCAGCTTCTGG |
| oPR-IGKV-2 | ATGGAAGCCCCAGCTCAGCTTC |
| oPR-IGKV-3 | CCCAGCTCAGCTTCTCTTCCTCCTG |
| oPR-IGKV-4 | TGGTGTTGCAGACCCAGGTCTTCATTTC |
| oPR-IGKV-5 | GTCCCAGGTTCACCTCCTCAGCTTC |
| oPR-IGKV-6 | GCCATCACAACTCATTGGGTTTCTGCTG |
| oPR-IGKV-7 | TCCCTGCTCAGCTCCTGGG |
| oPR-IGKV-8 | CCTGGGACTCCTGCTGCTCTG |
| ***oPR_IGLV mix*** | |
| oPR-IGLV-1 | CCCTGGGTCATGCTCCTCCTGAAATC |
| oPR-IGLV-2 | CTCTGCTGCTCCTCACTCTCCTCAC |
| oPR-IGLV-3 | ATGGCATGGATCCCTCTCTTCCTCG |
| oPR-IGLV-4 | CCTCTCTGGCTCACTCTCCTCACTC |
| oPR-IGLV-5 | ACACTCCTGCTCCCACTCCTCAAC |
| oPR-IGLV-6 | ATGGCCTGGATCCCTCTACTTCTCC |
| oPR-IGLV-7 | ATGGCCTGGGTCTCCTTCTACC |
| oPR-IGLV-8 | ATGGCCTGGACTCCTCTCTTTCTGTTC |
| oPR-IGLV-9 | ATGGCCTGGATGATGCTTCTCCTC |
| oPR-IGLV-10 | GTCCCCTCTCTTCCTCACCCTCATC |
| oPR-IGLV-11 | CTCCTCGCTCACTGCACAGG |
| oPR-IGLV-12 | CCTCTCCTCCTCACCCTCCTC |

| | |
|---|---|
| oPR-IGLV-13 | CTCCTCCTCACCCTCCTCACTC |
| oPR-IGLV-14 | ATGGCCTGGACCCCTCTCC |
| oPR-IGLV-15 | ATGGCCTGGACCCCACTCC |
| ***oPR_1st_rv mix*** | |
| Cg-RT | AGGTGTGCACGCCGCTGGTC |
| rv_IgA_HC_PCR1 | CAGGGCACAGTCACATCCT*G*G |
| 3' Cκ 543 | GTTTCTCGTAGTCTGCTTTGCTCA |
| 3' Cλ | CACCAGTGTGGCCTTGTTGGCTTG |
| ***IgG/IgA rv mix*** | |
| 3' IgG (internal) | GTTCGGGGAAGTAGTCCTTGAC |
| 3' IgA (internal) | GCGAYGACCACGTTCCCATC |
| ***reverse primers κ/λ (used separately)*** | |
| 3' Cκ 494 | GTGCTGTCCTTGCTGTCCTGCT |
| 3' XhoI Cλ | CTCCTCACTCGAGGGYGGGAACAGAGTG |

**Supplementary Table 4 Amino acid sequences of variable regions of expressed mouse antibodies.** Sequences marked with an asterisk (*) did not show antigen binding at the tested concentrations.

| name | sequence |
|---|---|
| **mOVA1_HC** | QVQLQQSGAELARPGASVKLSCKASGYTFTSYGISWVKQRTGQGLEWIGEIDPRSGNTYYNEKFKGKATLTADKSSSTAYMELRSLTSEDSAVYFCARGRAYWGQGTLVTVSA |
| **mOVA1_LC** | DIVMTQSPSSLSVSAGEKVTMSCKSSQSLLNSGNQKNYLAWYQQKPGQPPKLLIYGASTRESGVPDRFTGSGSGTDFTLTISSVQAEDLAVYYCQNDHSYPFTFGSGTKLEIK |
| **mOVA2_HC** | QVQLQQSDAELVKPGASVKISCKVSGYTFTDHKIHWMKQRPEEGLEWIGYIYPRDGSTKYNEKFKGKATLTADKSSSTAYMQLNSLTSEDSAVYFCARKRAGTWYFDVWGTGTTVTVSS |
| **mOVA2_LC** | NIVLTQSPASLAVSLGQRATISCRASESVDTYGNSFMHWYQQKPGQPPKLLIFLASNLESGVPARFSGSGSRTDFTLTIDPVEADDAATYYCQQNNEDPWTFGGGTKLEIK |
| **mRBD1_HC** | QVQLQQPGAELVKPGASVKVSCKASGYTFTSYWMHWVKQRPGQGLEWIGRIHPSDSDTNYNQKFKGKATLTVDKSSNTAYMQLSSLTSEDSAVYYCAILDCYWYFDVWGTGTTVTVSS |
| **mRBD1_LC** | QIVLTQSPAIMSASPGEKVTMTCSASSSVSYMYWYQQKPGSSPRLLIYDTSNLASGVPVRFSGSGSGTSYSLTISRMEAEDAATYYCQQWSSYPLTFGAGTKLELK |
| **mRBD2_HC** | QVQLQQSGAELARPGASVKLSCKASGYTFISNGISWVKQRTGQGLEWIGEIYPRNGNTYYNEKFKGKATLTADKSSSTAYMELRSLTSEDSAVFFCARDYYYGSFDYWGQGTTLTVSS |
| **mRBD2_LC** | DIQMTQSPSSLSASLGERVSLTCRASREISGYLTWLQQKPDGTIKRLIYAASTLDSGVPKRFSGSRSGSDYSLTISSLESEDFADYYCLQYASYPWTFGGGTKLEIK |
| **mRBD3_HC** | QVQLQQPGAELVMPGTSVKLSCKASGYIFTSYWMHWVKQRPGQGLEWIGEIDPSDNYTNYNQKFKGKSTLTVDKSSSTAYMQLSSLTSEDSAVYYCARGGLYYYGSSYDAMDYWGQGTSVTVSS |
| **mRBD3_LC** | DIVMTQAAPSVPVTPGESVSISCRSSKSLLHSNGNTYLYWFLQRPGQSPQLLIYRMSNLASGVPDRFSGSGSGTAFTLRISRVEAEDVGAYYCMQHLEYPFTFGSGTKLEIK |
| **mRBD4_HC** | QVQLQQPGAELVMPGASVKLSCKASGYTFTSYWMHWVKQRPGQGLEWIGEIDPSDSYTNYNQKFKDKSTLTVDKSSSTAYMQLSSLTSEDSAVYYCARGGVYYYGSSHYFDYWGQGTTLTVSS |
| **mRBD4_LC** | DIVMTQAAPSVPVTPGESVSISCRSSKSLLHSNGNTYLYWFLQRPGQSPQFLIYRMSNLASGVPDRFSGSGSGTAFTLRISRVEAEDVGVYYCMQHLEYPFTFGSGTKLEIK |
| **mRBD5_HC** | QVQLQQPGAELVKPGASVKVSCKASGYTFTSYWMHWVKQRPGQGLEWIGRIHPSDSDTNYNQKFKGKATLTVDKSSSTAYMQLSSLTSEDSAVYYCAIDWDGDYWGQGTTLTVSS |

| | |
|---|---|
| **mRBD5_LC** | DIVMSQSPSSLAVSAGEKVTMSCKSSQSLLNSRTRKNYLAWYQQKPGQS PKLLIYWASTRESGVPDRFTGSGSGTDFTLTISSVQAEDLAVYYCKQSYNL WTFGGGTKLEIK |
| **mRBD6_HC** | QVQLQQPGAELVKPGASVKVSCKASGYTFTSYWMHWVKQRPGQGLEWI GRIHPSDSDTNYNQKFKGKATLTVDKSSSTAYMQLSSLTSEDSAVYYCAM DWDVDYWGQGTTLTVSS |
| **mRBD6_LC** | DIVMSQSPSSLAVSAGEKVTMSCKSSQSLLNSRTRKNYLAWYQQKPGQS PKLLIYWASTRESGVPDRFTGSGSGTDFTLTISSVQAEDLAVYYCKQSYNL LTFGAGTKLELK |
| **mRBD7_HC** | QVQLKQSGAELVRPGASVKLSCKASGYTFTDYYINWVKQRPGQGLEWIA RIYPGSGNTYYNEKFKGKATLTAEKSSSTAYMQLSSLTSEDSAVYFCARY PLYDYDEEDYFDYWGQGTTLTVSS |
| **mRBD7_LC** | DIQMTQTTSSLSASLGDRVTISCRASQDISNYLNWYQQKPDGTVKLLIYYT SRLHSGVPSRFSGSGSGTDYSLTISNLEQEDIATYFCQQGNTLPYTFGGG TKLEIK |
| **mRBD8_HC*** | QVQLKQSGAELVRPGASVKLSCKASGYTFTDYYINWVKQRPGQGLEWIA RIYPGSGNTYYNEKFKGKATLTAEKSSSTAYMQVSSLTSEDSAVYFCARY PLYDYDEEDYFDYWGQGTTLTVSS |
| **mRBD8_LC*** | NIVLTQSPASLAVSLGQRATISCRASESVDSYGNSFMHWYQQKPGQPPKL LIYLASNLESGVPARFSGSGSRTDFTLTIDPVEADDAATYYCQQNNEDPW TFGGGTKLEIK |
| **mRBD9_HC*** | QVQLKQSGAELVRPGASVKLSCKASGYTFTDYYINWVKQRPGQGLEWIA RIYPGSGNTYYDEKFKGKATLTAEKSSSTAYMQLSSLTSEDSAVYFCARY PLYDYDEEDYFDYWGQGTTLTVSS |
| **mRBD9_LC*** | DIVMTQSQKFMSTSVGDRVSITCKASQNVRTAVAWYQQKPGQSPKPLIYL ASNRHTGVPDRFTASGSGTDFTLTISNVQSEDLADYFCLQHWNYPWTFG GGTKLEIK |
| **mRBD10_HC** | QVQLKQSGAELVRPGASVKLSCKASGYTFTDYYINWVKQRPGQGLEWIA RIYPGSGNTYYNEKFKGKATLTAEKSSSTAYMQLSSLTSEDSAVYFCARD YGSSYVDYFDYWGQGTTLTVSS |
| **mRBD10_LC** | DIQMTQTTSSLSASLGDRVTISCRASQDISNYLNWYQQKPDGTVKLLIYYT SRLHSGVPSRFSGSGSGTDYSLTISNLEQEDIATYFCQQGNTLPLTFGAGT KLELK |
| **mRBD11_HC** | QVQLQQSGAELARPGASVKLSCKASGYTFISNGISWVKQRTGQGLEWIG EIYPRNGNTYYNEKFKGKATLTADKSSSTAYMELRSLTSEDSAVFFCARD YYYGNFDYWGQGTTLTVSS |
| **mRBD11_LC** | NIVLTQSPASLAVSLGQRATISCRASESVDSYGNSFMHWYQQKPGQPPKL LIYLASNLESGVPARFSGSGSRTDFTLTIDPVEADDAATYYCQQNNEDPYT FGGGTKLEIK |
| **mRBD12_HC** | QVQLQQSGAELARPGASVKLSCKASGYTFTSYGISWVKQRTGQGLEWIG EIYPRSGNTYYNEKFKGKATLTADKSSSTAYMEVRSLTSEDSAVYFCARE GGYGYVMDYWGQGTSVTVSS |
| **mRBD12_LC** | DIQMTQTTSSLSASLGDRVTISCRASQDISNYLNWYQQKPDGTVKLLIYYT SRLHSGVPSRFSGSGSGTDYSFTISNLEQEDIATYFCQQGNTLPWTFGGG TKLEIK |

**mRBD13_HC*** EVKLVESGGGLVQPGGSLSLSCAASGFTFTDYYMTWVRQPPGKALEWLG FIRNKANGFTTESSASVKGRFTISRDNSQSILYLQMNALRAEDSATYYCAR YIRTGWYFDVWGTGTTVTVSS

**mRBD13_LC*** NIVLTQSPASLAVSLGQRATISCRASESVDSYGNSFMHWYQQKPGQPPKL LIYLASNLESGVPARFSGSGSRTDFTLTIDPVEADDAATYYCQQNNEDPYT FGGGTKLEIK

**mRBD14_HC** EVKLVESGGGLVQPGGSLSLSCAASGFTFTDYYMSWVRQPPGKALEWL GFIRNKANGYTTEYSASVKGRFTISRDNSQSILYLQMNALRAEDSATYYCA RYIRTGWYFDVWGTGTTVTVSS

**mRBD14_LC** DIVMTQSQKFMSTSVRDRVSITCKASQNVRTAVAWYQQKPGQSPKALIYL ASNRHTGVPDRFTGSGSGTDFTLTISNVQSEDLADYFCLQHWNYPWTFG GGTKLEIK

**mRBD15_HC** QIQLVQSGPDLKKPGETVKISCKASGYTFTEYPMHWVKQAPGKGFQWMG MIYTDTGEPTYAEEFKGRFAFSLETSASTAYLQINNLKNEDTSTYFCVRAG PSYAMDYWGQGTSVTVSS

**mRBD15_LC** NIVLTQSPASLAVSLGQRATISCRASESVDSYGNSFMHWYQQKPGQPPKL LIYLASNVESGVPARFSGSGSRTDFTLTIDPVEADDAATYYCHQINEDPYT FGGGTKLEIK

**mRBD16_HC** QIQLVQSGPELKKPGETVKISCKASGYTLTEYPMHWVKQAPGKGFKWMG MIYTDTGEPTYAEEFKGRFAFSLETSASTAYLQINNLKNEDTATYFCVRAG PSFAMDYWGQGTSVTVSS

**mRBD16_LC** NIVLTQSPASLAVSLGQRATISCRASESVDSYGNSFMHWYQQKPGQPPKL LIYLASNLESGVPARFSGSGSRTDFTLTIDPVEADDAATYYCQQNNEDPYT FGGGTKLEIK

**mRBD17_HC** QIQLVQSGPELKKPGETVKISCKASGYTLTEYPMHWVKQAPGKGFKWMG MIYTDTGEPTYAEEFKGRFAFSLETSASTAYLQINNLKNEDTATYFCVRAG PSFAMDYWGQGTSVTVSS

**mRBD17_LC** DIQMTQTTSSLSASLGDRVTISCRASQDISNYLNWYQQKPDGTVKLLIYYT SRLHSGVPSRFSGSGSGTDYSLTISNLEQEDFATYFCQQGNTLPYTFGGG TKLEIK

**mRBD18_HC** QIQLVQSGPELKKPGETVKISCKASGYTFTDYPMHWVQQAPGKGFKWMG MIYTDTGEPTYAEEFKGRFAFSLETSASTAYLQINNLKNEDTATYFCVRWA GYFDYWGQGTTLTVSS

**mRBD18_LC** DIVLTQSPASLAVSLGQRATISCRASESVDSYGNSFMDWYQQKPGQPPKL LIYRASNLESGIPARFSGSGSRTDFTLIINPVEADDVATYYCQQSNEHPLTF GAGTKLELK

**mRBD19_HC** QIQLVQSGPELKKPGETVKISCKASGYTFTEYPMHWVKQAPGKGFKWMG MIYTDTGEPTYAEEFKGRFAFSLETSASTAYLQINNLKNEDTATYFCVRGG PYYAMDYWGQGTSVTVSS

**mRBD19_LC** NIVLTQSPASLAVSLGQRATISCRASESVDSYGNSFMHWYQQKPGQPPKL LIYLASNLESGVPARFSGSGSRTDFTLTIDPVEADDAATYYCQQNNEDPW TFGGGTKLEIK

**mRBD20_HC** QIQLVQSGPELKKPGETVKISCKASGYTLTEYPMHWVKQAPGKGFKWMG MIYTDTGEPTYAEEFKGRFAFSLETSASTAYLQINNLKNEDTATYFCVRAG PSYAMDYWGQGTSVTVSS

| | |
|---|---|
| **mRBD20_LC** | NIVLTQSPASLAMSLGQRATISCRASESVDSYGNSFMHWYQLKPGQPPKL LIYLASNLESGVPARFSGSGSRTDFTLTIDPVEADDAATYYCQQNNEDPYT FGGGTKLEIK |
| **mRBD21_HC** | QIQLVQSGPELKKPGETVKISCKASGYTFTKYPMHWVKQAPGKGFKWMG MIYTDTGEPTYAEEFKGRCAFSLETSASTAYLQINNLKNEDTATYFCVRWG GSYALDYWGQGTSVTVSS |
| **mRBD21_LC** | DIVLTQSPASLAVSLGQRATISCRASESVDSYGNSFMHWYQQKPGQPPKL LIYRASNLESGIPARFSGSGSRTDFTLTINPVEADDVATYYCQQSNEDPLT FGAGTKLELK |
| **mRBD22_HC** | QIQLVQSGPELKKPGETVKISCKASGYTFTEYPMHWVKQAPGKGFKWMG MIYTDTGEPTYAEEFKGRFAFSLETSASTAYLQINNLKNEDTATYFCVRGG YYYFDYWGQGTTLTVSS |
| **mRBD22_LC** | NIVLTQSPASLAVSLGQRATISCRASESVDSYGNSFMHWYQQKPGQPPKL LIYLASNLESGVPARFSGSGSRTDFTLTIDPVEADDAATYYCQQNNEDPW TFGGGTKLEIK |
| **mRBD23_HC** | QVQLQQPGAELVRPGSSVKLSCKASGYTFTSYWMHWVKQRPIQGLEWIG NIDPSDSETHYNQKFKDKATLTVDKSSSTAYMQLSSLTSEDSAVYYCTRP GSLRSRGWYFDVWGTGTTVTVSS |
| **mRBD23_LC** | DIVMTQAAPSVPVTPGESVSISCRSSKSLLHSNGNTYLYWFLQRPGQSPQ LLIYRMSNLASGVPDRFSGSGSGTAFTLRISRVEAEDVGVYYCMQHLEYP YTFGGGTKLEIK |
| **mRBD24_HC** | EVQLQQSGPDLVKPGASVKISCKASGYTFTDYYMNWVKQSHGKSLEWIG DINPNNGGTYYNQKFKGKATLTVDKSSSTAYMELRSLTSEDSAVYYCARQ LRRWGQGTLVTVSA |
| **mRBD24_LC** | DIVLTQSPASLAVSLGQRATISCKASQSVDYDGDSYMNWYQQKPGQPPK LLIYAASNLESGIPARFSGSGSGTDFTLNIHPVEEEDAATYYCQQSNEDPR TFGGGTKLEIK |
| **mRBD25_HC** | QVQLQQSGPELVKPGASVKISCKASGYAFSNSWMNWVKQRPGKGLEWI GRIYPGDGDTNYNGKFKGKATLTADKSSSTAYMQLSSLTSEDSAVYFCAN YYGDYWGQGTTLTVSS |
| **mRBD25_LC** | QIVLTQSPALMSASPGEKVTMTCSASSSVSYMYWYQQKPRSSPKPWIYLT SNLASGVPARFSGSGSGTSYSLTISSMEAEDAATYYCQQWSSNPYTFGG GTKLEIK |
| **mRBD26_HC*** | QVQLQQSGAELVRPGASVKMSCKASGYTFTTYPIEWMKHNHGKSLEWIG NFHPFNDDTNYNENFKDKATLTVEKSSNTVYLELSRLTSDDSAVYYCARG GLLRPMDYWGQGTSVTVSS |
| **mRBD26_LC*** | DIVMSQSPSSLAVSVGEKVTMSCKSSQSLLYSSNQKNYLAWYQQKPGQS PKLLIYWASTRESGVPDRFTGSGSGTDFTLTISSEKDEDLAVYYCQQYYSY PFTFGSGTKLEIK |

**Supplementary Table 5 Amino acid sequences of variable regions of expressed human antibodies.** Sequences marked with an asterisk (*) did not show antigen binding at the tested concentrations.

| name | sequence |
|---|---|
| **hRBD1_HC** | QVQLVQSGSELKKPGASVKVSCKASGNTLTNHALNWVRQAPGQGLEWMG WINTNTGIPTYAQGFTGRFVFSLDTSVSTAYLQISSLKAEDTAVYYCARVGQ TAIAALDDAFDIWGQGTMVTVSS |
| **hRBD1_LC** | DIVMTQSPDSLAVSLGERATINCKSSQSVLYNSNNKNFLAWYQQKPGQPPK LLIYWASTRESGVPDRFSGSGSGTDFTLTISSLQAEDVAVYYCQQYYSTPLT FGGGTKVGIK |
| **hRBD2_HC** | QVQLVQSGSELKKPGASVKVSCKASGYTFSNYAINWVRQAPGQGLEWMG WINTNTGHPTYAQGFTGRFVFSLDASVTTAYLQISSLKAEDTAVYYCARVG RTAIAALDDAFDIWGQGTMVTVSS |
| **hRBD2_LC** | DIVMTQSPDSLAVSLGERATINCKSSQSVLYSSNNKNFLAWYQQKPGQPPK LLINWASTRKSGVPDRFTGSGSGTDFTLTISSLQAEDVAVYFCQQYYSTPLT FGGGTKVEIK |
| **hRBD3_HC** | QLQLQESGPRLVRPSETLSLTCTVSGGSINSGTHYCGWIRQSPGKGLEWIG SIYYSGTTYYNPSLKSRVTISVDTSKNQFSLKLSSVTAADTAVYFCARRRAG SYFKDLSDYWGQGTLVTVSS |
| **hRBD3_LC** | NFMLTQPHSVSESPGKTVTISCTRSSGSIASNYVQWYQQRPGGAPTTVIYE DNQRPSGVPDRFSGSIDTSSNSASLTISGLKTDDEADYYCQSYDSTNHEVF GGGTKLTVL |
| **hRBD4_HC** | QLQLQESGPGLVKPSETLSLTCTVSGGSISTSSHYCAWIRQPPGKGLEWIG SIYYRGSTYYNPSLKSRVTISVDTSKNHFSLKLSSVTAADTAVYYCARRRAG SYYKDLFDYWGQGTLVTVSS |
| **hRBD4_LC** | NFMLTQPHSVSESPGKTVTISCTRSSGSIASNYVQWYQQRPGSAPTTVIYE DNQRPSGVPDRFSGSIDSSSNSASLTISGLKTEDEADYYCQSYDNSNHEVF GGGTKLTVL |
| **hRBD5_HC** | QVQLQQWGAGLLQPSETLSLTCAVYGGSFSDYYWSWIRQPPGRGLEWIG EIDHSGSTNYNPSLKSRVTILVDTSKNQFSLKLSSVTAADTAVYYCARAHLIG HCGDGGCYSGPDPSSWFDPWGQGTLVIVSS |
| **hRBD5_LC** | SSELTQDPAVSVALGQTVRITCQGDSLRRYFASWYQQKPGQAPVLVIYGKN IRPSGIPDRFSGSSSGNTGSLTITGAQAEDEADYYCHSRDSSGHHPLFGGG TKLTVL |
| **hRBD6_HC** | QVQLQQWGAGLLKPSETLSLSCAVYGGAFSDYYWSWIRQPPGKGLEWIG EIDLSGSTNYNPSLKSRVTISVDTSKNQFSLKLISVTAADTAVYYCARTHLIG YCSGGSCYSGPDPSNWFDPWGQGTLVTVSS |
| **hRBD6_LC** | SSELTQDPAVSVALGQTVRITCQGDSLRSYYASWYQQKPGQAPVLVIYGKN NRPSGIPDRFSGSSSGNTASLTITGAQAEDEADYYCNSRDTSGHHPVFGG GTKLTVL |
| **hRBD7_HC*** | QVQLQQWGAGLLKPSETLSLTCAVYGGSFSGYYWSWIRQPPGKGLEWIG EINHSGSTNYNPSLKSRVTISLDTSKNQFSLTLTSVTAADTAVYSCARGHLIG YCSGGSCYSGPDPSNWFDPWGQGTLVTVSS |

| | |
|---|---|
| **hRBD7_LC*** | DIVLTQSPGTLSLSPGERATLSCRASQSVSNYLAWYQQKPGQAPRLLIYGASSRATGIPDRFSGSGSETDFTLTISRLEPEDFAVYYCQQYGRSSRTFGQGTKLEIK |
| **hRBD8_HC** | QVQLQQWGAGRLKPSETLSLTCAVYGGSFSGYYWSWIRQPPGKGLEWIGEIDHSGSTNYNSSLKSRVTISVDTSKKWFSLKLSSVTAADTAVYYCARAHLIGDCGGGRCYSGPDPSNWFDPWGQGTLVTVSS |
| **hRBD8_LC** | SSELTQDPAVSVALGQTVRITCQGDSLRSYYASWYQQKPGQAPILVIYGKDNRPSGIPDRFSGSSSENTASLTITGAQAEDEADYYCNCRDTSGNHPVFGGGTKLTVL |
| **hRBD9_HC** | EVQLVESGGGLIQPGGSLRLSCAASGLTVSSNYMNWVRQAPGKGLEWVSVLYSGGSTYYADSVKGRFTISRDNSKNTLYLQMNSLRAEDTAVYYCARDLVAYGMDVWGQGTTVTVSS |
| **hRBD9_LC** | DIQLTQSPSFLSASVGDRVTITCRASQGISSDLAWYQQKPGKAPKLLIYAASTLQSGVPSRFSGSGSGTEFTLTISSLQPEDFATYYCQQLNSYPTFGQGTRLEIK |
| **hRBD10_HC** | EVQLVESGGGLVQPGGSLRLSCSASGFTFTTYNMNWVRQAPGKGLEWVSYISSSSSTIYYADSVEGRFTISRDNAKNSLYLQMNSLRDEDTAVYYCARGVGATSYYYHGMDVWGQGTTVSVSS |
| **hRBD10_LC** | DIQMTQSPSSLSASVGDRVTITCRASQSIYSYLNWYKQKAGKAPKLLIYAASSLQSGVPSRFSGSGSGTDFTLTIISLQPEDFATYYCQQSYSTPPTFGQGTKVEIK |
| **hRBD11_HC** | QVQLQESGPGLVKPSQTLSLTCTVSGGSISSGGYYWSWIRQHPGKGLEWIGYIFYSGSTYYNPSLKSRVIISVDTSKNQFSLKLSSVTAADTAAYYCARVNRVYGSGSYSLDPWGQGTLVTVSS |
| **hRBD11_LC** | QSALTQPASVSGSPGQSITISCTGTSRDVGDYDYVSWYQQHPGKAPKLMIYEGSERPSGVSNRFSGSKSGNTASLTISGLQAEDEADYYCCSYTGSSTSFLFGTGTKVTVL |
| **hRBD12_HC** | EVQLVESGGGLVQPGGSLRLSCAASGLTVSSNYMSWVRQAPGKGLEWVSVIYSGGSTYYADSVKGRFTISRDNSKNTLYLLMNSLRAEDTAVYYCARDLQQLGGMDVWGQGTTVTVSS |
| **hRBD12_LC** | DIQMTQSPSSLSASVGDRVTITCQASQDISNYLNWYQQKPGKAPKLLIYDASNLETGVPSRFSGSGSGTDFTFTISSLQPEDIATYYCQQYDNLPQSFGGGTKVEIK |
| **hRBD13_HC** | QVQLVESGGGVVQPGRSLRLSCAVSGFTFRNYGMDWVRQAPGKGLEWVAVISYDGSNKYYADSVKGRFTISRDNSKNTLFLHLNSLRAEDTAVYYCAKAGGGPYCGGGNCYMHYFDYWGQGTQVTVSS |
| **hRBD13_LC** | DIQMTQSPSSLSASVGDRVTITCQASQDINNYLNWYQQKPGKAPKLLIYDASNLETGVPLRFSGSGSATHFSFTISSLQPEDIATYYCQQYDHLPCTFGQGTKLEIR |
| **hRBD14_HC** | EVQLVESGGGLVQPGGSLRLSCAASGFSFSSSDMHWVRQATGKGLEWVSGIGTSGDTYYLGSVKGRFTISRDDAKNSFYLQMNSLRAGDTAVYYCARGEMAATGLWYYFDYWGQGTLVTVSS |
| **hRBD14_LC** | DIQMTQSPSSLSASVGDRVTITCRASQSISSYLNWYQQKPGKAPKLLIYAASSLQSGVPLRFSGSGSGTDFTLTITSLQPEDFATYYCQQSYSTPPWTFGQGTKVEIK |

| | |
|---|---|
| **hRBD15_HC** | QVQLVQSGAEVKKPGASVKVSCKASGYTFTSQYMHWVRQAPGQGLEWMGIINPSGGSTSYAQKFQGRVTMTSDTSTSTVYMELSSLRSEDTAVYYCARDIYFVPARGGFDPWGQGTLVTVSS |
| **hRBD15_LC** | EIVLTQSPGTLSLSPGERATLSCRASQSIISSYLAWYQQKPGQAPRLLIYGASSRATGIPDRFSGSGSGTDFTLTISRLEPEDFAVYYCQQSGSAPRYTFGQGTKLEIK |
| **hRBD16_HC** | EVQLVESGGALIQPGGSLRLSCAASGFTVSSNYMYWVRQAPGKGLEWVSLIYPGGSTFYADSVKGRFTISRDNSENTLYLQMNSLRAEDTAVYYCARGPYGDSNWGQGTLVTVSS |
| **hRBD16_LC** | DIQMTQSPSSLSASVGDRVTITCQASQDIRNYLNWYQQKPGKAPKLLIYDASNLKTGVPSRFSGSGSGTDFTFTISSLQPEDIATYYCQQYDNLPITFGQGTRLEIK |
| **hS1-1_HC** | QVQLVESGGGVVQPGRSLRLSCAASGFTFSSYAMHWVRQAPGKGLEWVAVISYDGSNKNFADSVKGRFIISRDNSKNTLFLQMNDVRAEDTAVYYCASLSGADFDFWSGYYNWRDSYYYGKDVWGQGTTVTVSS |
| **hS1-1_LC** | DIVMTQSPLSLPVTPGEPASISCRSSQSLLHSNGYNYLDWYLQKPGQSPKLLIYLGSDRASGVPDRFSGSGSGTDFTLKISRVEAEDVGIYYCTQALQTPRTFGQGTKLEIK |
| **hS1-2_HC** | EVQLVESGGGLVKPGGSLRLSCAASGFTFSNAWMSWVRQAPGKGLEWVGRIKSRTDGGTKDYAAPLKGRFTMSRDDSKNTLYLQMNSLKTEDTAVYYCTADPLYDFWNSFYRSFGMDVWGQGTTVTVSS |
| **hS1-2_LC** | DIVMTQSPDSLAVSLGERATINCKSSQSVLYSSDNKNFLAWYQQKPGQPPKLLIYWASTRESGVPDRFSGSGSGTDFTLTISSLQAEDVAVYYCQQYYSTPLTFGGGTKVEIK |
| **hS1-3_HC** | QVQLQESGPGLVKPSQTLSLTCTVSGGSMSSGDFYWSWIRQPPGKGLEWIGYIYHSGSTNYNPSLKSRVIISVDTSKNQFSLKLNSVTAADTAVYYCAREKGDISTGSFNAWFDPWGQGTLVTVSS |
| **hS1-3_LC** | DIVMTQSPDSLAVSLGERATINCKSSQSVLYSSNNRNYLAWYQQKLGQPPKLLIYWASTRESGVPDRFSGSGSGTDFTLTISSLQAEDVAVYYCQQYYNAPRTFGQGTKVEIK |
| **hS1-4_HC** | EVQLVESGGGLVKPGGSLRLSCAASGFTFSSYNMIWVRQAPGKGLEWVSSIIISSSYIYYADSVKGRFTISRDNAKNSLYLQMNSLRVEDTAVYYCAREDVQLGYYYGMDVWGQGTTVTVSS |
| **hS1-4_LC** | QSALTQPASESGSPGQSITISCTGTSSDVGSYNLVSWYQQHPGKAPKLMIYEGSKRPSGVSNRFSGSKSGNTASLTISGLQAEDEADYYCCSYAGSYWLFGGGTKLTVL |

**Supplementary Table 6 Crystallographic data collection and refinement statistics.**

| **Title** | **Fab fragment of mRBD2 complexed with SARS-Cov-2 RBD** |
|---|---|
| PDB code | 8BE1 |
| **Data Collection:** | |
| Beamline | DLS i04 |
| Wavelength [Å] | 0.9795 |
| Resolution range [Å] | 52.05 - 1.98 (2.030 - 1.980) |
| Space group | $P\ 1\ 2_1\ 1$ |
| Cell (a b c) [Å] | 58.47 194.62 62.09 |
| Cell (α β γ) [°] | 90.00 97.16 90.00 |
| Total reflections | 1669839 (122256) |
| Unique reflections | 95377 (4825) |
| Multiplicity | 17.5 (17.2) |
| Completeness (%) | 100.0 (100.0) |
| Mean I/σ (I) | 13.2 (0.3) |
| $R_{merge}$ | 0.109 (9.63) |
| $R_{pim}$ | 0.027 (2.38) |
| $CC_{½}$ | 0.999 (0.33) |
| **Refinement:** | |
| $R_{factor}$ / $R_{free}$ | 0.247 / 0.284 |
| Number of total atoms | 9866 |
| Atoms in ligands | 85 |
| Atoms for waters | 286 |
| Number of polymer residues | 1221 |
| Average/Wilson B-factor [Å$^2$] | 65.0 / 59.2 |
| B-factor for ligands [Å$^2$] | 68.8 |
| B-factor for solvent [Å$^2$] | 60.2 |
| RMS (bonds) [Å] | 0.011 |
| RMS(bond angles) [°] | 0.99 |
| RMS(dihedral angles) [°] | 3.58 |
